# Generative steering to adapt proteins beyond native biological contexts

**DOI:** 10.1101/2025.07.11.664238

**Authors:** Xiaowei Zhang, Wenhui Sophia Lu, Ben Viggiano, Kevin Reynolds, Sophia Burick, Elena Sotillo, Haoyu Dai, Geonhu Lee, Zixuan Liu, Nuoya Yang, Luis S. Mille-Fragoso, Euan Ashley, Bingfei Yu, Crystal L. Mackall, Wing Hung Wong, Xiaojing J. Gao

**Affiliations:** Department of Bioengineering, Stanford University, Stanford, USA; Department of Statistics, Stanford University, Stanford, USA; Department of Biomedical Data Science, Stanford University, Stanford, USA; Department of Computer Science, Stanford University, Stanford, USA; Stanford Center for Cancer Cell Therapy, Stanford Cancer Institute, Stanford University, Stanford, USA; Biophysics Program, Stanford University, Stanford, USA; Weill West Coast Cancer Hub, Stanford, USA; Department of Chemical Engineering, Stanford University, Stanford, USA; Department of Immunology and Immune Therapeutics, Keck School of Medicine, University of Southern California, Los Angeles, USA; Department of Medicine, Stanford University, Stanford, USA; Center for Undiagnosed Diseases, Stanford University, Stanford, USA; Norris Comprehensive Cancer Center, University of Southern California, Los Angeles, USA; Alfred E. Mann Department of Biomedical Engineering, University of Southern California, Los Angeles, USA; Department of Pediatrics, Stanford University, Stanford, USA; Parker Institute for Cancer Immunotherapy, San Francisco, USA; Stanford Bio-X, Stanford University, Stanford, USA

**Author notes:** Corresponding author. (X.J.G.); (W.H.W.); (X.Z.). These authors contributed equally to this work (Xiaowei Zhang, Wenhui Sophia Lu, Ben Viggiano).

## Abstract

Proteins evolved for native biological contexts often fail when deployed in new cellular environments or therapeutic settings. We present ProVADA, a test-time framework that steers a pretrained protein generator toward externally defined properties while preserving functional constraints. Its Mixture-Adaptation Directed Annealing sampler combines coordinated mutations, population selection, and annealing to navigate rugged objective landscapes. Experimentally, ProVADA converted secreted human renin into an active, drug-inhibitable cytosolic protease and extensively deimmunized two murine single-chain antibody fragments to reduce HLA visibility while retaining antigen recognition. ESCAPE-seq measurements confirmed substantial depletion of HLA-I binder peptides across alleles. Deimmunized FMC63 variants retained CAR-T-cell killing and showed fewer binders than a tested humanized comparator. Together, these results demonstrate that broad sequence redesign can adapt functional proteins to non-native biological contexts.

---

Evolution shapes proteins for the biological contexts in which they naturally function, whereas engineering tasks often require them to operate under constraints absent from that history. An engineered protein can therefore retain its intended molecular activity yet fail in deployment because it is incompatible with its new biological context. Protein engineering for therapeutic and synthetic biology applications extends beyond optimizing target binding or catalysis (1–3). Two demanding problems illustrate this challenge: (1) Adapting proteins to non-native subcellular compartments, such as enabling targeted sensing and gene editing to address mitochondrial disorders (4, 5), and relocating secreted proteases to the cytosol to expand synthetic protein circuitry (6), requires preserving protein activity in a drastically distinct environment. (2) Non-human protein domains for biologic, cell, and gene therapies can be recognized as foreign when delivered, since they had no evolutionary history to adapt to the human immune system. In chimeric antigen receptor (CAR) T-cell therapy, where binding domains are often murine in origin, including those clinically approved to target CD19, host responses against CAR-derived sequences can impair cell persistence and limit effective repeat administration (7, 8). Humanization can partially reduce this risk by grafting essential non-human amino acid sequences onto a human framework to reduce immunogenic foreign epitopes (9). However, residual T-cell epitopes particularly at the junctions remain, and the approach does not readily extend to proteins without suitable human framework counterparts (10). Both problems illustrate the challenge where extensive sequence redesign while preserving primary function is required, and evaluation with task-specific assays could also be too costly or ethically prohibitive to run at scale or place inside iterative design loops.

Protein generative models have reshaped protein engineering by learning from natural protein sequences and structures shaped by evolution to generate new designs (11–15). Such models preferentially represent properties under native biological constraints in their training distributions, and provide little enrichment for objectives those constraints did not act on, which are commonly the properties critical for new context adaptation. Existing approaches can incorporate task-specific objectives through post-hoc filtering, model fine-tuning, guided generation, active learning, and combinatorial sequence optimization (16–25). Many of these strategies rely on locally guided changes to improve sequence properties within a protein’s native operation. Context adaptation, by contrast, requires global redesign across fitness barriers while preserving core function. When desirable variants are rare under the generative prior or separated by unfavorable intermediate sequences, direct sampling becomes inefficient and local optimization can stall. Overcoming these challenges requires coordinated global exploration while remaining within biologically plausible sequence space.

Motivated by this need, we introduce Protein Variant ADAptation (ProVADA), a test-time framework for global sequence adaptation toward externally defined properties while preserving structural and functional constraints. ProVADA combines two features that enable efficient exploration beyond local mutational neighborhoods. First, a pretrained generative model proposes broad, coordinated changes across many positions while conditioning on retained residues and available structural information, allowing extensive redesign while favoring biologically plausible sequences. Each complete candidate is then evaluated directly with a task-specific black-box objective, requiring neither gradients nor modification of the underlying model. Second, ProVADA uses the Mixture-Adaptation Directed Annealing (MADA) sampler for global exploration across sequence space. MADA couples context-conditioned mask-and-refill moves with annealed population resampling to steer broad protein sequence redesign through direct evaluation of complete candidates. Under suitable proposal conditions, MADA’s mutation kernel satisfies detailed balance with respect to the annealed target.

We computationally benchmarked ProVADA and experimentally demonstrated it across three protein-design challenges spanning two categories. ProVADA consistently outperforms baseline methods in enriching high-scoring sequences, with its greatest advantage on weakly aligned objectives with rugged fitness landscapes. Experimentally, ProVADA transforms secreted human renin into a catalytically active cytosolic enzyme and extensively redesigns two single-chain variable fragments (scFvs)—including the murine FMC63 domain used in approved CD19 CAR-T—to reduce peptide-HLA-I binding while retaining antigen recognition and cancer-ablation function. Sequence-wide ESCAPE-seq profiling demonstrates substantial depletion of experimentally detected HLA binders (26). Our FMC63 designs require no human framework template, yet all validated variants show fewer HLA binders than a tested humanized comparator, while the lead design also outperforms the parental CAR in tumor-cell killing.

Across all three design cases, ProVADA redesigned approximately one-third of the native sequence. Achieving such extensive, globally distributed redesign while preserving and improving function is virtually infeasible through manual design and remains challenging for conventional computational optimization. Together, these results establish a general strategy for adapting functional proteins to biological constraints that are poorly captured by protein plausibility alone.

## ProVADA enables global sequence adaptation to external objectives

The difficulty of steering a protein generative model depends on two key properties of the design problem: the alignment between the objective and the generative prior, and the local smoothness of the objective under mutation (Fig. 1A). Here, the objective denotes a quantitative evaluation of the target property being optimized, the generative prior denotes the distribution learned by the generative model, and alignment reflects how strongly high-scoring sequences are enriched under that prior. When alignment is strong (Scenario i), the prior assigns substantial probability mass to desirable variants, making direct filtering effective. However, in many adaptation applications, weak alignment limits the effectiveness of filtering because sequences favored by the prior rarely exhibit the target property. When the objective landscape is nevertheless smooth, with accessible paths of successive improvement (Scenario ii), iterative local search can progressively enrich high-scoring variants that are rarely recovered by unguided sampling alone. A particularly challenging regime arises when weak alignment coincides with a rugged objective landscape (Scenario iii). In this setting, desirable sequences remain rare under the prior, and pathways to higher-scoring variants may pass through unfavorable intermediate sequences, leaving local search trapped in suboptimal regions. This regime motivates multi-residue proposals that can bypass such intermediates while remaining within sequence space supported by the generative prior.

**Figure 1:**
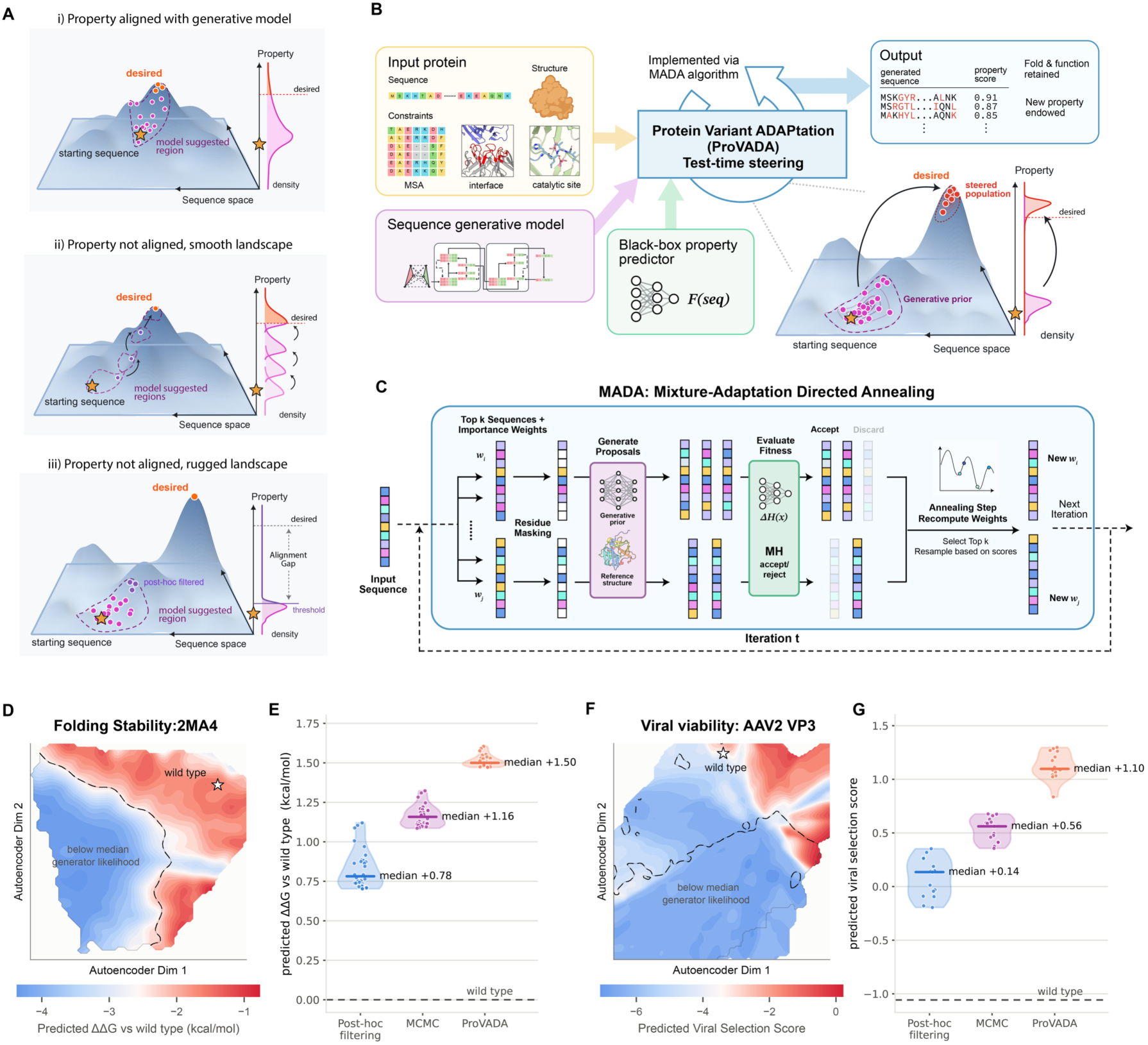
ProVADA steers protein generative models toward desired properties across rugged fitness landscapes. (A) Three schematic relationships between a generative model and a target property. When the two are aligned (top), direct sampling and post-hoc filtering can efficiently recover high-fitness variants. When they are misaligned but the fitness landscape is smooth (middle), iterative methods such as Markov chain Monte Carlo (MCMC) can explore higher-fitness regions. On a rugged landscape (bottom), weak prior alignment can make post-hoc filtering inefficient, while fitness barriers can hinder MCMC exploration. (B) ProVADA combines a sequence generative model with a black-box property predictor and user-specified structural and sequence constraints for test-time steering. (C) Mixture-Adaptation Directed Annealing (MADA), the core sampling algorithm of ProVADA. At each iteration, parent sequences are selected according to their current weights, masked, and refilled by ProteinMPNN conditioned on the retained residues and the reference structure. Proposals are scored and subjected to a temperature-dependent Metropolis acceptance step, retaining the parent on rejection. Population selection and annealed weight updates determine how proposals are allocated in the next iteration. The schematic illustrates greedy top-k selection; campaign-specific resampling choices are given in Methods. (D, F) Predicted fitness landscapes for folding stability of PDB 2MA4 (D) and AAV2 VP3 capsid viability (F), shown in a two-dimensional autoencoder projection. Colors indicate smoothed grid-cell mean predicted scores; stars mark wild-type sequence locations, and dashed contours indicate the median cell-averaged generator log-likelihood. (E, G) Predicted property scores of the top 30 (E) or 15 (G) unique sequences selected from each method’s full sampling history. Panel E shows a single run with a budget of 3,000 generated sequences per method; Panel G uses 1,500 generated sequences per method. Each generated sequence receives one predictor evaluation. Points represent individual sequences; violin plots summarize the score distributions, with horizontal bars marking the medians. Dashed horizontal lines mark wild-type scores. Predicted ΔΔ*G* values are transformed so that higher values indicate greater folding stability. Sampling configurations are detailed in Methods.

Existing approaches connect generative models to external objectives through several distinct paradigms. Fine-tuning and preference optimization modify model parameters to favor target properties and typically require objective-specific training data and model adaptation (22, 23). Guided-generation methods instead bias inference using a property predictor or guidance model (24). Because guidance is applied during decoding, the predictor must score a partially decoded sequence and estimate how the completed protein will perform, where the estimate depends on residues not yet decoded. Errors in these estimates can misdirect exploration under a finite computational budget, particularly early in generation, when most positions remain unresolved. (27–29). For objectives that are more naturally evaluated on complete sequences, evolutionary algorithms and Bayesian optimization provide flexible black-box alternatives (19, 30, 31). ProVADA uses a complementary strategy: broad, multi-residue proposals are evaluated directly as complete candidates under the task-specific black-box objective, while MADA couples population selection and annealing to explore beyond local mutational neighborhoods without requiring objective gradients or retraining the generative model.

ProVADA takes as input a reference protein, optional structural information, user-defined sequence constraints, a conditional mask-and-fill generative model, and an external scoring function (Fig. 1B). Constraints can preserve catalytic, binding, or structurally important residues while allowing broad redesign elsewhere. The scoring function may be any computable sequence-level objective, including a predictor, evaluator, or weighted combination of multiple terms, with optional penalties for sequence divergence or other design requirements. Because the generative model remains fixed and the objective is evaluated externally, ProVADA requires neither gradients through the objective nor retraining of the generative model, and no labels beyond those already used to train a supplied predictor. Thus, the same framework can be redirected to distinct design properties simply by changing the scoring function.

ProVADA defines an annealed target distribution by tilting the generative prior toward sequences with higher objective values, with temperature controlling the balance between the prior and the objective. Under suitable proposal conditions, conditional refilling from the prior allows the Metropolis acceptance rule to depend only on objective differences, without evaluating the full-sequence likelihood. To explore this sequence of annealed targets, the MADA algorithm maintains a population of parent sequences, masks blocks of designable residues, and refills those positions using the generative model conditioned on the retained sequence and, when supplied, the reference structure, enabling multi-residue moves across sequence space (Fig. 1C).

Each completed proposal is evaluated with the external objective and subjected to a temperature-dependent Metropolis acceptance criterion, retaining the parent sequence on rejection. Under suitable proposal conditions, this transition satisfies detailed balance with respect to the corresponding annealed target; the formal assumptions and proofs are provided in the Supplementary Text. Both the temperature and masking fraction decrease over the search, enabling larger exploratory moves early and more localized refinement later. After annealed importance reweighting, MADA uses two-stage stochastic resampling to allocate offspring generation across a smaller pool of parents, with campaign-specific selection schemes detailed in Materials and Methods. Together, generator-conditioned block proposals, direct scoring of complete sequences, population selection, and annealing allow ProVADA to explore broadly across sequence space while respecting the specified sequence constraints and conditioning on available structural information.

## ProVADA efficiently enriches high-scoring sequences across objectives with varying prior alignment and landscape ruggedness

Having established the ProVADA framework, we next asked how effectively it concentrates high-scoring sequences within a fixed computational budget as the relationship between the objective and generative prior changes. To enable a controlled comparison, we focused on inference-time steering methods that use the same fixed generative model, external predictor, and sequence-evaluation budget. Within this setting, post-hoc filtering provides the standard reference for sampling from the prior alone, whereas Markov chain Monte Carlo (MCMC) provides a closely related iterative-search baseline. We compared ProVADA with these methods in two predictive benchmarks. The first used a folding stability predictor trained on large-scale measurements of miniprotein folding stability (32, 33). Because structure-conditioned inverse-folding models are expected to capture sequence features associated with stable folding, this objective represents a setting in which the ProteinMPNN prior should already enrich improved variants, and post-hoc filtering should provide a strong baseline. The second benchmark used an AAV2 capsid viability predictor trained on deep-mutational-scanning data from a capsid hypervariable region (34, 35). Viral viability in this sequence context is a specialized functional property expected to be less directly represented by the generative prior. ProteinMPNN supplied the sequence prior in both benchmarks.

We first visualized the predicted objective landscapes to compare these two settings qualitatively. For each reference protein, we generated sequences containing random substitutions, projected their sequence representations into two dimensions with an autoencoder, and plotted the predicted objective as a third dimension. We then overlaid the contour marking the median ProteinMPNN likelihood to indicate the region more strongly supported by the generative prior. The projected folding-stability landscape appeared relatively smooth, with regions of high predicted stability overlapping substantially with regions of stronger ProteinMPNN preference (Fig. 1D and fig. S1D). By contrast, the projected AAV viability landscape appeared more rugged, with limited overlap between regions of high predicted viability and stronger ProteinMPNN preference (Fig. 1F and fig. S1E). Moreover, across exhaustive point mutations in both design tasks, generator likelihood correlated more strongly with predicted stability scores than with predicted AAV viability scores (fig. S1I-J). These place folding stability near scenario i of Fig. 1A, in which the objective is comparatively well aligned with the prior, and AAV viability near scenario iii, in which weaker alignment is accompanied by a more rugged predicted landscape.

We then evaluated how efficiently each method enriched high-scoring variants under matched budgets of generator invocations and objective evaluations. ProVADA yielded the highest median scores among the top sequences accumulated during each search (Fig. 1E,G and fig. S1G-H). For folding stability, post-hoc filtering alone recovered substantially improved variants, consistent with the prior already enriching sequences with favorable predicted stability. MCMC and ProVADA produced further gains, but their performance was comparatively similar in this aligned, smoother setting. The differences became more pronounced for AAV viability. Post-hoc filtering produced only modest improvement, while both guided methods shifted the sampled population toward higher viral-selection scores. ProVADA achieved a larger gain than MCMC, widening the separation between the two methods relative to the stability benchmark. Thus, ProVADA’s advantage increased in the setting where high-scoring sequences were less enriched by the prior and the predicted objective varied more irregularly under mutation, consistent with the motivation for combining population search with coordinated multi-residue proposals.

It is of note that ProVADA’s advantage was specific to broad redesign: in the AAV2 benchmark, masking only 3 of the 28 designable positions per generation allowed post-hoc filtering to outperform both guided methods, whereas ProVADA performed best when all 28 were masked, the regime the Fig. 1G benchmark tests (fig. S2 and Methods).

## ProVADA adapts human renin for cytosolic activity

After benchmarking ProVADA across distinct predicted fitness landscapes, we next asked whether it could address the biological problem that motivated the framework: adapting an existing protein to function outside its evolved context. We first considered relocating a secreted enzyme to the cytosol. Secreted proteins fold and mature under conditions distinct from the cytosol, which disfavors disulfide-bond formation and lacks N-linked glycosylation. Successful cytosolic adaptation should therefore require coordinated mutations that confer cytosolic compatibility while preserving the catalytic pocket, substrate recognition, and conformational flexibility required for turnover. Such redesign could expand the range of extracellular proteins available for intracellular synthetic circuits, particularly when no native cytosolic protein provides the desired molecular activity.

Human renin provides a stringent and potentially useful test of this challenge. Renin is a secreted human aspartic protease that initiates the renin–angiotensin system by recognizing and cleaving a specific peptide substrate in angiotensinogen. Its substrate specificity and inhibition by the approved small molecule aliskiren make it attractive as a drug-regulated protease for synthetic protein circuits. Engineered, membrane-localized renin has already been used to control secreted cytokine signals in mammalian cells, with aliskiren providing pharmacological regulation (36). A cytosol-compatible renin could extend this capability to the direct regulation of intracellular protein circuits.

However, renin offers little evolutionary signal for cytosolic function. Renin belongs to the pepsin-like A1 family, whose characterized members are generally secreted or localized to endolysosomal or vacuolar compartments (37, 38). In our mammalian reporter system, wild-type renin, with its signal peptide removed, exhibited poor expression and no detectable catalytic activity in the cytosol (Fig. 3G,H). This inactive starting scaffold limits conventional directed evolution campaigns, while the need to measure proteolytic activity in living mammalian systems constrains experimental throughput. Together, these constraints make cytosolic renin a compelling test case for ProVADA, which can prioritize broad computational search into diverse but highly plausible candidates for experimental validation.

To guide cytosolic renin design, we had to train a new classifier (Fig. 2A). Existing classifiers predict the subcellular locations a protein is targeted to, and leverage localization signatures within full-length proteins (39, 40). Instead, our goal was to predict whether the functional portion of a protein is compatible with a specific subcellular location. Accordingly, we curated cytosolic and extracellular proteins from Swiss-Prot, removed signal peptides, prodomains and transmembrane domains, encoded the resulting domains using ESM-2, and trained a classifier on their localization labels (see Methods for details). The resulting predictor outperformed existing methods on both our signal-peptide-depleted held-out dataset and a full-length HPA Secretome dataset (Fig. 2B; Table S1; (39,41–43)). We next examined how the classifier’s cytosolic scores relate to the sequence preferences encoded by ESM-2. Across exhaustive single substitutions in renin relative to wild type (WT), the two scores showed no linear association (Pearson *r* = 0.00; Fig. 2C). To assess whether the predictor captured biologically meaningful sequence features, we visualized predicted cytosolic compatibility change for each substitution relative to the wild-type sequence (Fig. 2D, fig. S3A). Reassuringly, substitutions that increased the cytosolic-compatibility score were enriched at two annotated NXT glycosylation sites and at cysteines forming three native disulfide bonds, consistent with our knowledge of the hallmarks of secreted proteins. Intriguingly, these alterations alone appear quantitatively inadequate to flip the localization probability to more cytosolic than secreted, while there are additional mutations that could move renin further in the cytosolization direction. This is also consistent with our prior unsuccessful cytosolization attempts where we mutated only the disulfide bonds and glycosylation motifs.

**Figure 2:**
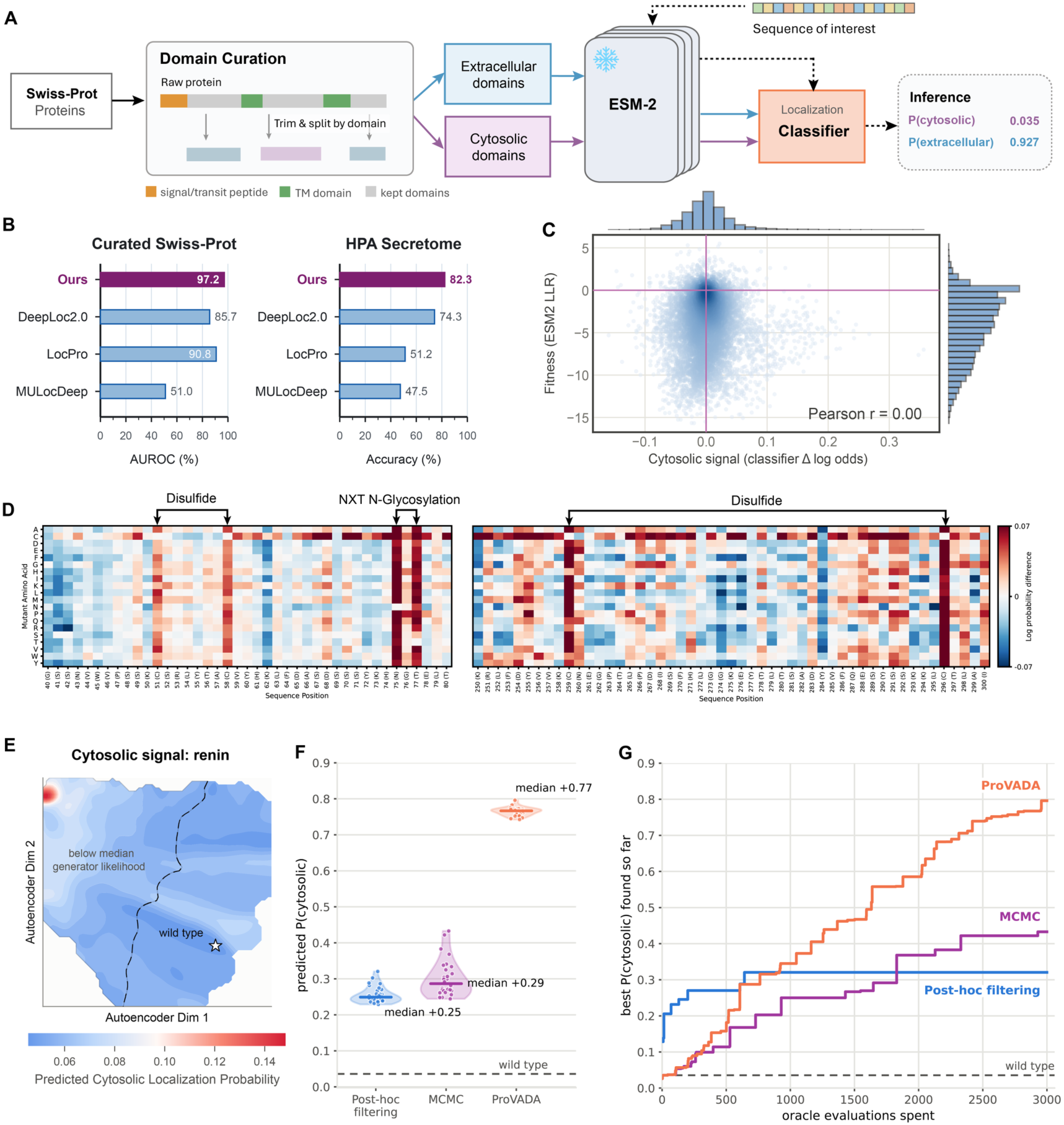
A localization–functionality predictor identifies sequence features associated with subcellular environments and guides ProVADA steering toward cytosolic compatibility. (A) Predictor construction and inference. Extracellular and cytosolic domains curated from Swiss-Prot are encoded using a frozen ESM-2 model to train a localization classifier, which predicts localization probabilities from an input sequence. (B) Predictor performance compared with DeepLoc2.0, LocPro, and MULocDeep on curated Swiss-Prot domains and the Human Protein Atlas (HPA) Secretome dataset, measured by area under the receiver operating characteristic curve (AUROC) and accuracy, respectively. The full evaluation metrics table on curated Swiss-Prot can be found in Table S1. (C) Relationship between classifier-derived cytosolic Δ log odds and ESM-2 LLRs for renin variants relative to WT renin. Purple lines mark wild-type reference values; marginal histograms show the distributions of both scores. (D) Cytosolic log probability ratios to WT for single amino acid substitutions across selected regions on renin. Arrows mark disulfide-linked cysteines and the NXT glycosylation motif. (E) Predicted cytosolic localization probability landscape for renin, visualized as in Fig. 1D C F. (F) Comparison of post-hoc filtering, Markov chain Monte Carlo (MCMC), and ProVADA using the top 30 sequences from each method’s full sampling history, plotted as in Fig. 1E C G. Each method uses 3,000 generated sequences, with one predictor evaluation per sequence. (G) Highest cytosolic localization probability found as a function of cumulative predictor evaluations for the runs shown in F. Sampling configurations are detailed in Methods.

We next visualized the local predicted cytosolic objective landscape around renin. The projection was consistent with a rugged, weakly aligned objective: most sampled variants had low predicted cytosolic probability, whereas a localized region of elevated scores was separated from wild-type renin in the embedding and lay largely below the median ProteinMPNN likelihood contour (Fig. 2E and fig. S1F). Moreover, no linear association was observed between cytosolic signal and generator likelihood on exhaustive point mutations on renin (Pearson *r* = −0.02; fig. S3B). These observations support scenario iii in Fig. 1A, in which weak prior alignment and potential fitness barriers are expected to limit post-hoc filtering and local MCMC search. As expected, under the same matched budget, ProVADA increased the median final cytosolic probability to approximately 0.77, exceeding the classifier’s 0.5 decision threshold; the corresponding medians for MCMC and post-hoc filtering remained below 0.3 (Fig. 2F). ProVADA also reached a substantially higher best-so-far score over the evaluation budget compared with the two baseline methods (Fig. 2G and fig. S3C).

Having established ProVADA’s *in silico* steering performance, we next applied it to design cytosolic renin variants for experimental testing. To favor retention of substrate specificity and catalytic activity, we fixed the most conserved and substrate-contacting residues while requiring mutation of every cysteine to eliminate the native disulfide bonds (Fig. 3A and Methods). An AF3 prediction of monomeric renin served as the structural input (44). ProteinMPNN provided the structure-conditioned generative prior, while the predicted cytosolic probability was regularized by sequence divergence, ProteinMPNN score, and ESM-2 pseudo-perplexity to discourage unnecessary or implausible substitutions (Fig. 3B). ProVADA steers the variants towards increasing cytosolic scores (Fig. 3C and fig. S4A). We filtered the top-scoring sequences along the trajectories based on AF3 prediction metrics and other criteria described in the Methods. We selected 24 variants that preserved the cytosolic score distribution of the broader endpoint population (Fig. 3C), yielding a mean score of 0.78 compared with 0.04 for wild-type renin (fig. S4B). The selected variants are diverse between each other and from wild-type renin (Fig. 3D), while retaining good metrics from structure prediction (fig. S4B). The native disulfide-forming cysteines and NXT glycosylation sites were consistently mutated away (fig. S4C). Altogether, we produced sequence-diverse and structurally plausible variants with substantially improved predicted cytosolic compatibility.

**Figure 3:**
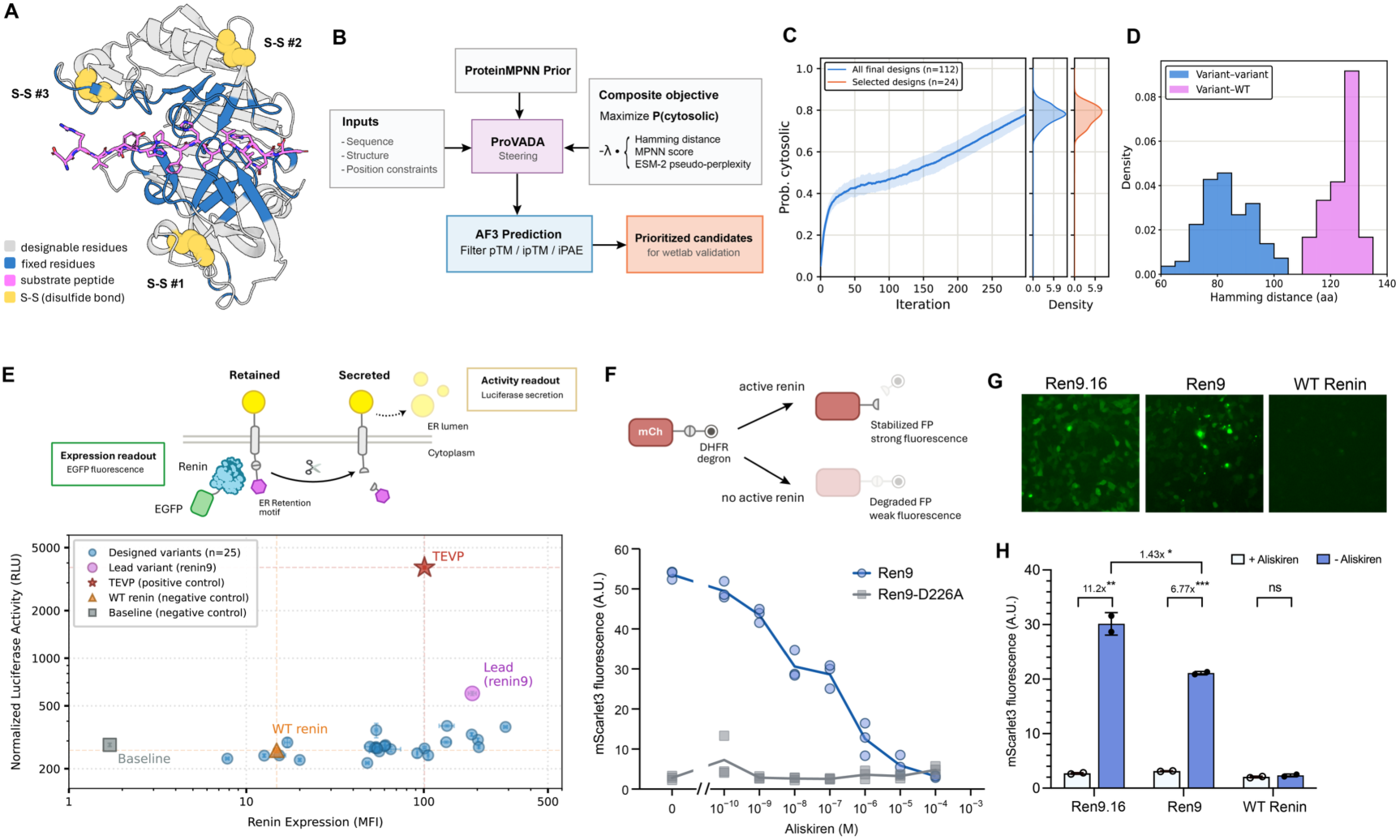
Design and experimental characterization of cytosolically active renin. (A) AlphaFold 3 (AF3)–predicted complex of renin and its substrate peptide (magenta). Residues within 8 Å of the substrate and most conserved residues are fixed during design (blue, see Methods); designable residues are in gray, and disulfide-linked cysteines are shown as yellow spheres. (B) Constrained ProVADA design and candidate-selection workflow. Steering maximizes predicted cytosolic localization probability while penalizing sequence divergence, lower ProteinMPNN likelihood, and higher ESM-2 pseudo-perplexity. Candidates are subsequently filtered using AF3 structural metrics (Methods). (C) Left: mean predicted cytosolic localization probability across 112 design trajectories; shading indicates ± SD. Right: distributions for the best final-iteration sequence from each trajectory (blue; n = 112) and positive candidates selected for experimental validation (orange; n = 24). (D) Hamming distance distributions between selected designs (blue) and between each design and WT renin (magenta). (E) Top: luciferase secretion assay for renin expression and activity characterization. Renin cleavage removes an endoplasmic reticulum (ER) retention motif, enabling luciferase secretion as an activity readout; fluorescence from renin-fused EGFP reports expression. An adjacent tobacco etch virus protease (TEVP) cleavage site enables a positive control. Bottom: expression and reporter activity of designed variants (n = 25, 24 positive designs and 1 negative design that failed the iPAE filter), compared with WT renin (orange), baseline negative controls (no protease, gray) and the TEVP positive control (red). The lead design, Renin9 (Ren9), is highlighted in magenta; dashed lines mark WT and TEVP reference values. Points show means ± SEM from two biological replicates. (F) Top: degron-based fluorescence reporter assay. Renin cleavage separates a red fluorescent protein (FP) from a dihydrofolate reductase (DHFR) degron, stabilizing FP and increasing fluorescence. Bottom: aliskiren dose responsiveness in HEK cells co-transfected with the reporter and Ren9 or its catalytically inactive mutant, Ren9-D226A. Aliskiren was added at transfection. Points show individual measurements from three biological replicates; lines connect means. (G) Fluorescence microscopy of EGFP-fused Ren9.16, Ren9, and WT renin. Ren9.16 is an optimized Ren9 variant with reduced aggregation propensity. Image contrasts were linearly adjusted individually to visualize aggregation; intensities are not directly comparable. (H) Degron-reporter fluorescence for Ren9.16, Ren9, and WT renin with or without 10 μM aliskiren added at transfection. Bars show means ± SEM from two biological replicates; points indicate individual measurements.

To experimentally evaluate expression and catalytic activity, we tested 25 cytosolic renin variants, including the 24 prioritized designs and one additional variant that showed favorable overall structural metrics but did not meet the AF3 ipAE cutoff. Each variant was expressed as a C-terminal EGFP fusion in HEK293 cells and evaluated using a RELEASE reporter, in which proteolytic removal of a cytosol-facing ER-retention motif permits secretion of a luciferase cargo (Fig. 3E) (45). Most variants showed higher EGFP fluorescence than wild-type renin, with several exceeding the Tobacco Etch Virus protease (TEVP, a well-behaving cytosolic protease) positive control under matched plasmid input. Among them, the lead design, Ren9, produced the largest increase in luciferase reporter activity. To further characterize Ren9, we used a fluorescent reporter in which proteolytic removal of a degron stabilizes a red fluorescent protein, enabling single-cell protease activity to be quantified by flow cytometry (Fig. 3F). Aliskiren, a clinically approved transition-state-analogue inhibitor of renin, retains its ability to inhibit Ren9. By contrast, Ren9-D226A, with one catalytic aspartate replaced with alanine, showed only background reporter fluorescence and no response to aliskiren. Together, these results established that ProVADA adapts Ren9 for cytosolic activity while maintaining the catalytic and inhibitory characteristics in and around its active site.

Although Ren9 was catalytically active, this variant and several other designs formed intracellular puncta, consistent with aggregation (fig. S5). We explored whether we could rationally further improve the starting point enabled by ProVADA. We computationally identified and redesigned surface residues that potentially contribute to aggregation (fig. S4D-E). We identified multiple variants that form few puncta and are more active than Ren9 (Fig. 3G–H, fig. S4F-G, fig. S5). The lead design, Ren9.16, showed approximately 1.4-fold higher reporter activity and an 11.2-fold aliskiren-sensitive dynamic range. These variants further yield promising leads for using Renin for synthetic intracellular protease circuits.

In summary, ProVADA adapted renin for the cytosol in a single round of low-throughput screen. It demonstrates that ProVADA can traverse a long distance in the sequence space to retain enzymatic function in a new biological context under a limited computational and experimental budget. The resulting active scaffold also provided a tractable starting point for targeted refinement and could support further improvement through conventional optimization approaches such as directed evolution.

## Preparation for applying ProVADA to deimmunize proteins via minimizing HLA-I visibility

Another especially compelling protein adaptation problem is to minimize the immunogenicity of therapeutic proteins while preserving their functionality. Immunogenicity is the capacity of a therapeutic protein or its processed peptides to elicit antibody or T-cell responses that can neutralize the therapy, alter its pharmacology, or eliminate cells expressing it (46), which has been a long-standing problem in the field of protein therapeutics. Although humanization can substantially reduce this risk in antibody-derived sequences, it’s far from fully resolving the issue. Conventional humanization via CDR grafting retains nonhuman CDR residues and creates novel CDR–framework junctions, leaving potential T-cell epitopes within the humanized sequence (9, 10). Grafting can also compromise folding stability or target affinity, which further requires compensatory mutations that may introduce additional non-self peptides. Most importantly, scaffold-based humanization requires a suitable human framework and therefore does not readily extend to nonhuman enzymes, gene editors, or de novo proteins, yet those modalities are becoming increasingly relevant to cell and gene therapies. These limitations motivate more generalizable protein deimmunization strategies.

A complementary strategy is T-cell-epitope depletion, in which sequence substitutions are introduced to reduce the presentation of protein-derived peptides on human leukocyte antigen (HLA) molecules (47). T-cell-epitope depletion can address both HLA-I-and HLA-II-mediated responses by targeting distinct presentation pathways; here, we focused on HLA-I presentation, which can elicit CD8^+^ T-cell recognition and lead to clearance of engineered cells expressing therapeutic proteins. Formation of a stable peptide–HLA-I complex is a necessary upstream step in conventional CD8^+^ T-cell recognition, because it provides the cognate ligand for the T-cell receptor. Depleting HLA-I-binding epitopes can thus minimize the pool of peptides available for presentation, which would preclude subsequent CD8^+^ T-cell recognition regardless of the T-cell-receptor repertoire.

However, sequence-wide epitope depletion is a demanding multi-objective design problem. Extensive HLA polymorphism requires candidate peptides to be evaluated across multiple HLA allotypes. Because potential epitopes are distributed throughout a protein, broad deimmunization requires coordinated substitutions at multiple positions while avoiding neo-epitope formation and preserving native fold and function. Experimental validation presents another bottleneck: conventional peptide–HLA binding and T-cell assays can evaluate only a limited number of candidate epitopes, while functional validation must often be tailored to each therapeutic protein in relevant context. Although previous studies have successfully deimmunized individual proteins, these efforts have generally required target-specific design and extensive experimental screening, limiting their scalability (48–50). More recent approaches combine protein models with HLA predictors to identify functional variants with reduced predicted immunogenicity (51). However, the post-hoc workflow remains limited because candidates are sampled without direct HLA guidance. Guided decoding has also been explored, but remains limited to *in silico* validation (52). ProVADA addresses these limitations through test-time steering: it preserves the protein prior during proposal generation, evaluates complete sequences using a sequence-wide, multi-allelic HLA objective, and concentrates an experimentally tractable candidate set.

We started by constructing an immunogenicity score as the ProVADA steering oracle: a protein-level measure of predicted HLA-I visibility for peptides absent from the canonical human proteome. (Fig. 4A). We focused initially on HLA-I because (1) HLA-I predictors are comparatively mature and extensively benchmarked (53, 54), (2) therapeutic proteins expressed in cells directly enter this presentation pathway, and (3) high-throughput assays are available to massively characterize peptide-HLA-I display (26, 55), although the framework itself is not restricted to HLA-I and could similarly incorporate an HLA-II objective. For each protein, we enumerated all overlapping 9-mer peptides and excluded exact matches to the canonical human proteome, which are unlikely to elicit T-cell responses because of central and peripheral tolerance. The remaining peptides were evaluated against the selected HLA allotypes using the pan-allelic predictor NetMHCpan-4.2 (54), and their predicted binding profiles were aggregated into a protein-level score used for steering. Details are described in the Methods.

**Figure 4:**
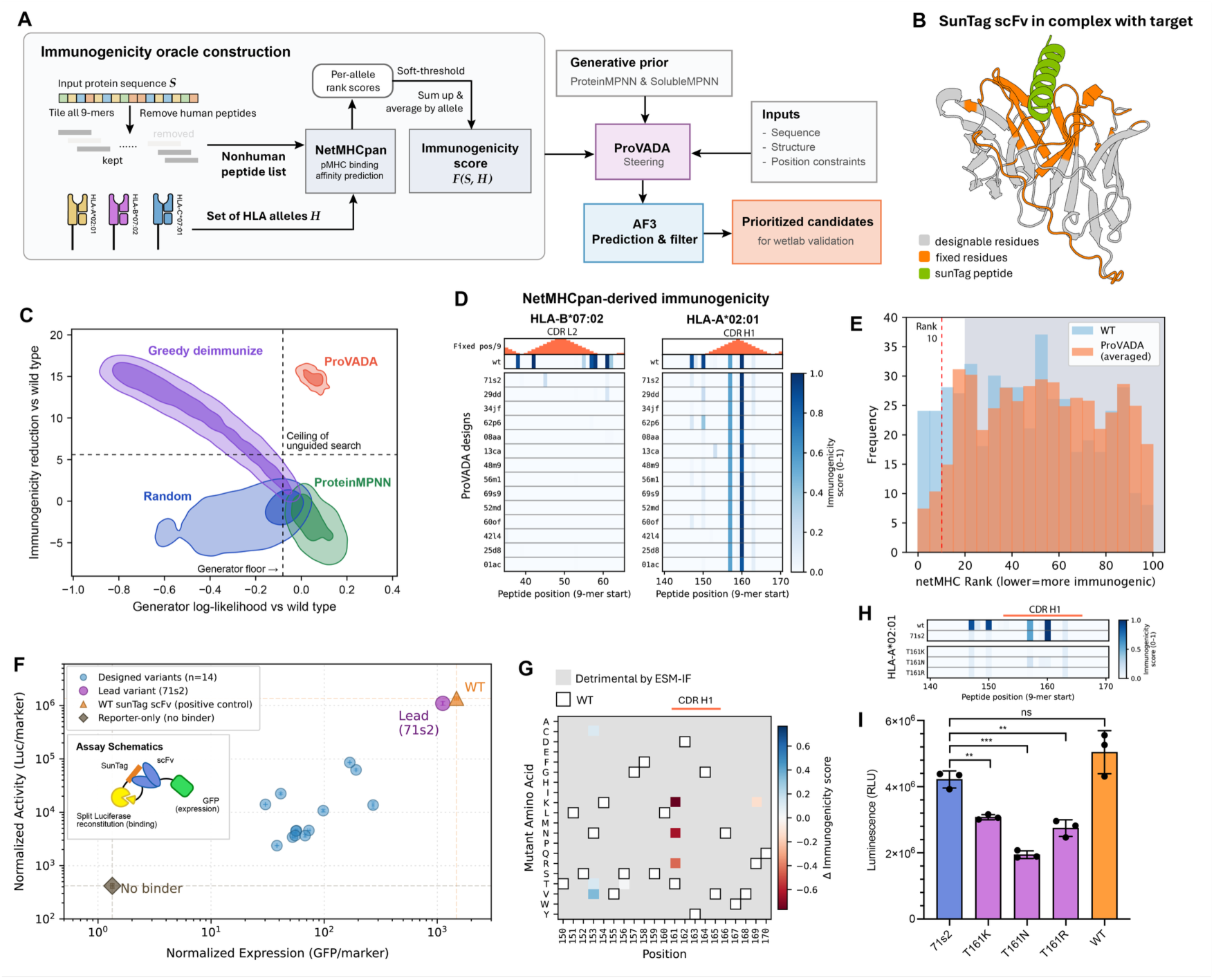
ProVADA reduces predicted immunogenicity of the SunTag scFv while retaining expression and antigen binding. (A) Construction of the protein immunogenicity oracle and constrained deimmunization workflow. (B) AF3-predicted complex of the SunTag scFv and SunTag V4 peptide (green). Residues on scFv fixed during design, including complementarity-determining regions (CDRs), vernier zones, and the linker, are colored in orange; designable residues are in gray. (C) Density plots of generator log-likelihood and predicted protein immunogenicity reduction, both relative to the starting scFv, for random mutagenesis (blue), greedy deimmunization (purple), ProteinMPNN sampling (green), and ProVADA (orange). Filled contours enclose 50% and 90% of each population. Dashed lines mark the generator likelihood floor and the greatest reduction achieved by the baseline methods above that floor. See Methods sections for details. (D) NetMHCpan-derived immunogenicity scores for consecutive 9-mer peptides on regions from the parental WT scFv and selected designs, shown for HLA-B*07:02 (left) and HLA-A*02:01 (right). Peptides are indexed by their starting position; upper bars indicate the number of fixed residues within each peptide, corresponding CDR regions are annotated on top. (E) Distributions of NetMHCpan binding-affinity (BA) percentile ranks for the starting scFv and selected ProVADA designs, with design frequencies averaged across variants. Lower ranks indicate stronger predicted binding; the dashed line marks a rank of 10; the shaded region marks ranks above 20, which contribute minimally to the calculated protein immunogenicity score. (F) Expression and antigen-binding reporter activity of prioritized designs (n = 14), WT SunTag scFv (orange), and no-binder control condition (gray). The lead design, 71s2, is highlighted in magenta. Inset: assay schematic. The scFv carries N-terminal SmBiT and C-terminal GFP; The reporter contains a nuclear-localized SunTag peptide fused to LgBiT, and an mCherry (not shown) for normalization of transfection efficiency and visualization of nuclear localization. GFP fluorescence reports expression, and split-luciferase complementation reports target binding. Points show means ± SEM from two biological replicates. (G) Predicted changes in immunogenicity score for point mutations around CDR H1 of 71s2. CDR H1 positions are marked by the orange line above the heatmap. Gray cells indicate substitutions excluded by ESM-IF filtering; outlined white cells mark the parental 71s2 residues. (H) Peptide-level immunogenicity scores around CDR H1 for HLA-A*02:01, comparing WT SunTag scFv, 71s2, and its T161K, T161N, and T161R variants, displayed as in D. The orange line above the heatmap marks peptides that contain at least one residue in CDR H1. (I) Antigen-binding reporter activity of 71s2 (blue), WT SunTag scFv (orange), and the three T161 variants (purple). Bars show means ± SEM from three biological replicates; points indicate individual measurements.

## Deimmunization of the SunTag scFv

As an initial proof of concept, we applied ProVADA to deimmunize a mouse-derived scFv intrabody used in the SunTag system (56). We fixed all complementarity-determining regions (CDRs), CDR-contacting Vernier zones, and the linker between the V*_H_* and V*_L_* domains (Fig. 4B). We then used the immunogenicity oracle described above to steer ProVADA against two common HLA alleles, HLA-A*02:01 and HLA-B*07:02. We used this design campaign to first benchmark ProVADA against baseline approaches (Fig. 4C). Random mutagenesis produced mostly implausible sequences with little predicted immunogenicity reduction, whereas unsteered ProteinMPNN sampling maintained high plausibility but slightly increased predicted immunogenicity. Greedily combining immunogenicity-reducing point mutations achieved substantial deimmunization at the expense of sequence plausibility, exiting the plausible region after only a ∼25% reduction in immunogenicity score. In contrast, ProVADA pushed beyond the plausibility-immunogenicity trade-off, maintaining high generator likelihood while matching the immunogenicity reduction of the best greedy designs at a similar mutation budget. Thus, ProVADA reduces predicted immunogenicity without compromising sequence plausibility *in silico*.

From the ProVADA-generated sequences, we selected 14 diverse designs that passed structural filters based on AF3 predictions (Methods). Before testing them experimentally, we performed closer computational assessments of their features. On average, these designs exhibited an approximately fourfold reduction in predicted immunogenicity score and fewer peptides with low NetMHCpan percentile ranks (fig. S7E and Fig. 4E). In contrast, the lowest-immunogenicity sequence obtained by sampling the ProteinMPNN prior with a matched computational budget showed only a modest reduction (fig. S7C-D). Sequence-wide heatmaps further revealed fewer regions of strong predicted HLA binding than in the parental sequence (Fig. 4D and fig. S7A-B). Peptides containing few or no fixed residues were more readily deimmunized, whereas those overlapping extensively with fixed regions were more likely retained because of the imposed constraints. The selected designs are also distinct from the parental sequence while being structurally favorable (fig. S7E). Together, ProVADA introduces extensive sequence-level changes that reduce predicted epitope burden while maintaining favorable predicted structural metrics.

We next evaluated the expression and target-binding activity of the redesigned scFvs. Expression was measured using a C-terminal EGFP fusion, while binding was assessed through split-luciferase reconstitution and nuclear colocalization with a cognate reporter. Most variants fell along a diagonal spanning the background and parental-scFv controls (Fig. 4F). The lead design, 71s2, exhibited expression and activity comparable to those of the parental scFv (Fig. 4I and fig. S8). To further address the remaining predicted epitopes overlapping with the fixed CDR regions, we adapted a workflow similar to that of Wolfsberg et al. (51). We used a structure-informed protein language model to identify tolerated point mutations throughout the scFv, including the CDRs (17). At CDR-H1 residue 161 of the lead design 71s2, several substitutions were predicted to be tolerated while strongly reducing immunogenicity (Fig. 4G). In particular, T161K eliminated two predicted immunogenic peptides (Fig. 4H). Experimental testing revealed all selected substitutions to retain target binding (Fig. 4I). Thus, targeted follow-up mutagenesis can address residual epitopes within regions fixed during ProVADA design, extending toward more comprehensive deimmunization.

Finally, we applied ESCAPE-seq to systematically assess the impact of our strategy on peptide–HLA-I binding. ESCAPE-seq is a massively parallel assay based on the cell-surface display of peptide–MHC-I single-chain trimers (SCTs) (26). HLA-I, β2-microglobulin (B2M), and cognate peptide are expressed as a single construct (Fig. 5A). Peptides that bind strongly to HLA stabilize the complex and promote its trafficking to the cell surface, whereas weakly binding peptides result in destabilized complexes with reduced surface display. ESCAPE-seq is scaled through combinatorial pooled screening (Fig. 5B). We constructed a pooled library containing all unique nonhuman 9-mer peptides from all relevant proteins in this manuscript pre-and post-deimmunization, including peptides from the subsequent FMC63 campaign, and assayed them against the two HLA alleles targeted during design. Details are provided in Methods. ESCAPE-seq enrichment-scores (E-scores) were highly reproducible across independent biological replicates (Fig. 5C). Although the control pHLA measurements were largely consistent with the reported E-scores, the library-wide E-score distribution was more bimodal (Fig. 5C)(26). This difference likely reflects the lower proportion of binders in our library, which required broader medium and high FACS bin gates to recover sufficient cells. Based on the trough position of the bimodal distribution, we used an E-score threshold of 5 to identify peptide-HLA binders. For visualization and analysis, all human peptides excluded from ESCAPE-seq measurements are denoted with an E-score of 0, indicating minimal immunogenicity risk.

**Figure 5:**
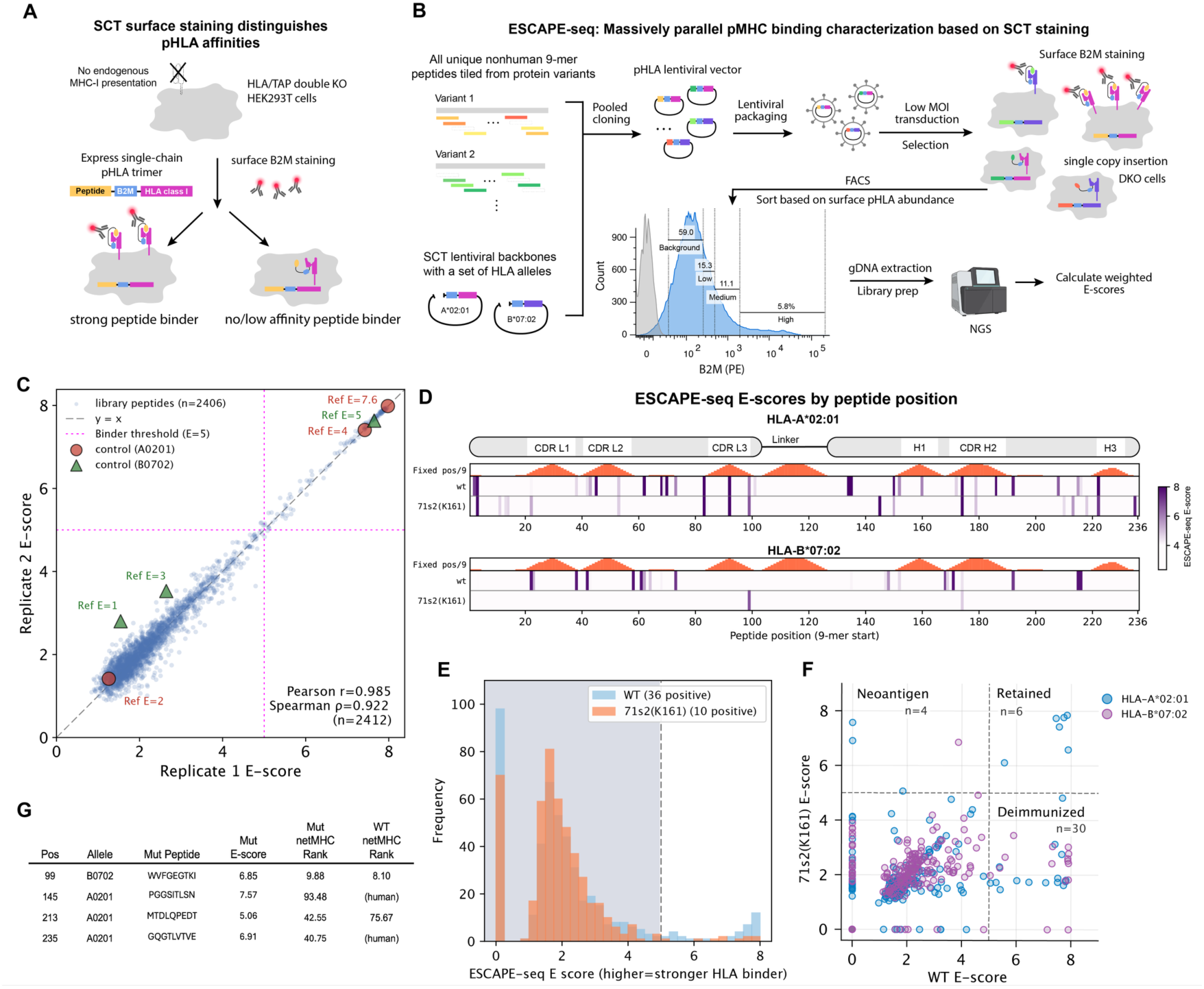
ESCAPE-seq validation of reduced peptide-HLA binding after deimmunization. (A) Single-chain trimer (SCT) surface-display assay. SCTs comprising a peptide, β2-microglobulin (B2M), and an HLA class I molecule are expressed in HLA/TAP-deficient HEK293T cells. Stronger pHLA binding supports greater SCT surface display, measured by anti-B2M staining. (B) ESCAPE-seq workflow, adapted from Shi et al. with modifications described in Methods (26). Unique nonhuman 9-mer peptides from selected SunTag and FMC63 scFv designs, corresponding WT sequences, and a humanized scFv control were paired with HLA-A*02:01 and HLA-B*07:02 in a pooled SCT library. Following low-multiplicity lentiviral transduction and selection, cells were stained for surface B2M and sorted into four fluorescence bins. Next-generation sequencing of each bin was used to calculate weighted E-scores. The representative sorting histogram shows stained cells (blue), an unstained control (gray), bin boundaries and frequencies. (C) Reproducibility of E-scores across two biological replicates. Reference pHLA controls from Shi et al. are highlighted for HLA-A*02:01 (red) and HLA-B*07:02 (green), with published E-scores annotated. Magenta dashed lines mark the binder-detection threshold (E = 5); the gray dashed line indicates agreement between replicates. Pearson and Spearman correlations are reported for all 2,412 measurements, including controls. (D) Peptide-level E-scores across WT SunTag scFv and the deimmunized variant 71s2-T161K for both HLA alleles. Schematics above the heatmaps mark the CDR-overlapping peptides on the scFv; a peptide is annotated on a specific CDR if it contains at least one residue on that CDR region. Peptides are indexed by their 9-mer starting position; upper bars indicate the number of fixed residues within each peptide. In all ESCAPE-seq related figures including this one and below, human peptides that were excluded from the assay are assigned with E-score = 0. (E) E-score distributions for WT SunTag scFv and 71s2-T161K, with 36 and 10 pHLA pairs above the detection threshold, respectively. The dashed line marks the E-score threshold of 5; scores below this threshold are shaded. (F) Comparison of WT and variant E-scores for pHLA pairs matched by peptide position and allele. Dashed lines at E = 5 separate pairs below threshold in both sequences (negative, lower left), pairs shifted below threshold after design (deimmunized, lower right; n = 30), retained binders (retained, upper right; n = 6), and newly detected binders (potential neoantigen, upper left; n = 4). HLA-A*02:01 and HLA-B*07:02 are shown in blue and purple, respectively. (G) Variant peptide sequences, measured E-scores, and NetMHCpan-BA percentile ranks for the four newly detected binders along with ranks for their corresponding WT peptides.

For the SunTag scFv, the tiled peptide heatmaps of E-scores revealed a marked reduction in experimentally detected HLA binders for both alleles, including nearly complete depletion for HLA-B*07:02 (Fig. 5D-E). The stronger performance for HLA-B*07:02 was consistent with its lower false-omission rate compared to HLA-A*02:01 in our dataset (fig. S10A). We also observed allele-specific discrepancies between NetMHCpan predictions and ESCAPE-seq measurements (fig. S10B–C). The predictor underestimated binding for HLA-A*02:01 across most of the NetMHCpan rank-score range, which is more likely to produce false negatives that remain unaddressed during design and places an upper bound on deimmunization performance.

Next, we quantified deimmunization effects by comparing ESCAPE-seq E-scores for matched peptides from the parental scFv and the redesigned variant. Of the 36 binders detected in the parental scFv, 30 fell below the detection threshold after redesign. Four newly detected binders emerged as potential neoantigens (Fig. 5F). Three of these peptides are NetMHCpan false-negatives, while the remaining was predicted to have weaker binding after redesign, whereas the measured E-score increased (Fig. 5G). All told, ESCAPE-seq confirmed a substantial reduction in experimentally detected peptide-HLA binding while revealing newly created binders that the oracle failed to capture. These findings support the effectiveness of the ProVADA deimmunization pipeline and show that its performance is partly limited, as expected, by the accuracy of peptide–HLA binding predictions.

## Deimmunization of the FMC63 CD16-CAR scFv

Having established the feasibility of our deimmunization pipeline with the SunTag scFv, we next applied it to the clinically relevant murine FMC63 scFv used in approved anti-CD19 CAR T-cell therapies. Anti-CAR T-cell immune responses have been associated with reduced persistence of infused CAR-T cells and therapeutic failure, particularly following repeat administration (7, 8). Immune responses have also been reported in patients receiving CAR-T cells incorporating humanized scFvs (57), underscoring the inadequacy of conventional humanization. While surface display of the scFv makes it accessible to antibody responses, the CAR is endogenously synthesized and naturally presented on MHC-I, and MHC-I–restricted CD8 responses have shown to impair CAR-T cell potency (8). Reducing the MHC-I visibility of CAR antigen-binding domains therefore possesses direct therapeutic relevance.

The FMC63 scFv was deimmunized using the same strategy described above. ProVADA was steered against HLA-A*02:01 and HLA-B*07:02, with key residues fixed (Fig. 6A). We selected 18 designs after structural filtering. These candidates exhibited an approximately fourfold reduction in predicted immunogenicity and favorable structural metrics relative to the parental FMC63 (fig. S11B), and the representative tiled peptide heatmap on NetMHCpan-derived immunogenicity also showed a clear reduction in predicted binder peptides (fig. S11A). On average, approximately 28% of their residues differed from the parental sequence, indicating extensive sequence-wide redesign (fig. S11B).

**Figure 6:**
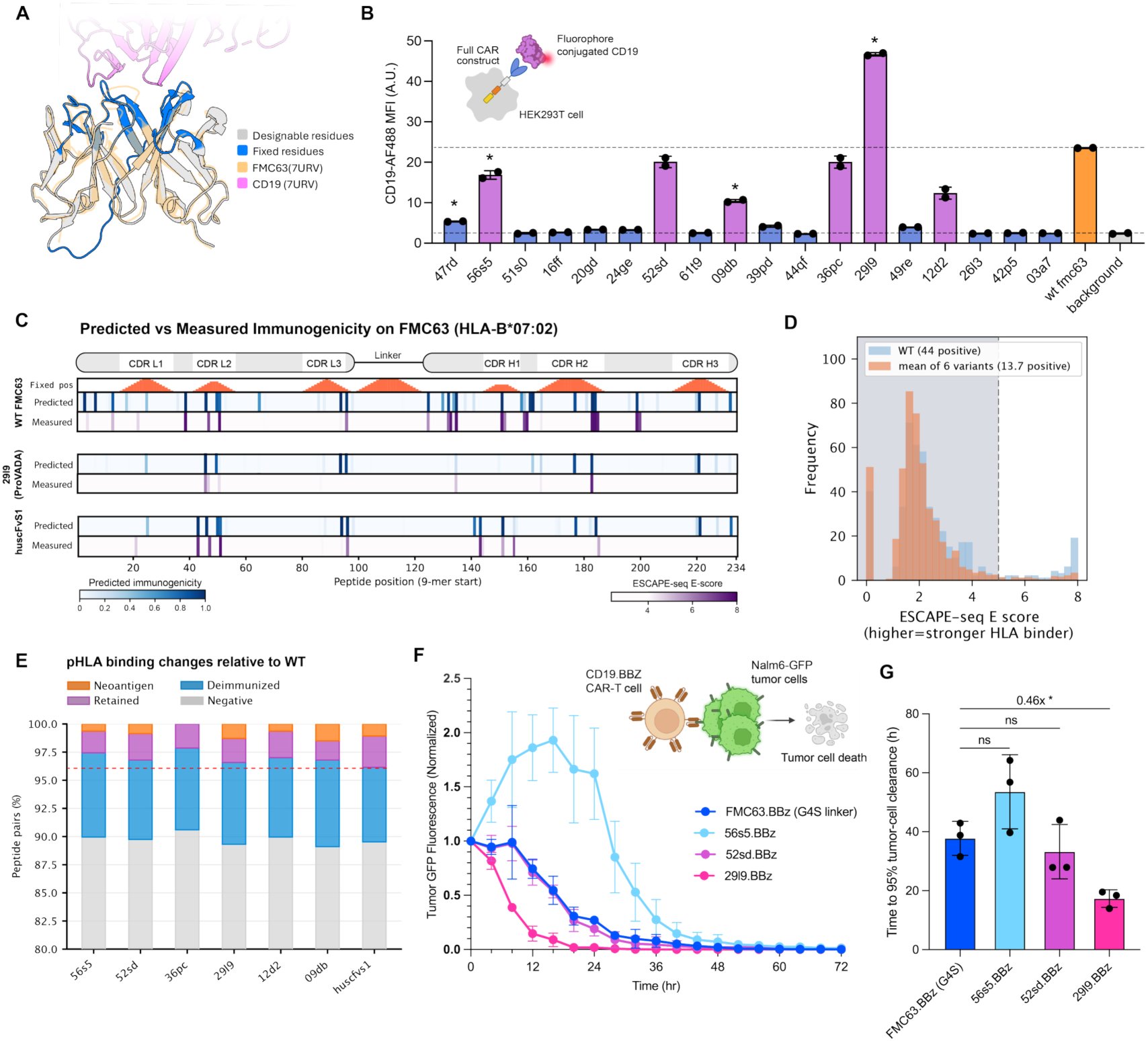
Deimmunization of the FMC63 CAR scFv while retaining target killing capacity. (A) AF3-predicted FMC63 structure aligned to the experimentally determined FMC63–CD19 complex (PDB 7URV). The experimental FMC63 structure is colored in orange and CD19 epitope is colored in magenta; fixed CDR residues and designable residues in the predicted structure are in blue and gray, respectively. (B) CD19-AF 488 staining of HEK293 cells expressing 18 selected FMC63 designs on a Gen2 CAR construct. Inset: assay schematic. WT FMC63 CAR and background controls are in orange and gray, respectively; designed variants are in blue, with six variants selected for further analysis highlighted in magenta. Dashed lines mark WT and background staining levels. Bars show means ± SEM from two biological replicates; points indicate individual measurements. Asterisks indicate significant differences from background after Bonferroni correction. (C) Heatmap showing predicted (blue) vs ESCAPE-seq measured (purple) peptide immunogenicity on HLA-B*07:02 for overlapping 9-mers on WT FMC63 (top), ProVADA designed lead variant 29l9 (middle), and a humanized FMC63 scFv huscFvS1 (bottom). Schematics above the heatmaps mark the CDR-overlapping peptides on the scFv; a peptide is annotated on a specific CDR if it contains at least one residue on that CDR region. Peptides are indexed by their 9-mer starting position; upper bars indicate the number of fixed residues within each peptide. The comprehensive heatmaps of all selected variants and the heatmaps for HLA-A*02:01 can be found in fig. S11. (D) ESCAPE-seq E-score distributions for WT FMC63 (blue), compared with the mean bin frequencies across the six selected designs (orange). Distributions combine HLA-A*02:01 and HLA-B*07:02 measurements; dashed lines mark the binder-detection threshold (E = 5). Regions with score below this threshold are shaded. (E) Fractions of pHLA pairs in each binding category, matched between each variant and WT by peptide position and allele and classified using the E = 5 threshold as in Fig. 5F. Categories comprise pairs below threshold in both sequences (negative, gray), shifted below threshold after design (deimmunized, blue), retained binders (retained, purple), and newly detected binders (neoantigen, orange). Both HLA alleles are included. Horizontal red dashed lines indicate the fraction of nonimmunogenic peptides in huscFvS1. The allele-specific plots can be found in fig. S12C. (F) Killing curve of CD19-expressing Nalm6-GFP tumor cells by primary T cells expressing CARs containing FMC63 or the indicated deimmunized scFv variants. Tumor-cell GFP fluorescence was normalized to its value at 0 h. Points and error bars show mean ± SD (*n* = 3 biological replicates). (G) Time to 95% tumor-cell clearance (*T*_95_), defined as the time at which normalized tumor GFP fluorescence reached 0.05. *T*_95_ was calculated separately for each replicate by linear interpolation between the two measurements bracketing this threshold. Bars show mean ± SD, and points indicate individual replicates. Fold change denotes mean *T*_95_ relative to FMC63. Each variant was compared with FMC63 using a two-sided unpaired Welch’s *t*-test with Bonferroni correction for three comparisons.

Each selected variant was then cloned into a second-generation CAR containing a 4-1BB costimulatory domain and a CD3ζ signaling domain. The resulting CAR constructs were expressed in HEK293 cells, and CD19 recognition was evaluated by staining with Alexa Fluor 488 (AF488)-conjugated recombinant CD19. Six of the 18 designs showed CD19-staining more than twofold above baseline (Fig. 6B). The lead design, 29l9, exhibited nearly twofold higher staining than the parental FMC63. These results show that extensive ProVADA-mediated redesign did not abolish antigen recognition and identified variants with strong CD19-binding signals in the CAR context.

We next asked whether the six CD19-binding variants also showed reduced peptide–HLA-I binding. Using ESCAPE-seq, we tested their peptides alongside those from parental FMC63 and the FMC63-derived humanized scFv huscFvS1 (58). The tiled peptide heatmaps on E-scores showed fewer detected binders in every ProVADA variant than in parental FMC63 (fig. S11C). Similar to the SunTag scFv, the reduction was greater for HLA-B*07:02 than for HLA-A*02:01. Humanization also reduced binding, but a substantial number of binders remained near CDR–framework junctions. Across both alleles, 44 parental FMC63 peptide–HLA-I pairs exceeded the detection threshold, compared with a mean of 13.7 on ProVADA variants and 18 on huscFvS1 (Fig. 6D and fig. S11D). Similar to SunTag scFv, we then quantified the deimmunization effects on the variants compared to the parental FMC63. Both ProVADA redesigns and humanization removed a large fraction of parental binders, with differing individual profiles, and all validated ProVADA redesigns had fewer remaining HLA binders than the humanized comparator (Fig. 6E). The allele-specific comparisons again showed a greater reduction for HLA-B*07:02 than for HLA-A*02:01 (fig. S12A-C).

Finally, we evaluated the potency of CAR-T cells bearing the deimmunized scFvs in primary T cells. All of the tested variants eliminated CD19-expressing Nalm6 tumor cells within 72 hours (Fig. 6F). Notably, as shown by Fig. 6G, variant 29l9 mediated approximately twofold faster tumor-cell killing than the wild-type FMC63 CAR, consistent with its higher overall expression and CD19 staining (fig. S12D and Fig. 6B).

Together, these results demonstrate that ProVADA can generate extensively redesigned scFv variants with substantially reduced predicted and experimentally measured peptide HLA-I binding while retaining functionality. In the FMC63 campaign, ProVADA achieved greater overall epitope depletion than the tested humanized comparator. Unlike conventional scaffold-based humanization, ProVADA does not require a predefined human framework and is therefore applicable to non-antibody-based therapeutic proteins. This generality offers a route toward reducing the MHC-I visibility of therapeutic modalities for which conventional humanization is not available.

## Discussion

ProVADA offers a general framework for adapting proteins to specific biological contexts by steering a generative model at test time. We demonstrated this strategy across two distinct engineering challenges: restoring cytosolic activity for the secreted enzyme renin, and depleting HLA-I epitopes in scFvs while retaining antigen binding. In both campaigns, a single design-and-test cycle yielded functional variants from a small screen, which provides leads for downstream refinement as needed. Together, these results show that context-aware redesign can create functional variants with extensive sequence alterations.

The renin campaign addressed an especially difficult starting condition: wild-type renin shows no detectable cytosolic activity. Despite replacing a substantial fraction of the native sequence, ProVADA successfully generated active cytosolic variants in its first attempt. The initial designs were prone to aggregation, likely reflecting a mismatch between the ProteinMPNN prior and the intracellular environment. ProteinMPNN is trained on solved structures, a dataset heavily biased toward stable, purified proteins rather than cytosolic solubility (13). To tackle this, explicitly penalizing aggregation propensity during steering should help in future runs. Even without that penalty in the initial search, follow-up post-hoc redesign reduced visible aggregation while boosting expression and activity in the Ren9.16 variant. Although Ren9.16 is less active than the TEVP positive control, deriving a functional baseline from a dead scaffold provides a key foundation for downstream optimization, such as laboratory evolution, a path supported by recent work using AI-redesigned proteins as starting points (59).

While ProteinMPNN has been extensively applied to optimize enzymes, earlier efforts did not tackle the broader problem of compartment adaptation. For instance, Sumida et al. (16) optimized TEVP starting from an enzyme that already had measurable baseline activity, whereas our renin campaign began with zero detectable cytosolic function. Our functional renin variants also tolerated higher mutation rates than the functional threshold reported in that study. Besides, their optimized TEVP variant was known to suffer from a reduced intracellular activity when expressed in mammalian cells, a context drastically distinct from their in vitro screening assays. This mismatch underscores why performance gains in standard screening assays cannot be assumed to translate into deployment context, and why generative models must be guided by context-specific objectives. Another related precedent comes from ProteinMPNN-assisted antibody-to-intrabody conversions (60). However, in antibody scaffolds, improved stability can be a dominant route to intracellular function. Enzymes present a stricter trade-off: over-stabilizing an active state can interfere with the conformational flexibility required for catalysis, and antibody-specific design rules do not directly solve enzyme translocation. Tailoring enzymes to new compartments thus requires active, context-aware guidance rather than broad stabilization alone.

The deimmunization campaigns show that ProVADA can handle entirely different biological objectives. Across both SunTag and FMC63 campaigns, broad sequence redesign substantially reduced detected binding to two common HLA-I alleles without compromising antigen recognition. To our knowledge, this is the first systematic HLA-I binder depletion strategy for therapeutic proteins that closes the design loop with combined experimental validation of both target function and epitope depletion. In FMC63, ProVADA designs achieved greater binder depletion than their humanization counterpart without relying on a human framework scaffold. Our immunogenicity objective currently evaluates peptide-HLA binding, an upstream step in T-cell recognition-induced immune response. Because generative redesign already eliminates the vast majority of predicted epitopes, downstream validation can pair T-cell reactivity assays, such as ELISpot, with targeted post-hoc edits at any residual reactive sites. Expanding coverage to additional HLA-I alleles is straightforward, as is incorporating HLA-II predictors like NetMHCIIpan or MARIA for therapeutics where CD4+ responses are critical (61, 62). We prioritized HLA-I experimentally due to the feasibility of high-throughput binding screens, whereas thorough HLA-II and *in vivo* validation require longer development cycles. Although demonstrated on scFvs to allow direct comparison with humanized controls, ProVADA is scaffold-agnostic and equally applicable to nonhuman enzymes, de novo designed proteins, or CRISPR-associated proteins, for which conventional humanization is not applicable.

A key practical requirement for test-time steering is fast sequence evaluation during sampling. Because the search scores thousands of candidate sequences per trajectory, lightweight sequence-level predictors are essential. Incorporating full structural evaluation at every step could incur prohibitive computational cost and thus limit sampling depth. ProVADA instead reserves structural evaluation for candidate filtering or local refinement, keeping the primary search fast and scalable. As sequence-based models for traits like solubility, aggregation, and developability improve, they can be plugged directly into the generative loop to enable multi-objective steering.

Beyond the two applications studied here, context-aware steering offers a general strategy for complex protein engineering tasks. Tailoring proteins to organelle-specific contexts, such as mitochondria or lysosomes, could unlock new therapeutic targets in those specialized cellular compartments that are traditionally difficult to access. For deimmunization, pairing improved HLA predictors with faster, higher-throughput HLA presentation assays, especially for HLA-II, could establish standardized workflows for safer therapeutic development. More broadly, therapeutic translation of proteins requires balancing activity with developability metrics like solubility, stability and yield under manufacturing conditions; incorporating developability metrics directly into generative search could capture these liabilities much earlier in discovery. Finally, ProVADA is also capable of closing the design-build-test-learn loop by feeding experimental results back to the predictor via active learning. In this way, context-aware generation links computational search, functional screening, and iterative optimization across diverse engineering challenges.

## Acknowledgments

We thank Prof. Brian Trippe, Prof. Michael Z. Lin and Prof. Possu Huang for helpful discussions and insightful remarks. We thank Connor C. Call for providing help and suggestions on lentiviral transduction. We thank Xinyu Ashlee Feng, Hank Jones and Jingjia Liu for help on NGS Library preparation. We thank the Stanford Shared FACS Facility (RRID:SCR_017788) for flow cytometry instrumentation and sorting support.

## Funding

X.Z. was supported by the Stanford Interdisciplinary Graduate Fellowship affiliated with ChEM-H. W.S.L. was supported by the Stanford Data Science Fellowship and NIH grant GM 134483. W.S.L. and X.Z. were both supported by Stanford HAI-Google Cloud Credits Grant.B.V. was supported by a predoctoral training grant from the National Library of Medicine and by the MAC3 Data Science and Precision Medicine Fellowship. S.B. was supported by the Stanford Molecular Biophysics Training Program (T32 GM136568). Z.L. was supported by Sarafan ChEM-H Chemistry/Biology Interface Program and National Institute of General Medical Sciences of the National Institutes of Health (NIH 5T32GM139790). L.S.M.-F. discloses support for the research of this work from Stanford Bio-X Fellowship, the Stanford Graduate Fellowship, the Stanford University Sarafan ChEM-H Chemistry-Biology Interface training program and the Stanford Interdisciplinary Graduate Fellowship. X.J.G. was supported by the Stanford Bio-X Interdisciplinary Initiatives seed grant program (R12-8, to X.J.G.). W.H.W. was supported by NSF DMS 2310788. K.R., E.S., and C.L.M. were supported by NIH Grants: R35CA283888 (C.L.M.), 2P30CA124435-16, the Virginia and D.K. Ludwig Fund for Cancer Research (C.L.M.). C.L.M. is a member of the Parker Institute for Cancer Immunotherapy, which supports the Stanford University Cancer Immunotherapy Program and is a Weill West Coast Cancer Hub Investigator. B.Y. was supported by the V Foundation, Margaret E. Early Medical Research Trust, the Pew Charitable Trusts and the Alexander and Margaret Stewart Trust.

## Author contributions

X.Z., W.S.L., and X.J.G. conceived the study. W.S.L. and W.H.W. developed its mathematical foundation. W.S.L., B.V., and X.Z. implemented the computational pipeline. B.V., X.Z., and W.S.L. performed the computational benchmarks. X.Z. designed, filtered, and experimentally validated the renin variants with assistance from S.B., H.D., and L.S.M.-F. X.Z. designed, filtered, and experimentally validated the SunTag and FMC63 deimmunization variants with assistance from G.L. X.Z. performed the ESCAPE-seq experiments with assistance from B.Y., N.Y. and Z.L.; B.Y. and X.Z. analyzed the sequencing data. K.R., E.S., and X.Z. evaluated deimmunized CAR function in primary T cells. X.Z., W.S.L., B.V., and X.J.G. wrote the manuscript with input from all authors. X.J.G. supervised the overall study. W.H.W. supervised the theoretical work. E.A. supervised B.V.’s computational work. C.L.M. supervised the work of K.R. and E.S. on the primary T-cell CAR experiments.

## Competing interests

E.S. is coinventor on multiple patents related to CAR T cell therapies, holds equity and received royalties from Lyell Immunopharma and consults for Galaria and Lepton Pharmaceuticals. E.A. is a founder of Personalis, Deepcell, Svexa, Saturnus Bio, and Swift Bio; a founder and advisor to Candela and Parameter Health; an advisor to Pacific Biosciences; a non-executive director of AstraZeneca and Dexcom; a holder of publicly traded stock in Personalis, Pacific Biosciences, and AstraZeneca; and a recipient of collaborative support in kind from Illumina, Pacific Biosciences, Oxford Nanopore, and Cellsonics, as well as Cache. C.L.M. is a coinventor on multiple patents related to CAR T cell therapies, received royalties from Lyell Immunopharma and CARGO Therapeutics, holds equity in Link Cell Therapies and Ensoma, which are developing CAR-based therapies and consults for Link, Immatics, Ensoma, Astra-Zeneca, Moderna, Kite Pharma, Nektar and Red Tree Capital. X.J.G. is a co-founder and serves on the scientific advisory board of Radar Tx. All other authors declare no competing interests.

## Data, code and materials availability

The source code for ProVADA can be found on GitHub (https://github.com/SUwonglab/ProVADA). All key plasmids used in this study can be found in Data S5 with link to the Benchling plasmid maps, and the physical plasmid DNA can be provided upon request. All prioritized protein variant sequences that are experimentally validated can be found through Data S1-S3. Peptide level ESCAPE-seq E-scores are provided in Data S4.

## Materials and Methods

### Computational Methods

To benchmark ProVADA against other sequence-optimization strategies, we assembled three *in silico* design tasks: improving protein folding stability of 2MA4, improving AAV2 capsid viability, and altering the predicted subcellular localization of renin. Each task paired a starting protein sequence with a sequence-based predictive fitness oracle, which allowed the optimization metrics to be compared using the same predictive measure. For each task, we evaluated candidate sequences using the corresponding oracle and tracked how predicted fitness changed over the course of optimization. The construction of the fitness oracles is described below, followed by the procedures used to visualize their fitness landscapes and compare optimization methods. These computational benchmarks are distinct from the subsequent protein redesign campaigns.

#### Construction of fitness oracles

##### (1) Folding stability oracle

The changes in folding stability were predicted using a structure-conditioned regression model developed by Nisonoff et al. (33), which was trained on the experimental stability measurements of Tsuboyama et al. (32). The model takes a fixed backbone together with a candidate sequence and returns a single predicted stability value. For the optimization task, fitness was defined as the predicted stability change relative to the wild type sequence, or ΔΔ*G* = Δ*G*(variant) − Δ*G*(WT), in kcal/mol, with the wild type at zero by construction. Note that in this work, we adopt the sign convention of Tsuboyama et al. (32), in which positive ΔΔ*G* denotes stabilization, which is opposite to the convention used in Xiong et al. (24).

We focused on 2MA4, a 70-residue mini-protein corresponding to cluster 107 of the Tsuboyama dataset, with its provided AlphaFold2-predicted wild-type structure used for backbone conditioning (63). Nisonoff et al. reported a mean Spearman correlation of 0.80 across held-out wild-type clusters when developing the stability regression model, and then released the checkpoint we used after training on the full dataset (33). We independently evaluated the accuracy of their released model by scoring the 3,117 measured variants of cluster 107 and comparing the predicted ΔΔ*G* against the values reported by Tsuboyama et al. The model was able to achieve a Spearman *ρ* = 0.926, Pearson *r* = 0.915 and RMSE 0.622 kcal/mol.

##### (2) AAV viability oracle

To construct the AAV2 capsid viability oracle, we used the Proto framework (64) to train an ESM-C 600M based predictor (65) on capsid viability data from the Bryant et al. dataset (34). The dataset experimentally characterizes 289,805 variants of a 28-residue hypervariable segment of the AAV2 as viral selection score, which reports the log enrichment of each variant in the assembled viral pool relative to the input plasmid library. We framed viability prediction as regression on the VP3 sequences with the inserted variable region. Prior to training we removed duplicated variants and discarded all variants that had premature stop codons. We then partitioned the remaining variants into an 80/10/10 train/validation/test split under a fixed random seed, holding the test split out of both training and model selection. Our predictor model employed a seven-layer perceptron with a hidden layer width of 2048 and ReLU activations. We conducted full model training for five epochs, and attained a Spearman correlation of *ρ* = 0.920 (Pearson *r* = 0.959, RMSE 0.96, MAE 0.70) on the held-out test set, closely matching its validation performance (*ρ* = 0.921).

##### (3) Localization-functionality oracle

We curated vertebrate cytosolic and extracellular proteins from UniProt Swiss-Prot (66), restricting to entries with experimental evidence and excluding those localized to lysosomes, endosomes, peroxisomes, or mitochondria. To isolate the intrinsic sequence determinants of subcellular fitness, we removed signal peptides and transmembrane domains that correlate with localization but do not reflect functional fitness (39). We retained only mature peptide domains, discarding annotated propeptide segments. When multiple domains are annotated, we kept only individual domains of 50–1000 residues; for transmembrane proteins, we extracted extracellular and cytosolic domains based on their transmembrane geometry within the same size range. To balance protein-family representation, we clustered the sequences at 30% identity using MMSeq2 (67), retaining one representative per cluster (reducing the dataset to approximately 40% of its original size). Finally, we stratified and split the data into training, validation, and test sets in a 70:20:10 ratio. We embedded each protein sequence using the ESM2-650M-UR50D model due to its high downstream-task performance and balanced compute–accuracy trade-off (68). Unless otherwise specified, all references to *ESM-2* in this work denote this model. For each sequence, we extracted per-residue representations from the final (33rd) transformer layer and mean-pooled them into a 1280-dimensional vector. We then trained a multi-label logistic regression classifier on these embeddings, with separate outputs for cytosolic and extracellular localization.

##### (4) HLA-I immunogenicity oracle

We constructed the protein-level HLA-I immunogenicity oracle to quantify the predicted burden of HLA-I-binding peptides on a given protein, restricted to 9-mers with no exact match in the canonical human proteome. It is based on NetMHCpan-4.2 peptide–HLA binding predictions and serves as a proxy for HLA-I immunogenicity (54). NetMHCpan-4.2 is a pan-specific neural-network model that predicts peptide interactions with MHC-I molecules of known sequence. The model provides two relevant outputs: a binding-affinity (BA) prediction trained on quantitative peptide–MHC binding measurements and an eluted-ligand (EL) likelihood trained on MHC-associated peptides identified by mass spectrometry (54, 69, 70). The BA output primarily models intrinsic peptide–MHC binding, whereas the EL output also captures sequence features associated with antigen processing and presentation (69, 70). Although the EL output is more directly aligned with predicting naturally presented peptides, we selected the BA output because it correlated more strongly with ESCAPE-seq E-scores in our benchmarking analysis (fig. S6A), consistent with ESCAPE-seq directly measuring peptide-dependent pHLA stabilization while bypassing much of the endogenous antigen-processing pathway (26). Moreover, because our objective was deimmunization, we adopted a conservative criterion in which peptides predicted to bind HLA-I were treated as potential liabilities even if they might not be efficiently generated or presented in their native sequence context. This approach may classify some non-presented peptides with potential risks but reduces the likelihood of retaining potential HLA-I binders during sequence design, consistent with conservative binding-based approaches to computational protein deimmunization (71).

To evaluate HLA-I immunogenicity on a protein sequence, we first tile it into all consecutive overlapping 9-mer peptides. Peptides that exactly matched a canonical human 9-mer were excluded from subsequent scoring. The human 9-mer reference set was generated from canonical protein sequences in the UniProt human reference proteome (proteome ID UP000005640, UniProt release 2026_01) (66): All human proteins were tiled into overlapping 9-mers, and are collected to yield a reference set of 11,728,617 unique human peptides.

Each remaining non-human peptide on the protein of interest was evaluated against the selected set of HLA alleles using the NetMHCpan-4.2 BA percentile rank, denoted %*Rank*_BA_, yielding *n_P_* * *n_H_* peptide–HLA predictions, where *n_P_* and *n_H_* are the numbers of peptides and HLA alleles, respectively. For peptide *i* and HLA allele ℎ, the predicted rank *r_i_*_ℎ_ was converted to a peptide–HLA immunogenicity score using a decreasing logistic function:

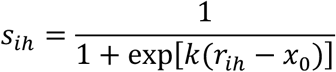

NetMHCpan by default classifies MHC-I peptides with percentile ranks below 0.5 and 2.0 as strong and weak binders, respectively (69). However, as deimmunization prioritizes on capturing any potential HLA binders to avoid false-negatives, we used a more permissive detection range based on the false-omission rate analysis shown in fig. S6B. We set *x*_0_ = 10 and *k* = 0.5, such that lower percentile ranks received higher scores. Under this parameterization, a pHLA pair with %*Rank*_BA_ = 10 receives a score of 0.5, whereas pairs with ranks over 20 contribute minimally to immunogenicity (fig. S6C). The final protein-level MHC-I immunogenicity score was calculated by summing the pHLA-immunogenicity scores across all peptide–HLA pairs and dividing by the number of evaluated HLA alleles:

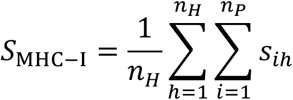

#### Fitness landscape visualization

To provide visualizations of the high-dimensional fitness landscape of each optimization task, we trained a multilayer perceptron-based autoencoder for each target to reconstruct one-hot-encoded sequences through a two-dimensional bottleneck via per-position cross-entropy loss. We then divided the resulting two-dimensional latent space into a grid of cells and calculated the mean predicted fitness of a range of sampled sequences. To produce the resulting surface, we smoothed fitness values with a Gaussian kernel with a width of *σ* = 2 cells. Cells were only displayed in the surface if they met the minimum effective support of five sequences. Sequence sets used in the visualization were constructed separately for each target, utilizing provided sequences as well as sequences generated through random substitutions to create landscapes with at least 70% of cells to be filled in. For the various targets, the final landscapes included 40,331, 36,091, and 4,854,434 variants for 2MA4, Renin, and AAV2, respectively.

To indicate the generator’s preference within the landscape, we also calculated the mean generator log-likelihood by grid cell and included a contour at the median value among displayed cells, as shown in the 2D fitness plot landscapes in Fig. 1 (D,F) and Fig. 2(E).

#### In silico steering with ProVADA and benchmarking

Across the three *in silico* optimization tasks, we evaluated ProVADA against two reference methods: post-hoc filtering, which selects the highest-scoring sequences from an unguided sample, and Markov chain Monte Carlo (MCMC), which uses the same proposal mechanism and Metropolis–Hastings acceptance rule as ProVADA but does not resample the population between iterations. The MCMC baseline used parallel chains, each maintaining its own state across iterations without population resampling. The highest-scoring unique sequences were selected from the combined sampling history of all chains. For all tasks, we used ProteinMPNN as a structure-conditioned generator (72), conditioned on the wild-type backbone. Backbones were obtained from the AlphaFold2 model accompanying the Tsuboyama dataset for 2MA4 (32), from AlphaFold3 for renin (44), and from Protenix for AAV2 VP3 (73). In each setting, candidate sequences were generated by masking a subset of the designable positions in a current sequence and filling them with ProteinMPNN. Each method received an identical budget of *N* generations over *T* iterations, returning the best sequences encountered across the entire trajectory that generated *N* × *T* total evaluated sequences. Folding stability and localization were run at 100 × 30 = 3,000 oracle evaluations. Due to generation being more expensive with the larger VP3 protein, AAV viability was run at 50 × 30 = 1,500 evaluations. In all settings, the top 1% unique sequences (by score) were collected and reported as the final returned distribution. Cysteine was omitted from the proposal alphabet throughout to avoid introducing potential unintended disulfide bonds. To ensure a fair comparison, the masking schedules used to select designable positions in MCMC and ProVADA were identical. We set annealing temperatures separately for each objective to account for differences in fitness scale across tasks. The masking fraction for post-hoc filtering was adjusted to match the final mutation fraction achieved in MCMC and ProVADA for each benchmarking scenario. Per-task sampling parameters and temperature schedules are given in Table S2.

In the AAV2 benchmark, the relative performance of these methods depended strongly on how much of the AAV2 hypervariable region was supplied as existing sequence. In an additional comparison, we masked only three of the 28 designable positions per ProteinMPNN generation, preserving most of the starting sequence. Under this setting, post-hoc filtering substantially outperformed both MCMC and ProVADA on predicted viral-selection score (fig. S2). By contrast, when all 28 positions were initially masked, ProVADA performed best (Fig. 1G and fig. S2). Thus, for AAV2 designs requiring only a few substitutions, sampling near an existing sequence and filtering by predicted fitness can outperform iterative guided search. If the design is restricted to three predefined positions, all 20^3^amino acid combinations can also be evaluated directly, leaving little need for an iterative search method. The AAV benchmark in Fig. 1G tests broader redesign of the region, the setting for which ProVADA is primarily intended.

#### Protein variant redesign campaigns for experimental validation

Candidate sequences were generated using ProteinMPNN with predictor-guided steering. This section describes the sequence-generation, scoring, and sampling configurations used in ProVADA; downstream AlphaFold 3 (AF3)-based structural filtering is described separately below. Campaign-specific sampling parameters, numbers of trajectories, and score weights for each design task are summarized in Table S3. For random masking, the stage-specific masking fraction determines the number of designable positions masked at each iteration. Conditional on this number, positions are selected uniformly without replacement from the fixed set of designable positions, making the mask-selection distribution independent of the current sequence. Some campaigns additionally use greedy parent selection as a computationally efficient adaptation that concentrates the finite evaluation budget on high-scoring candidates. For greedy parent selection, the post-acceptance population, including parents retained after rejected proposals, is ranked by the campaign-specific composite fitness score. The top *K* = max{1, ⌈*ρN*⌉} entries are retained, where *N* denotes the population size and *ρ* the specified retention fraction.

Following the cytosolic renin design campaign, ProVADA was updated to improve computational efficiency, introducing campaign-specific implementation differences that make certain parameter values not directly comparable across campaigns. Specifically, we replaced the original PyTorch implementation of ProteinMPNN (72) with the JAX implementation provided by ColabDesign (74) to improve sampling efficiency. The original PyTorch-based ProVADA pipeline stored mean log-likelihood scores, whereas the ColabDesign-based implementation reports mean negative-log-likelihood scores; in both conventions, values closer to zero indicate higher likelihood. In the updated implementation, task-specific property scores were min–max scaled using fixed reference bounds, clipped to [0,1], and oriented so that higher values indicate better performance. Composite fitness was then calculated as a weighted mean of changes in these normalized scores relative to the starting sequence.

**Cytosolic renin design..** The mature human renin sequence (UniProtKB P00797; chain identifier PRO_0000026082) was used as the wild-type (WT) input. The input renin structure was generated using AF3 through AlphaFold Server, with the renin amino acid sequence supplied as the sole input (44). To preserve substrate recognition and catalytic activity, 92 substrate interfacing positions and 6 conserved positions were fixed during sequence redesign. Interface positions were defined as renin residues containing at least one heavy atom within 8 Å of the N-terminal angiotensinogen segment DRVYIHPFHLVIHNE in the predicted human renin-angiotensinogen complex structure. The two catalytic residues are included in this set of interface residues. A set of non-cysteine positions that were invariant across sequences in the ConSurf-generated MSA were also fixed (75). Cysteines were excluded from the conservation-based constraints, allowing them to be redesigned because their native disulfide bonds are unlikely to form in the reducing cytosolic environment. The complete set of fixed positions is listed below:

Interface positions: 15, 16, 17, 18, 19, 20, 35, 36, 37, 38, 39, 40, 41, 42, 43, 45, 61, 79, 80, 81, 82, 83, 84, 85, 86, 87, 88, 115, 116, 117, 118, 119, 120, 121, 122, 123, 124, 126, 127, 128, 129, 131, 132, 134, 135, 136, 137, 138, 200, 202, 224, 225, 226, 227, 228, 229, 230, 231, 232, 233, 234, 251, 252, 253, 254, 286, 288, 289, 293, 294, 295, 297, 298, 299, 300, 301, 302, 303, 304, 305, 306, 307, 308, 309, 310, 312, 313, 314, 315, 316, 317, 318

Conserved positions: 27, 29, 69, 129, 316, 319

The cytosolic renin redesign campaign was done with an earlier version of ProVADA with PyTorch ProteinMPNN implementations, and the objective was to maximize cytosolic probability with a mismatch penalty, an MPNN score penalty, and an ESM-2-based One Fell Swoop (OFS) pseudo-perplexity penalty (76) that penalizes sequences with lower likelihood scores. Detailed sampling parameters can be found in Table S3.

**Deimmunized SunTag scFv design.** The amino acid sequence of the SunTag scFv (scFv-GCN4), including the variable heavy (V_H_) and variable light (V_L_) domains and the linker in between, were obtained from Addgene plasmid pHR-scFv-GCN4-sfGFP-GB1-dWPRE (plasmid no. 60907). The optimized v4 SunTag peptide, EELLSKNYHLENEVARLKK, was used as the cognate ligand (56). A monomeric structure of the scFv was generated with AF3 through AlphaFold Server (44) and used for sequence generation in ProVADA. Separately, an AF3 predicted scFv–SunTag peptide complex structure was used to define sequence constraints. Complementarity-determining regions (CDRs) and vernier-zone residues were identified using the pipeline described by Galindo et al. and the Martin antibody-numbering scheme (60, 77). These positions together with the GS linker positions were held fixed during sequence generation. Because cysteine was omitted during sequence generation, native cysteine positions were excluded from the fixed set, allowing their redesign to avoid disulfide bond formation. The complete set of fixed positions is listed below:

Fixed positions on SunTag scFv: 4, 25, 29, 30, 31, 32, 33, 34, 35, 36, 37, 38, 48, 49, 50, 51, 52, 53, 54, 55, 56, 57, 58, 72, 92, 93, 94, 95, 96, 97, 98, 99, 100, 112, 113, 114, 115, 116, 117, 118, 119, 120, 121, 122, 123, 124, 125,126, 127, 128, 129, 130, 131, 159, 160, 161, 162, 163, 164, 165, 166, 167, 177, 178, 179, 180, 181, 182, 183, 184, 185, 186, 187, 188, 189, 191, 202, 228, 229, 230, 231, 232, 234

**Deimmunized FMC63 CD19-CAR scFv design.** The DNA and amino acid sequences of the FMC63 scFv were obtained as a gift from Prof. Rogelio Hernandez-Lopez. This scFv uses 3xGGGGS as the linker between V_H_ and V_L_ domains. Monomeric AF3-predicted FMC63 structure using AlphaFold Server was used as the structural input for sequence generation.

Only the CDR and the linker regions are fixed on FMC63. CDR regions were determined according to those reported for huCTL019, a humanized FMC63-derived scFv, in U.S. Patent Application Publication US 2014/0271635 A1 (78). The complete set of fixed positions for deimmunized FMC63 is listed below:

Fixed positions on FMC63 scFv: 24, 25, 26, 27, 28, 29, 30, 31, 32, 33, 34, 50,51, 52, 53, 54, 55, 56, 89, 90, 91, 92, 93, 94, 95, 96, 97, 108, 109, 110, 111, 112, 113, 114, 115, 116, 117, 118, 119, 120, 121, 122, 153, 154, 155, 156, 157, 172, 173, 174, 175, 176, 177, 178, 179, 180, 181, 182, 183, 184, 185, 186, 187, 220, 221, 222, 223, 224, 225, 226, 227, 228, 229, 230, 231

The deimmunization redesign campaigns were done with the updated ProVADA codebase with JAX ProteinMPNN implementations. The objective was to minimize immunogenicity score, with an MPNN score penalty to penalize sequences with lower likelihood scores. Detailed sampling parameters can be found in Table S3.

#### Sequence downsample, structural prediction and filtering

Following ProVADA sampling, candidate sequences were downsampled based on their fitness scores. Around 100 prioritized sequences from each design campaign were subsequently evaluated by structure prediction using AF3 with campaign-specific configurations. Sequences are filtered based on a set of structural metrics derived from structure predictions to acquire a final set of variants for experimental validation. Campaign-specific downsample and structure prediction settings are described as follows:

**Cytosolic renin.** From the final iteration of each design trajectory, the highest-fitness sequence was selected, yielding 112 candidates for structural predictions. Each renin variant was predicted in complex with the substrate peptide, supplied as a separate chain. Because MSA-free AF3 predictions did not reliably recover the WT renin structure in preliminary testing, variant complexes were predicted using AlphaFold Server with its default automatically generated MSA. For each candidate, the top-ranked AF3 prediction was used for structural analysis. The predicted renin chain was aligned to the AF3-predicted WT renin structure used for sequence generation, and the C*α* RMSD was calculated over all aligned renin residues. The mean interchain minimum PAE was calculated by averaging the off-diagonal elements of the AF3 chain_pair_pae_min matrix. Candidates were selected for experimental validation if they satisfied all of the following criteria: mean pLDDT of renin chain > 88, mean interchain minimum PAE < 3 Å, C*α* RMSD < 1.5 Å, and ProteinMPNN score > −1.15. The sequences selected for experimental validation, together with the associated computational and experimental metrics, can be found in Data S1.

**Deimmunized SunTag scFv.** For all generated sequences, at most 4 sequences are selected per trajectory, including: (1) sequence with the lowest immunogenicity score across all iterations; (2) sequence with the highest composite fitness score across all iterations; (3) sequence with the highest composite fitness score in the last iteration; (4) Among sequences whose unnormalized immunogenicity score is within 0.1 above the minimum, the one with the highest MPNN likelihood based on MPNN score. Sequences that contain potential cryptic NLS are filtered out, identified by pattern KKxK, KxKK, or KKK. A total of 84 sequences were selected for AF3 structural prediction. The structures of SunTag scFvs are predicted with the SunTag v4 peptide as a separate chain, and with a local implementation of AF3 without MSA. Each sequence is predicted with 5 random seeds, and each structure prediction undergoes 3 recycling steps. The top-rank structure for each protein was selected for extraction of structural metrics. We also calculated interaction prediction Score from Aligned Errors (ipSAE) based on AF3 predicted structural complexes (79). The C*α* RMSD was calculated based on the input scFv structure for sequence generation. Candidates were selected if they satisfied all of the following criteria: mean pLDDT of the scFv chain > 85, interface pTM (ipTM) > 0.75, and ipSAE > 0.6. Finally, to validate a diverse set of sequences, for all the sequences that pass the structural filter, we only selected one sequence per trajectory that has the highest composite fitness score.

During the initial ProVADA design, linker positions 127–131 were not held fixed. The computational metrics reported for these designs were calculated using the originally generated sequences. Before synthesis for experimental validation, we manually restored the original SGGGS linker sequence at these positions, resulting in a slight increase in immunogenicity scores. All experimentally validated sequences contained the restored linker. Data S2 lists both the originally prioritized sequences and the linker-restored sequences used for experimental validation, together with their associated metrics.

**Deimmunized FMC63 scFv.** Initial sequence downsampling was performed as described for the SunTag scFv. A total of 65 sequences were selected for AF3 structure prediction in MSA-free mode. Monomeric scFv variant structures were predicted because AF3 did not reliably predict CD19 even with an MSA, and a reliable interface with WT FMC63 was not obtained even when CD19 was structure-templated using PDB 7URV. AF3 structure prediction was performed as described above. C*α* RMSD was calculated relative to the input scFv structure used for sequence generation. Candidates with pTM > 0.8 were retained. Similar to the SunTag scFv, only the highest-fitness sequence from each trajectory was selected in the final candidate set for experimental validation. The sequences selected for experimental validation together with the associated metrics can be found in Data S3.

#### Fluorescence microscopy assessment of aggregation propensity

Aggregation propensity was visually evaluated from Green-channel fluorescence microscopy images of transiently transfected renin variants 48 hours post transfection. Ren1-Ren25 were transfected at 50 ng plasmid per well in 24 well plates; Ren9 variants, TEVP and WT renin controls were transfected in a separate experiment at 25 ng plasmid per well. TEVP was used as the non-aggregating reference, and wild-type renin (poor expression) was included as an additional comparator.

Aggregation was classified by comparing the spatial distribution and morphology of the fluorescent signal in each variant compared with the TEVP control. An aggregation-positive phenotype was qualitatively identified by a clear field-wide enrichment of compact, rounded, sharply localized fluorescent puncta or condensate-like structures. Images in which fluorescence remained predominantly diffuse without a clear increase in compact puncta were classified as non-aggregating. Occasional isolated bright objects were not considered sufficient evidence of aggregation when the predominant morphology remained diffuse or cell-associated. Conditions with poor expression were classified as non-aggregating because the limited detectable fluorescence showed no enrichment of punctate structures relative to TEVP.

#### Aggregation propensity assessment and optimization of renin variant for reduced aggregation

Aggrescan3D in static mode was used to evaluate aggregation-prone surface residues in Renin9 variant using the AF3-predicted structure generated during structural evaluation (80). Surface residues with per-residue A3D scores greater than 0.2 were identified as aggregation-prone, excluding positions in the fixed-residue set. The set of aggregation-prone residues for redesign is listed as below:

1, 2, 3, 6, 7, 8, 55, 73, 94, 148, 151, 152, 165, 166, 172, 196, 214, 285

ProteinMPNN and SolubleMPNN were used to redesign these positions in a single forward pass. The Renin9 amino acid sequence and AF3-predicted Renin9–substrate complex structure were used as inputs. Cysteines were omitted and sampling temperature was set to 0.1. 50 sequences were sampled from each model. Because of the low sampling temperature, generated sequences converged to 9 sequences from ProteinMPNN and 23 sequences from SolubleMPNN. The generated sequences were evaluated using AF3 with the same settings used for the initial cytosolic renin structure predictions, with the additional calculation of ipSAE. Sequences with ipSAE > 0.6 were retained, and seven diverse sequences with pairwise Hamming distances ≥ 3 were selected for experimental validation.

#### Comparison of sequence likelihood and predicted deimmunization in *Fig. 4C*

We compared ProVADA with random mutagenesis, ProteinMPNN sampling, and greedy deimmunization using the SunTag scFv as the starting sequence and a shared set of 166 designable positions. Random mutagenesis sampled subsets of these positions and replaced each selected residue with a different amino acid. ProteinMPNN proposals were generated by masking 5%, 10%, 20%, 35%, 50%, 75%, or 100% of designable positions and sampling at temperatures of 0.1 or 0.3, with ten independent masks per mask-fraction– temperature combination. For greedy deimmunization, an *in silico* single-substitution scan identified mutations that individually reduced predicted immunogenicity. Combinations were then drawn from these mutations, with the number of substitutions sampled uniformly and upper limited to 64 positions. The ProVADA population comprised final-iteration sequences from 30 independent trajectories. For each sequence, we calculated the difference in generator log-likelihood relative to the starting scFv and the reduction in predicted immunogenicity, defined as the starting score minus the variant score. The generator-likelihood floor was defined as the lowest likelihood observed among ProteinMPNN samples at a working sampling temperature of 0.3, and the baseline ceiling as the greatest immunogenicity reduction among baseline sequences meeting that floor.

#### ESM-IF informed affinity maturation

To identify additional immunogenicity-reducing mutations, particularly at positions held fixed during ProVADA sampling, we applied the ESM-IF-informed structural evolution pipeline to the experimentally best-performing variant, 71s2 (17). The AF3-predicted structure of the scFv in complex with the sunTag peptide was used as input, and per-residue amino acid log-likelihoods were obtained. Candidate substitutions were retained when their ESM-IF log-likelihood exceeded that of the corresponding residue in 71s2. The effect of each retained substitution on the predicted immunogenicity score was then evaluated, and substitutions producing higher predicted reductions were prioritized.

### Experimental Methods

#### Plasmid generation and cloning

Plasmids were generated by standard molecular cloning practices including In-Fusion assembly, Ligation, or Golden-gate assembly. Backbones were linearized via restriction digestion, and inserts were generated using PCR, phosphorylated annealing of oligonucleotides, or purchased from Twist Bioscience.

DNA sequences encoding the initial candidates of cytosolic renin variants were synthesized by Twist Bioscience as individual gene fragments. DNA sequences encoding the initial candidates of deimmunized SunTag scFvs and deimmunized FMC63 scFvs were synthesized and cloned into a pcDNA3.1(+)-CMV-C-EGFP expression vector by GenScript. DNA sequences encoding the wt SunTag scFv were acquired from addgene (Plasmid #154140).

#### Combinatorial vector pool generation for ESCAPE-seq

A library of 1,203 unique non-human 9-mer peptide sequences derived from the proteins of interest was reverse-translated using DNA Chisel (81), with codon selection optimized to minimize sequence overlap among library members while keeping preferential codon usage for expression in human cells. Each peptide-coding sequence was flanked by Esp3I recognition sites and compatible cleavage overhangs. The sequences were divided into four subpools according to their source protein. Subpool-specific 24–25-nucleotide adaptor sequences were then added to both ends, producing oligonucleotides 98–99 nucleotides in length. The pooled oligonucleotide library was synthesized by Twist Bioscience. Each subpool was selectively amplified from the synthesized master pool using adaptor-specific primers. PCR was performed using the oligonucleotide pool at a final concentration of 5 pM for 11 cycles. Amplified products were purified using the DNA Clean & Concentrator kit (Zymo Research, D4014).

Lentiviral vectors encoding single-chain trimers of HLA-A*02:01 and HLA-B*07:02 were provided by Prof. Bingfei Yu. The peptide-insertion regions were modified to introduce Esp3I sites compatible with Golden Gate assembly. The two vector backbones were digested with Esp3I, gel-purified, and combined at a 1:1 molar ratio. The pooled backbones and amplified peptide inserts were assembled by Golden Gate assembly (NEB, R0739S). The assembly products were concentrated using the DNA Clean & Concentrator kit and transformed into Stellar chemically competent *E. coli* (Takara Bio, 636766).

Following transformation, a small aliquot was serially diluted and plated to determine the number of independent transformants. The remaining cells were expanded overnight in LB. A minimum transformation coverage of 100-fold relative to the expected library complexity was required. Plasmid libraries were extracted by midiprep, and library composition and representation were evaluated by PCR amplification of the peptide-insertion region followed by amplicon sequencing.

#### Tissue Culture

All cell lines were cultured in a humidity-controlled incubator under standard culture conditions (37°C with 5% CO2). HEK293 cells (catalog no. CRL-1573, ATCC), HEK293T-LentiX (Takara Biosciences, catalog no. 632180), and HLA/TAP-knockout HEK293T cells (DKO cells, gift from Prof. Bingfei Yu), were cultured using Dulbecco’s modified Eagle’s medium, supplemented with 10% fetal bovine serum (FBS) (Fisher Scientific catalog no. FB12999102), 1 mM sodium pyruvate (EMD Millipore catalog no. TMS-005-C), 1× penicillin– streptomycin (Genesee catalog no. 25-512), and 1× MEM non-essential amino acids (Genesee catalog no. 25-536).

#### DNA transient transfection

Cells were cultured in tissue culture-treated 24- or 96-well plates under standard culture conditions. At 70–90% confluence, cells were transiently transfected with plasmid DNA using jetOPTIMUS transfection reagent (Polyplus, catalog no. 117-15) according to the manufacturer’s instructions. For each well of a 24-well plate, 500 ng of total plasmid DNA was diluted in 50 µL of jetOPTIMUS buffer and mixed with 0.375 µL of jetOPTIMUS reagent. For each well of a 96-well plate, 130 ng of total plasmid DNA was diluted in 12.5 µL of jetOPTIMUS buffer and mixed with 0.13 µL of jetOPTIMUS reagent.

#### Flow cytometry and analysis

Flow cytometry was performed using a ZE5 Cell Analyzer (Bio-Rad). Approximately 48 h after transfection, cells not requiring surface staining were harvested by trypsinization, passed through a 40-µm cell strainer, and resuspended in flow buffer (Hank’s Balanced Salt Solution (Cytiva, SH30588.01) supplemented with 0.25% w/v FBS and 1 mM EDTA). Fluorescence-intensity data were processed using a custom analysis pipeline built on the Cytoflow Python package (available through https://github.com/GaoLabSynBio/Glow). Cells were first gated to identify singlets, then transfected cells were identified by gating on transfection-marker fluorescence relative to an untransfected control. Unless otherwise indicated, identical gates were applied to all samples within a comparison set. Population distributions were visualized and analyzed using FlowJo software (V10.10, BD Biosciences). Data are presented as the mean ± SEM of at least two independent biological replicates. For experiments involving multiple groups, statistical significance was assessed using two-tailed Student’s t-tests with Bonferroni correction for multiple comparisons. Statistical significance is denoted as follows: *P <0.05, **P <0.01, ***P <0.001, and ****P <0.0001.

#### Cell surface staining

At 48 h after transfection, or following establishment of stable cell lines, cells were harvested by nonenzymatic dissociation with CellStripper (Corning, catalog no. 25-056-CI) for 15 min at 37°C and washed with flow buffer. Cells were resuspended at approximately 1 × 10^7^cells/mL and incubated with the indicated fluorescent staining reagent for 30 min at 4 °C with intermittent mixing. For β2-microglobulin (B2M) surface staining, cells were incubated with PE-conjugated anti-human B2M antibody diluted 1:50 (BioLegend, catalog no. 316306). To evaluate binding of the anti-CD19 CAR, cells were incubated with AF488-labeled recombinant human CD19 protein (Acro Biosystems, catalog no. CD9-HA2H3) at a final concentration of 10 µg/mL. Following staining, cells were washed twice with flow buffer, passed through a 40 µm cell strainer, and analyzed by flow cytometry or subjected to fluorescence-activated cell sorting.

#### Image acquisition and analysis

All microscopy images were acquired using an EVOS M7000 Cell Imaging System. Where indicated, linear adjustments to brightness and contrast were applied using ImageJ. Identical processing settings were applied to all images within the same comparison group.

#### Luciferase assays

To quantify secreted luciferase activity, 5 µL of culture supernatant was collected from each well at 48 h post-transfection and diluted to a final volume of 50 µL with ultrapure deionized water. Luciferase activity was measured at room temperature using the Nano-Glo Luciferase Assay System (Promega, catalog no. N1120) according to the manufacturer’s instructions. Luminescence was recorded in opaque 96-well plates using an Infinite M Plex microplate reader (Tecan) with an integration time of 1,000 ms.

Secreted alkaline phosphatase (SEAP) activity was measured as an internal normalization control using the Phospha-Light SEAP Reporter Gene Assay System (Applied Biosystems, catalog no. T1015). Culture supernatants were diluted tenfold with ultrapure deionized water and analyzed at room temperature according to the manufacturer’s instructions. Chemiluminescence was recorded in clear 96-well plates using the same microplate reader with an integration time of 100 ms. Luciferase activity was normalized to the corresponding SEAP activity for each well.

To measure intracellular split-luciferase reconstitution in live cells, cells cultured in 24-well plates were trypsinized and resuspended in 250 µL of flow buffer. A 100 µL aliquot of each cell suspension was transferred to an opaque 96-well plate, while the remaining cells were passed through a 40 µm cell strainer and retained for flow-cytometric analysis of fluorescence. Intracellular luminescence was measured at room temperature using the Nano-Glo Live Cell Assay System (Promega, catalog no. N2012) according to the manufacturer’s instructions. Luminescence was recorded using the Infinite M Plex microplate reader with an integration time of 1,000 ms.

#### Lentivirus production and titration

Lenti-X cells were seeded in 6-well plates and transfected at > 90% confluence. For each well, cells were transfected with 1.8 µg of transfer plasmid, 1.2 µg of psPAX2 packaging plasmid, and 0.6 µg of pMD2.G envelope plasmid using 3 µL of jetOPTIMUS DNA transfection reagent. At 24 h post-transfection, half of the culture media was replaced with fresh media. Lentiviral supernatants were collected at 48 h post-transfection and filtered through a 0.45 µm filter. For lentivirus production from the pooled pHLA libraries, transfections were scaled to 10 cm dishes while maintaining the same plasmid and transfection-reagent ratios. To determine the viral dilution required for low-multiplicity-of-infection (MOI) transduction, serial dilutions of freshly collected viral supernatant were added directly to the culture media of HEK293T cells at approximately 50% confluence. Culture media was replaced 24 h after transduction, and the percentage of transduced cells was measured by flow cytometry 48 h post-transduction. The remaining lentiviral supernatant was aliquoted and stored at −80^∘^*C*.

To generate pHLA control cell lines enriched for single-copy integrants, cells were transduced across a series of lentiviral dilutions. Cultures exhibiting approximately 10% transduction efficiency by flow cytometry were expanded and selected with 2 µg/mL puromycin for 6 days.

#### Pooled pHLA presentation screening

Pooled pHLA-presentation screens were performed as described by Shi et al. (26), with the modifications described below. Following lentiviral titration, cells at approximately 50% confluence in 10 cm dishes were transduced with the pooled lentiviral library at a target multiplicity of infection (MOI) of 0.1. At the time of transduction, at least 15,000 cells were included per expected library member to maintain a minimum library coverage of 1,000-fold after selection. The culture medium was replaced 24 h after transduction. At 48 h, cells were passaged, and an aliquot was analyzed by flow cytometry for expression of the transduction marker to verify transduction efficiency and estimate the MOI. Transduced cells were subsequently selected with 2 µg/mL puromycin for 6 days.

After selection, cells from the four subpools were combined in proportion to their respective library sizes. Six individually generated pHLA control cell lines were added concurrently. Cells were stained as described in the Cell surface staining section and resuspended at 1 × 10^7^ cells/mL in sorting buffer (HBSS supplemented with 1.5% (w/v) BSA and 5 mM EDTA).

Cells were sorted into four bins according to surface B2M abundance using a FACSAria II cell sorter (BD Biosciences; obtained using NIH S10 Shared Instrument Grant S10RR025518-01 by Stanford Shared FACS Facility). Cells were sequentially gated to identify singlets and GFP-positive cells (fig. S9B-E). The threshold for B2M+ population was established using an unstained control. Beginning at the highest B2M fluorescence intensities within the GFP-positive population, sequential nonoverlapping gates were defined to collect at least 5%, 10%, and 15% of cells in the highest, medium, and low bins, respectively. The remaining B2M-positive cells were collected in the background bin, whereas B2M-negative cells were excluded (Fig. 5B). Two independent biological replicates were sorted, with at least 1 × 10^7^ cells processed per replicate.

#### Genomic DNA extraction, library generation, sequencing and analysis for ESCAPE-seq

Sorted cells were processed as described by Shi et al. (26), with the modifications described below. Cells were pelleted by centrifugation, and genomic DNA (gDNA) was extracted using the QIAamp DNA Blood Mini Kit (Qiagen, catalog no. 51104) according to the manufacturer’s instructions.

Amplicon libraries spanning the peptide-coding region and associated HLA barcode were generated through two rounds of PCR. Primer sequences are provided in Table S5. In the first PCR, the pHLA region was amplified from gDNA using a mixture of staggered primers containing Illumina Read 1 and Read 2 adaptor sequences. The staggered sequences were included to increase base diversity during sequencing. PCR cycling conditions were 98 °C for 1 min, followed by 24 cycles of 98 °C for 10 s, 63 °C for 30 s, and 72 °C for 20 s. Amplified products were purified using SPRIselect beads (Beckman Coulter, catalog no. B23318). For each sample, 5 ng of the purified first-round PCR product was used as input for the second PCR, which added the Illumina P5 and P7 adaptors and a unique dual index (UDI). Cycling conditions were 98 °C for 1 min, followed by six cycles of 98 °C for 10 s, 63 °C for 20 s, and 72 °C for 20 s. The resulting products were purified using SPRIselect beads.

Libraries from the four sorting bins were pooled at a DNA mass ratio of 2:1:1:1 for the background, low, medium, and high bins, respectively. Equal amounts of material from each biological replicate were included. The pooled library was sequenced by Admera Health on an Illumina NextSeq 1000 using P1 reagents with 2 × 150-bp paired-end reads and a 20% PhiX spike-in. Sequencing reads were processed and ESCAPE-seq enrichment scores (E-scores) were calculated as described by Shi et al. (26).

#### T cell purification and CD1S-CAR T manufacturing

Primary human T cells from healthy donors were purchased from the Stanford Blood Center under an IRB-exempt protocol, isolated using the RosetteSep Human T cell enrichment kit (Stem Cell Technologies) according to manufacturer’s protocol and Sep-Mate 50 tubes. T cells were cryopreserved at 2 × 10^7^ cells per mL in Cryostor CS10 medium (StemCell Technologies) until use. Cryopreserved T cells were thawed, counted, and activated the same day using CD3/CD28 beads (Thermo Fisher Scientific, cat. no. 11131D) at a beads/T cell ratio of 1:1 and cultured in RPMI-1640 medium (Corning, cat. no. 15-040-CV) supplemented with 10% FBS (GeminiBio, custom order), 10 mmol/L HEPES (Corning, cat. no. 25-060-CI), 2 mmol/L L-glutamine, 100 U/mL penicillin, and 100 μg/mL streptomycin (Corning, cat. no. 30-009-CI), and 100 U/mL recombinant interleukin-2 (PeproTech, cat. no. 200-02). On day 2, T cells were transduced with CD19-CAR viral supernatant in Retronectin (Takara Bio, cat. no. T100B) coated plates as previously described (82). Following transduction, cells were expanded and passaged every 2-3 days to maintain a concentration of 0.3 − 2 × 10^6^ cells/mL. CD3/CD28 activation beads were magnetically removed on day 4. CAR expression was assessed by flow cytometry before downstream experiments.

#### IncuCyte Assay

Cocultures of CD19 CAR-T cells and Nalm6-GFP cells were set up at 1:1 effector to tumor (E:T) ratios all in RPMI-1640 supplemented with 10% FBS, 10 mM HEPES, 2 mmol/L L-glutamine, 100 U mL^−1^ penicillin and 100 μg mL^−1^ streptomycin consisting of 1 × 10^4^effector cells and 1 × 10^4^ tumor cells per well. Tumor fluorescence was monitored every 4 hours with a x10 objective using the IncuCyte S3 Live-Cell Analysis System (Sartorius), housed in a cell culture incubator at 37°C and 5% CO2, set to take 4 images per well at each timepoint. The total integrated GFP intensity was quantified using the IncuCyte basic analyzer software feature (IncuCyte S3 v.2019B Rev2; Sartorius). Data were normalized to the first timepoint and plotted as the fold change in tumor fluorescence over time. Coculture experiments were set up using day 8 T cells.

## Supplementary Text

### Mathematical formulation of MADA

We formulate Mixture-Adaptation Directed Annealing (MADA) as an annealed sequential Monte Carlo sampler that combines importance reweighting, stochastic population resampling, and masked Metropolis mutation. We refer to the formulation based on stochastic prototype resampling as stochastic MADA, while greedy parent selection is treated separately as a campaign-specific optimization adaptation. Propositions 1 and 2 establish, respectively, detailed balance of the mutation kernel under the stated proposal assumptions and conditional unbiasedness of the two-stage stochastic resampling scheme. Theorem 1 establishes fixed-stage consistency of stochastic MADA with multinomial prototype resampling as the population and prototype counts increase. Campaign-specific adaptations used for protein optimization are described separately at the end of this section.

#### State space, prior, and objective

Let X*_C_* denote the finite set of sequences satisfying the fixed design constraints *C*, including chain lengths, residues held fixed, and the allowed amino acids at each designable position. Any structural information supplied to the generator is also held fixed. Let *p_C_* denote a probability distribution that assigns positive mass to every sequence in X*_C_*; if the original prior assigns zero mass to some feasible sequences, we restrict X*_C_* to its support. The prior and proposal distributions are therefore defined on the same constrained state space, with the generator configuration treated as part of the specification of each probability distribution.

Let *F*: X*_C_* → ℝ denote the deterministic objective function used by MADA, with larger values corresponding to more preferred sequences. For a composite objective, all fixed component transformations, weights, and sequence penalties are absorbed into *F*, so every sequence is evaluated under the same function throughout the algorithm. Fix a finite number *H* of annealing stages and a temperature schedule *τ*_1_, …, *τ_H_* > 0, and let *β_t_* = 1/*τ_t_* denote the corresponding inverse temperatures, with 0 = *β*_0_ ≤ *β*_1_ ≤ ⋯ ≤ *β_H_* < ∞. At stage *t*, define the unnormalized mass *γ_t_*, normalizing constant *Z_t_*, and target distribution *π_t_* by

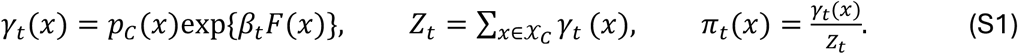

Because X*_C_* is finite and *F* is finite, these quantities are well defined. In particular, *π*_0_ = *p_C_*, and the target approaches *p_C_* as the inverse temperature tends to zero. The generator’s token-sampling temperature is a separate, fixed parameter that forms part of the specification of the prior and proposal distributions.

#### Masked proposals and the general acceptance rule

At stage *t*, let *r_t_*(*m* ∣ *x*) denote the mask-selection distribution and *q_t_*_,*m*_(*y* ∣ *x*) the corresponding refill proposal distribution. Given the current sequence *x*, select a mask *m* according to *r_t_*(*m* ∣ *x*) and draw a refill *y* from *q_t_*_,*m*_(*y* ∣ *x*). The proposal leaves all positions outside *m* unchanged, so *y*_–*m*_ = *x*_–*m*_, where −*m* denotes the complement of *m* and *x*_–*m*_ the corresponding retained residues. The mask distribution and refill proposal are specified for each stage. For any proposed move with positive forward probability, use the Metropolis–Hastings acceptance probability

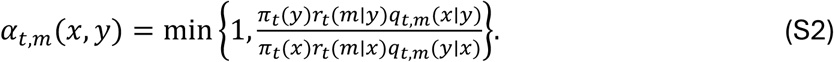

If the reverse mask–proposal probability is zero, the move is rejected. Upon acceptance, the next state is *y*; otherwise, it remains *x*. Because the normalizing constant *Z_t_* cancels from the ratio, it need not be evaluated.

The following result formalizes the mutation step: it establishes detailed balance for the general masked Metropolis kernel and identifies conditions under which the acceptance probability depends only on the objective difference.

#### Proposition 1 (masked Metropolis detailed balance)

*The transition kernel M_t_ defined by the preceding mask-selection, proposal, and acceptance rule satisfies detailed balance with respect to π_t_. Suppose further that the mask distribution does not depend on the current sequence and that each refill is an exact block conditional of the prior p_C_, so that*

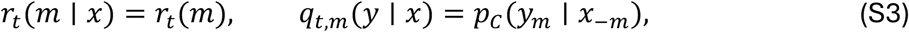

for sequences that agree outside m. Then the acceptance probability simplifies to

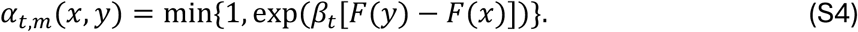

*Proof.* For *x* ≠ *y* and a fixed mask *m*, the Metropolis–Hastings rule in Eq. S2 gives

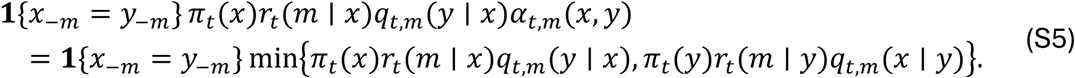

The right-hand side is invariant under exchanging *x* and *y*. Summing over masks therefore yields *π_t_*(*x*)*M_t_*(*x*, *y*) = *π_t_*(*y*)*M_t_*(*y*, *x*) for all distinct states. Rejection contributes only to the self-transition probability, for which detailed balance is immediate.

Under Eq. S3, let *c* = *x*_–*m*_ = *y*_–*m*_. Because the mask distribution is state-independent and the refill probabilities are exact block conditionals of *p_C_*, the prior and proposal terms cancel:

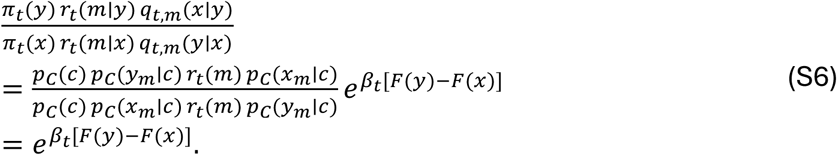

Substituting this ratio into the Metropolis–Hastings rule gives Eq. S4.

The fitness-only simplification is exact when the refill distribution for each mask coincides with the corresponding conditional distribution under the prior *p_C_* that defines the annealed target. For a neural refill model, the decoding order, sampling temperature, and any position-specific residue restrictions applied during refill are part of the proposal distribution. Equation S3 requires these proposal distributions to equal the corresponding conditionals of *p_C_*. When it holds, the prior and proposal factors cancel in the Metropolis– Hastings ratio, yielding Eq. S4. Otherwise, the general acceptance rule in Eq. S2 applies whenever the required prior and proposal ratios can be evaluated.

The same fitness-only acceptance rule also arises for independent full-sequence proposals. Suppose each proposal redraws all designable positions from a fixed configured generator, independently of the current sequence, and let *p_C_* denote the resulting full-sequence output distribution. Then *q_t_*(*y* ∣ *x*) = *p_C_*(*y*), so the prior and proposal factors cancel exactly in the Metropolis–Hastings ratio: *p_C_*(*y*)*p_C_*(*x*)/[*p_C_*(*x*)*p_C_*(*y*)] = 1. Hence both this full-sequence proposal and the compatible block-conditional construction in Eq. S3 lead to the fitness-only acceptance probability in Eq. S4. In the block-conditional construction, the mask distribution may vary with *t*, allowing the masking fraction to decrease across annealing stages while remaining independent of the current sequence.

#### Initialization and annealing weights

Let *N* denote the population size. Initialization is specified with respect to the initial target *π*_0_ = *p_C_*. For the formulation analyzed here, initialize the particles independently from *p_C_* and set *W^i^* = 1/*N*. More generally, the initial particles may be drawn independently from a known distribution *q*_0_ on X*_C_* whose support contains that of *p_C_*, with initial importance weights proportional to *p_C_*(*X^i^_o_*)/*q*_0_(*X^i^_o_*) Reference-centered partial infilling instead induces a reference-conditioned distribution. Unless the retained context is incorporated into the permanent design constraints defining *p_C_*, assigning equal weights to particles drawn from this conditional distribution does not correspond to initialization from *π*_0_ = *p_C_*.

The incremental importance weight for moving from *π_t_*_–1_ to *π_t_* is

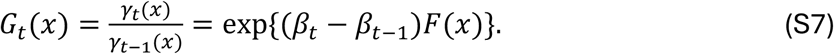

Given particles *X^i^_t-1_* with normalized weights *W^i^_t-1_*, the reweighted normalized weights are

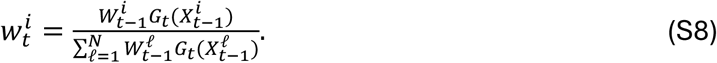

The reweighting step shifts the stage-(*t* − 1) empirical measure toward the next annealed target *π_t_*. At the population level, the corresponding importance-sampling identity is

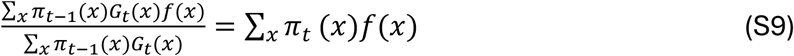

for any test function *f*. In the formulation used here, reweighting and resampling are performed before mutation by the *π_t_*-invariant kernel *M_t_*. A kernel invariant with respect to *π_t_*_–1_ may instead be applied before reweighting; applying the *π_t_*-invariant kernel before the bridge reweighting defines a different transition.

#### Two-stage stochastic resampling

Condition on the current particles *x*_1_, …, *x_N_* and their normalized weights *w*_1_, …, *w_N_*. Draw *K* prototype indices *A*^1^, …, *A^K^* from {1, …, *N*}, and let

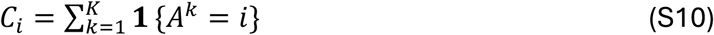

denote the number of times source particle *i* is selected. We assume that the prototype counts satisfy

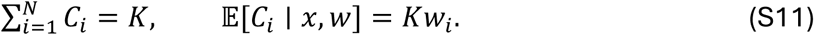

Conditional on the selected prototype indices *A*^1^, …, *A^K^*, each of the *N* offspring independently chooses one of these *K* indices uniformly and inherits the corresponding source particle. If a source index is selected multiple times among the prototypes, each occurrence is retained separately in this uniform resampling step.

The following result establishes conditional unbiasedness of the two-stage resampling scheme, showing that the expected offspring empirical measure agrees with the current weighted empirical measure.

#### Proposition 2 (conditional unbiasedness)

Under Eq. S11, the two-stage stochastic resampling operation is conditionally unbiased. Specifically, for every bounded function f: X_C_ → ℝ, the offspring Y_1_, …, Y_N_ satisfy

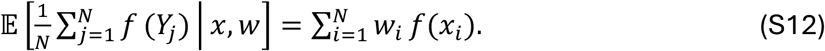

*Proof.* Conditional on the selected prototypes, each offspring is drawn uniformly from the *K* prototype indices. Hence

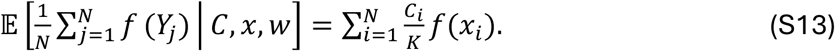

Applying the tower property and Eq. S11 gives

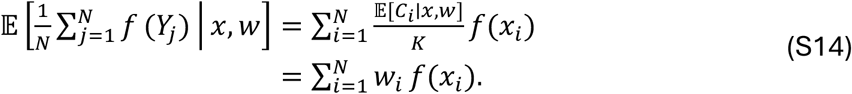

Only the conditional marginal distribution of each offspring is needed for this identity; conditional independence is not required.

Both independent multinomial sampling and randomized systematic sampling satisfy the expected-count condition in Eq. S11. For randomized systematic sampling, let *I_i_* denote the cumulative-weight interval associated with source index *i*, with length *w_i_*. Draw *U* ∼ Unif(0,1/*K*) and use thresholds *U* + (*k* − 1)/*K*, *k* = 1, …, *K*. Then

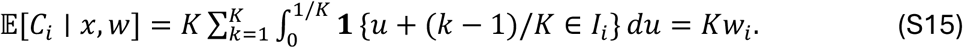

Thus, randomized systematic sampling satisfies Eq. S11, and Proposition 2 applies to both prototype-sampling schemes.

If each resampled offspring is subsequently mutated according to *M_t_*, then conditional on *x*, *w* the expected post-mutation mean is

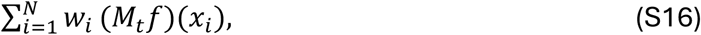

where (*M_t_f*)(*x*) = ∑*_z_ M_t_* (*x*, *z*)*f*(*z*). Thus, the expected post-mutation empirical mean equals the weighted empirical average of *M_t_f*.

***Stochastic MADA algorithm*.**

The default stochastic formulation uses multinomial prototype sampling. For the formulation below, fix the prior *p_C_*, objective *F*, inverse-temperature schedule *β*_0:*H*_, population size *N*, prototype counts 1 ≤ *K_t_* ≤ *N*, and stage-specific masked Metropolis kernels *M_t_* defined in Proposition 1.

1. Draw *X^1^_0_*, …, *X^N^_0_* independently from *p_C_* and set *W^i^_0_* = 1/*N*.
2. 2. At stage *t*, compute the incremental weights *G_t_* (*X^i^_t_*_-1_) from Eq. S7 and the normalized weights *w^i^_t_* from Eq. S8.
3. Conditional on the current population, draw prototype indices *A^1^_t_*, …, *A_t_^kt^* independently from the categorical distribution with probabilities *w^1^_t_*, …, *w^N^_t_*, and set 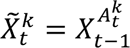. Then draw *B*^1^, …, *B^N^* independently and uniformly from {1, …, *K*} and define the resampled parents by 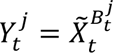. Repeated prototype indices are retained with their multiplicities.
4. Given the resampled parents, draw *X^j^_t_* ∼ *M_t_*(*Y^j^_t_*),. independently across *j*, using independent randomization for mask selection, proposal generation, and acceptance. If a proposal is rejected, retain the corresponding parent. Set *W^j^_t_* = 1/*N* and proceed to the next stage.

Repeat Steps 2–4 for *t* = 1, …, *H*. Multiple applications of *M_t_* may be used within a stage. If resampling is omitted in a variant, retain the reweighted normalized weights *w^i^_t_* rather than resetting them to 1/*N*.

Proposition 1 establishes invariance of the mutation kernel with respect to *π_t_*, while Proposition 2 establishes conditional unbiasedness of the resampling step. Building on these results, Theorem 1 establishes fixed-stage consistency of stochastic MADA under multinomial prototype sampling as the population and prototype counts increase.

#### Theorem 1 (fixed-horizon consistency of stochastic MADA)

*Assume that* X*_C_ is finite, p_C_ is strictly positive on* X*_C_, F is finite-valued, and H* < ∞*. Let* 0 = *β*_0_ ≤ ⋯ ≤ *β_H_* < ∞ *be the deterministic inverse-temperature schedule defined above. For each t* = 1, …, *H, let M_t_ be a fixed Markov kernel that leaves π_t_ invariant, so that π_t_M_t_* = *π_t_. Assume that* X*_C_, p_C_, F, H, the schedule β*_0:*H*_*, and the kernels M*_1:*H*_ *are fixed as N varies. Consider the stochastic MADA formulation with K_t_* = *K_t_*(*N*) *and assume that K_t_*(*N*) → ∞ *as N* → ∞ *for each t* = 1, …, *H. Then, for every fixed t* ≤ *H and every function f*: X*_C_* → ℝ,

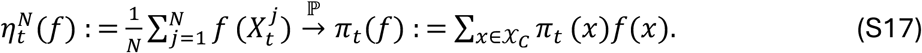

*Proof.* Let *F^N^_t_*_-1_ contain the algorithm’s random variables through stage *t* − 1. Conditional on this history, the normalized reweighted empirical measure is

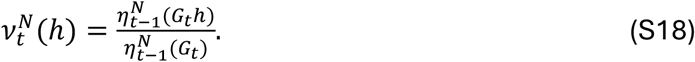

Here ℎ is any real-valued function on X. Let 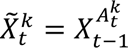 denote the prototype sequences and let 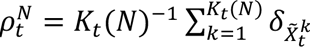 be their empirical measure, where *δ_z_* is the unit point mass at *z*. Conditional on *F^N^_t_*_-1_ the *X^k^_t_* are independent draws from *v^N^_t_*. Conditional additionally on these prototypes, the independently expanded and mutated offspring are independent draws from *ρ^N^_t_M_t_*. Thus, conditional on the previous stage, the stochastic error at stage *t* has two sources: sampling the *K_t_*(*N*) prototypes and generating the *N* post-mutation offspring from their empirical distribution.

For a probability measure *μ*, write Var*_μ_*(ℎ) = *μ*(ℎ^2^) − *μ*(ℎ)^2^ and (*μM_t_*)(ℎ) = *μ*(*M_t_*ℎ). The laws of total expectation and total variance give

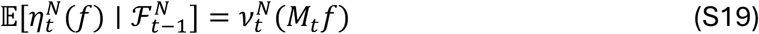

and

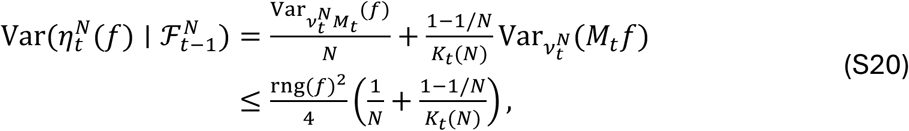

where rng(*f*) = max*_x_f*(*x*) − min*_x_f*(*x*). The first term is the variance of the empirical mean of *N* independent draws from *v^N^_t_M_t_*, while the second is the additional variance induced by the finite prototype pool. To obtain the variance identity, first condition on the prototypes: the conditional mean is *ρ^N^_t_* (*M_t_f*) and the conditional variance is Var 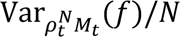

Independent prototype sampling gives 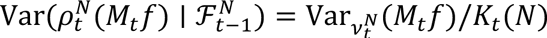 substituting this into the total-variance formula yields Eq. S20. The bound follows because both *f* and *M_t_f* take values between min*f* and max*f*.

It remains to combine the reweighting limit with the stochastic fluctuation bound. We now proceed by induction on *t*. At *t* = 0, Eq. S17 follows from independent prior initialization and the law of large numbers. Suppose it holds at stage *t* − 1 for every function on X*_C_*. The bridge *G_t_* is bounded above and bounded away from zero on the fixed finite state space. Consequently, Eq. S18 and the induction hypothesis imply

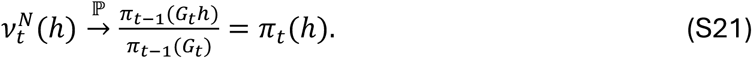

The fluctuation bound in Eq. S20 tends to zero. By Chebyshev’s inequality, *η^N^_t_* (*f*) – *v^N^_t_* (*M_t_f*) → 0. Since *M_t_f* is a fixed bounded function and *π_t_M_t_* = *π_t_*, *v^N^_t_*(*M_t_f*) → *π_t_*(*M_t_f*) = *π_t_*(*f*). Combining these two limits proves the induction step and the theorem.

The variance decomposition in Eq. S20 also quantifies the effect of the prototype count. Conditional on the preceding population, prototype sampling contributes a variance term of order 1/*K_t_*, whose magnitude is determined by the variation of the expected post-mutation test-function value across parents. Consequently, choosing *K_t_* proportional to *N* preserves an *O*(*N*^−1^) bound on the stagewise sampling variance, while a fixed prototype count can leave a nonvanishing prototype-sampling contribution as *N* increases. For example, *K_t_*(*N*) = max{1, ⌊*κ_t_N*⌋} with fixed 0 < *κ_t_* ≤ 1 satisfies the growth condition of Theorem 1. Thus, for any fixed annealing horizon, the empirical particle measure consistently approximates the corresponding sequence of annealed targets as both the population and prototype counts increase.

Proposition 2 applies to a broader class of prototype resampling schemes than Theorem 1. In particular, its conditional-mean identity holds for randomized systematic prototype sampling, whereas Theorem 1 uses independent multinomial prototype sampling to obtain the explicit variance bound in Eq. S20. Extending the consistency argument to another prototype resampler therefore requires a corresponding control of its sampling fluctuations in addition to conditional unbiasedness.

#### Stage-specific objectives

The same formulation extends to a predetermined sequence of stage-specific objectives *F_t_*. Define

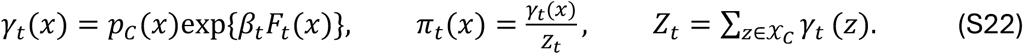

The corresponding incremental importance weight is

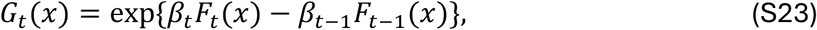

and the stage-*t* Metropolis acceptance probability is evaluated using *F_t_*. Accordingly, when the objective changes across stages, both *F_t_*(*x*) for the current state and *F_t_*(*y*) for the proposed state are evaluated under the stage-*t* objective. Population-dependent normalization or stochastic objective evaluation requires a corresponding target formulation; the results above assume fixed deterministic objectives.

#### Campaign-specific adaptations

The protein-design campaigns build on this stochastic formulation with several optimization-oriented adaptations designed to improve computational efficiency and allocate a finite evaluation budget toward high-scoring candidates. These include greedy parent selection, selection of the best unique sequences from the sampling trajectory, reference-centered partial-mask initialization, and, in some implementations, shared mask randomization among offspring generated from a common parent. Theorem 1 provides the formal population-sampling result for the stochastic MADA formulation with initialization from *p_C_*, multinomial prototype resampling, and independent mutation randomization across offspring.

Position-specific residue constraints are incorporated directly into X*_C_* when they are imposed consistently on the prior, proposal, and initial population. For example, cysteine may be excluded at designable positions while fixed cysteines remain permitted. One-way restrictions introduced after initialization define directed optimization transitions and are treated as campaign-specific adaptations.

**Figure S1:**
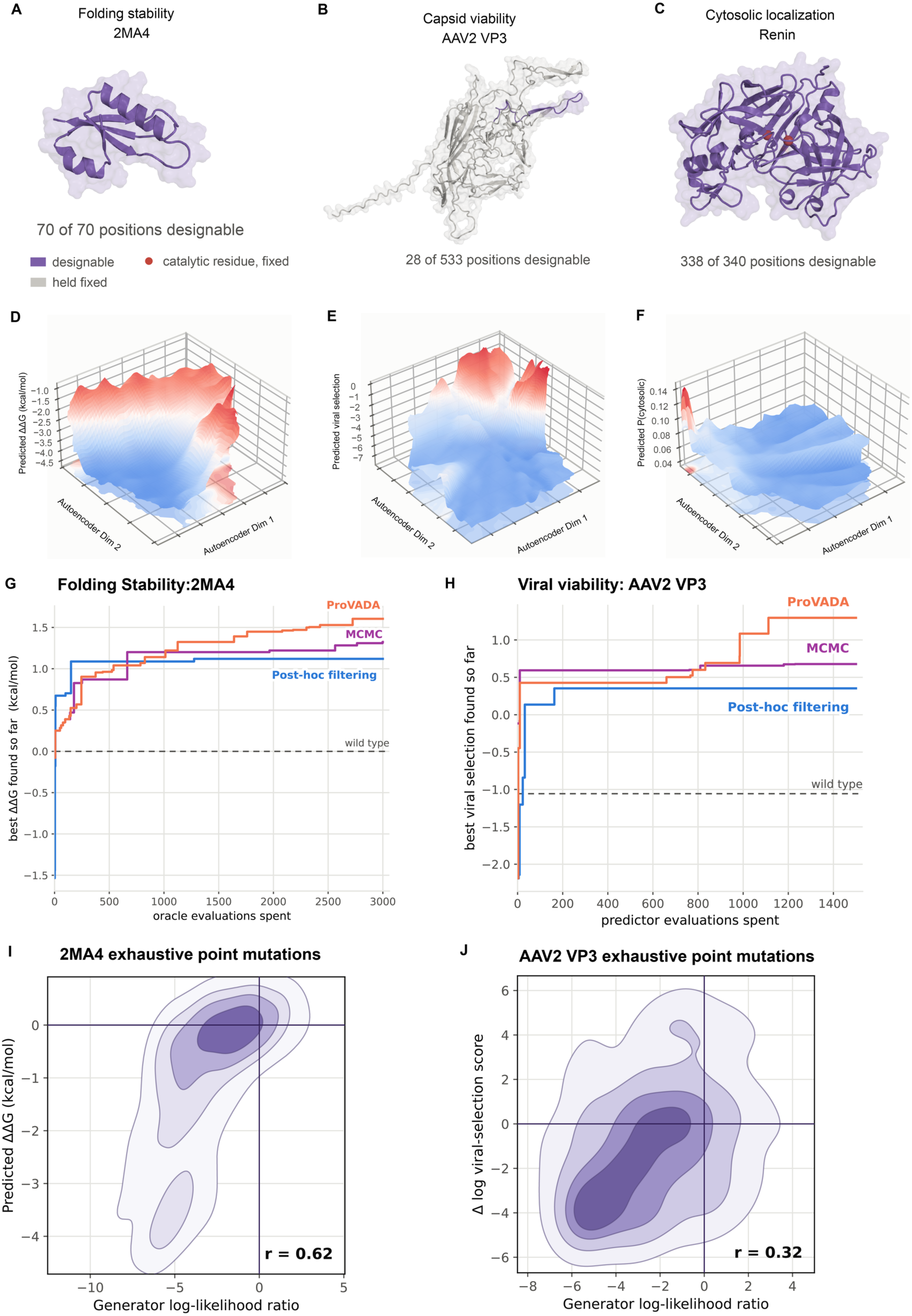
Benchmarking distinct protein fitness landscapes. (A-C) Structures of the proteins used in benchmarking experiments. Designable residues are in purple, fixed residues are in gray, and fixed catalytic residues are in red. The number of designable positions is indicated below each structure. (A) AF2-predicted YahO (PDB 2MA4) for the folding-stability benchmark (32). (B) Protenix-predicted AAV2 capsid protein VP3 for the capsid-viability benchmark (73). (C) AF3-predicted human renin for the cytosolic-localization benchmark. (D-F) Predicted fitness landscapes for 2MA4 folding stability (D), AAV2 VP3 capsid viability (E), and renin cytosolic localization (F), projected onto two autoencoder dimensions. Surface height and color denote the predicted value. (G-H) Best predicted score found as a function of cumulative predictor evaluations for 2MA4 folding stability (G) and AAV2 VP3 capsid viability (H), corresponding to Fig. 1E and G, respectively. Curves compare ProVADA, Markov chain Monte Carlo (MCMC), and post-hoc filtering; dashed horizontal lines indicate WT scores. (I-J) Density plots of all single-residue substitutions in 2MA4 (I) and the AAV2 VP3 variable region (J). The x-axis shows generator (ProteinMPNN) log-likelihood ratios relative to WT, and the y-axis shows predicted stability score change for 2MA4 (I) and Δ log viral-selection score of the variants to WT for AAV2 VP3 (J). Contours are drawn at density levels of 0.25, 0.5, 0.75, and 0.9; purple lines mark the WT value of zero. Pearson correlation coefficients are shown in each panel.

**Figure S2:**
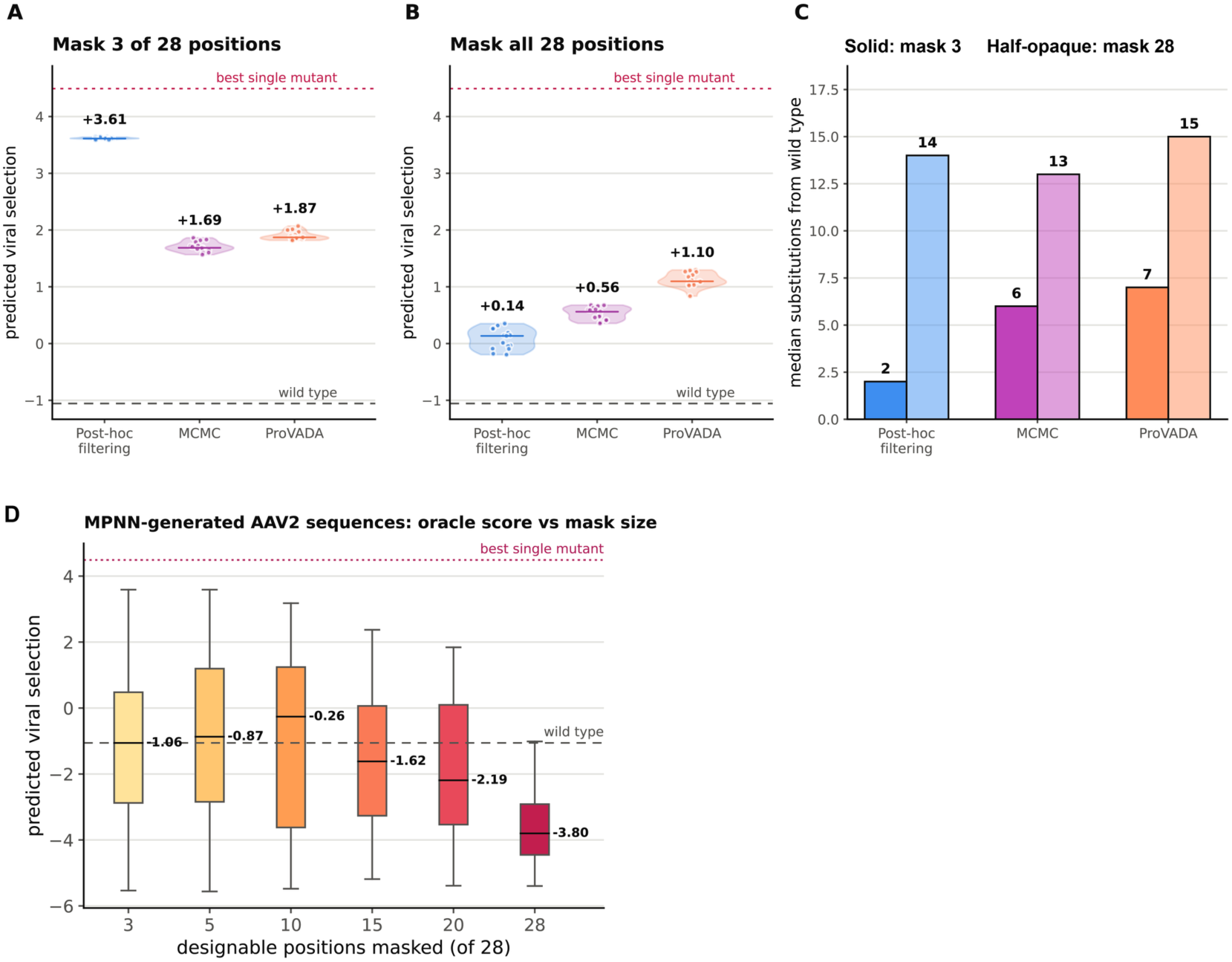
Effect of retaining the starting sequence during AAV2 optimization. (A,B) Predicted viral-selection scores of the top 15 sequences identified by post-hoc filtering, Markov chain Monte Carlo (MCMC), and ProVADA when ProteinMPNN masked three (A) or all 28 (B) designable positions in the AAV2 VP3 hypervariable region. Points represent individual sequences; horizontal bars and labels indicate medians. The gray dashed line marks the WT score, and the red dotted line marks the highest score among all single-residue substitutions from WT. (C) Median number of substitutions from WT among the sequences shown in (A) and (B). Solid bars indicate three-position masking, and lighter bars indicate masking of all 28 positions. (D) Distributions of predicted viral-selection scores for ProteinMPNN-generated sequences with 3, 5, 10, 15, 20, or 28 designable positions masked. Box plots show the score distributions, with annotated medians. The gray dashed and red dotted lines mark the WT and best single-mutant scores, respectively.

**Figure S3:**
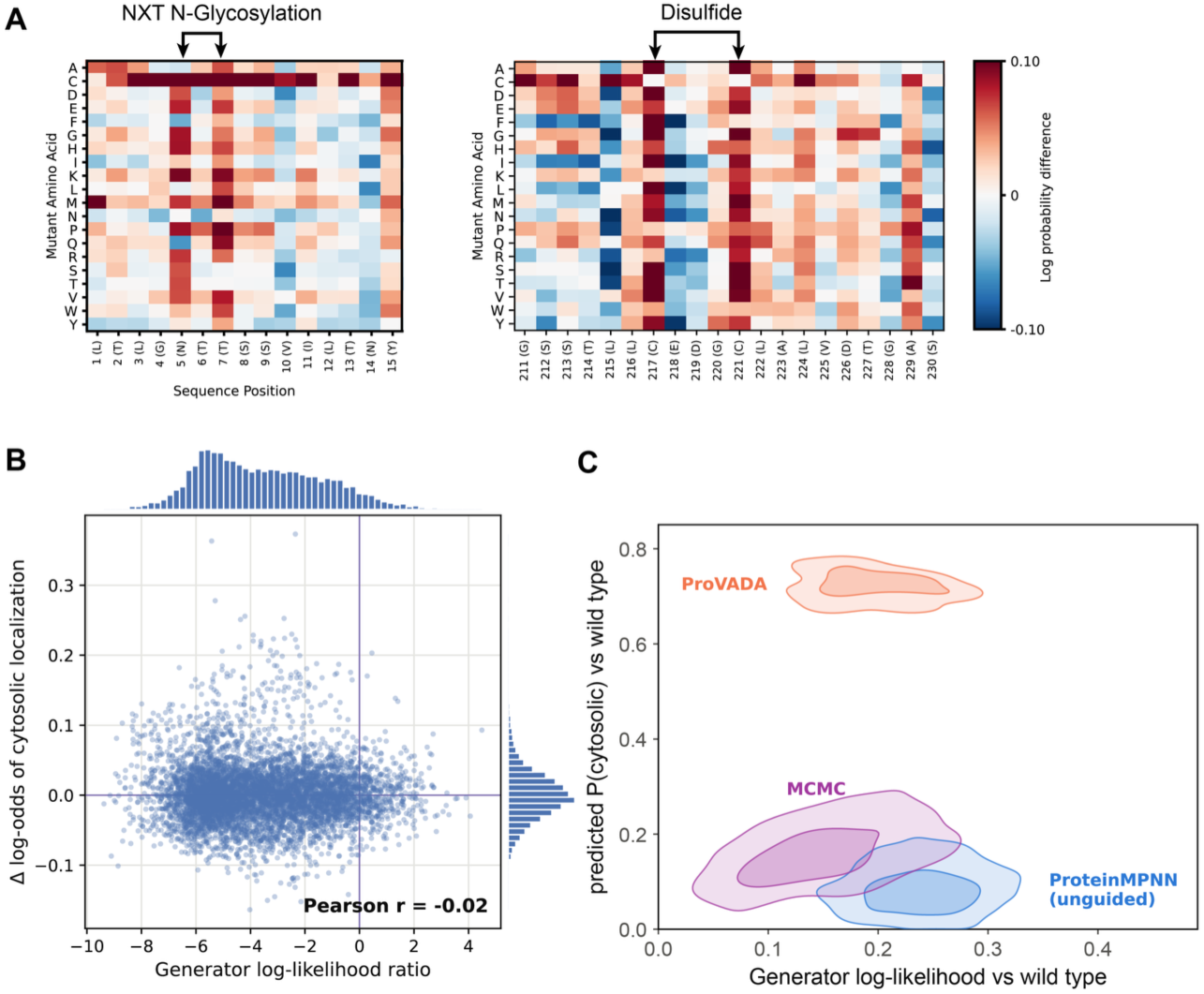
Additional analyses of renin *in silico* cytosolic-localization steering. (A) Effects of single amino acid substitutions in regions containing an NXT glycosylation motif (left) and disulfide-bonded cysteines (right). Heatmaps show the change in predicted log probability of cytosolic localization relative to WT; arrows mark the motif and cysteine positions. (B) ProteinMPNN LLRs versus predicted cytosolic log-odds ratio for all single-residue substitutions in renin, excluding the two fixed catalytic residues. Purple lines mark the WT value of zero, and marginal histograms show the distributions. The Pearson correlation coefficient is indicated. (C) Distributions of generator likelihood relative to WT and predicted cytosolic-localization probability for sequences generated by three methods, corresponding to Fig. 2F-G. Density contours show 100 randomly selected, unfiltered ProteinMPNN sequences (blue), 100 sequences from the final iteration of Markov chain Monte Carlo (MCMC) (purple), and the 100 highest-scoring ProVADA sequences pooled across three independent trajectories (orange).

**Figure S4:**
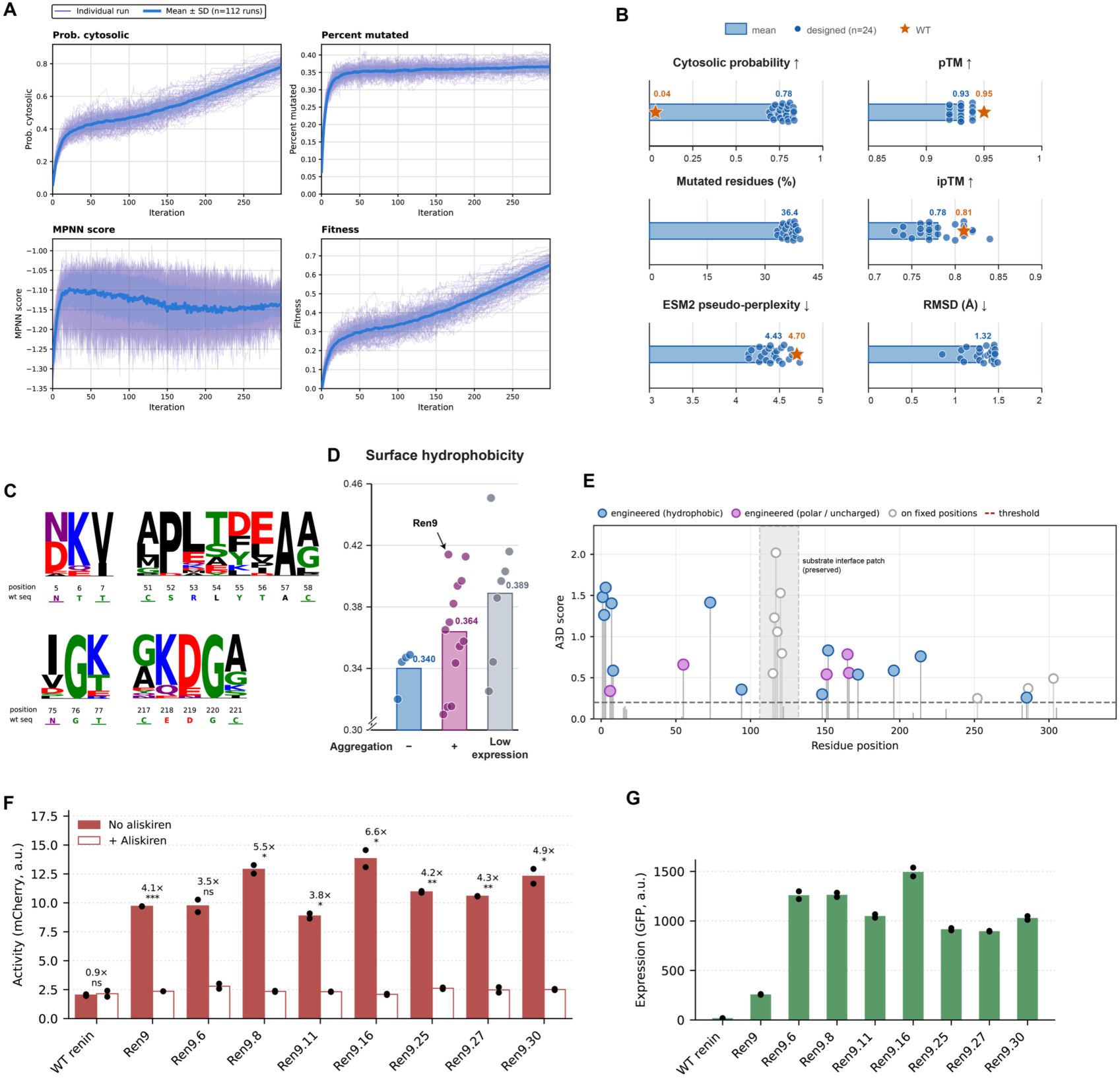
Cytosolic renin design trajectories and analysis. (A) Evolution of mean sequence features across 112 trajectories, including cytosolic probability, percent mutated, MPNN score and composite fitness score, with individual trajectories overlaid in purple lines. (B) Metrics of the prioritized 24 variants (blue) shown in bar plot with jittered dots, in comparison with WT (orange star). Metrics include cytosolic probability, mutation percentage, ESM-2 pseudo-perplexity, AF3 predicted pTM and ipTM of the predicted renin variant-substrate complex, and the RMSD of the aligned structure of AF3 predicted renin variant to the input renin structure used for ProVADA steering. Arrow indicates which direction is better for each metric. (C) LogoPlots of selected renin variant sequences on positions around extracellular sequence signatures presented on WT renin. (D) bar plots of surface hydrophobicity calculated from AF3 predicted structures of renin variants, categorized by their aggregation propensity and expression levels (See fig. S5). Ren9 variant was highlighted with an arrow. (E) Aggrescan3D scores of individual surface residues on Ren9. Horizontal dashed line denotes 0.2 detection threshold. Residues above this threshold were highlighted with lollipop, where those on fixed positions were colored white. The designable and aggregating residues were colored blue for hydrophobic residues and magenta for polar or uncharged residues. The region around the substrate interface patch was denoted in gray. (F) Degron-reporter activities and aliskiren responsiveness of aggregation-reducing designs from Ren9, as well as WT renin and Ren9 as control, from transient transfection of 25 ng protease plasmid and 200 ng of the reporter plasmid per well in a 24-well plate. Singlet HEK cells are gated for analysis if they have moderate to high transfection marker expression. Solid bar indicates conditions without aliskiren, and bars with white fills refer to conditions supplemented with 10 *μ*M aliskiren at transfection. (G) EGFP fluorescence of renin variants with EGFP fusion from transient transfection, using the same gating as in (F).

**Figure S5:**
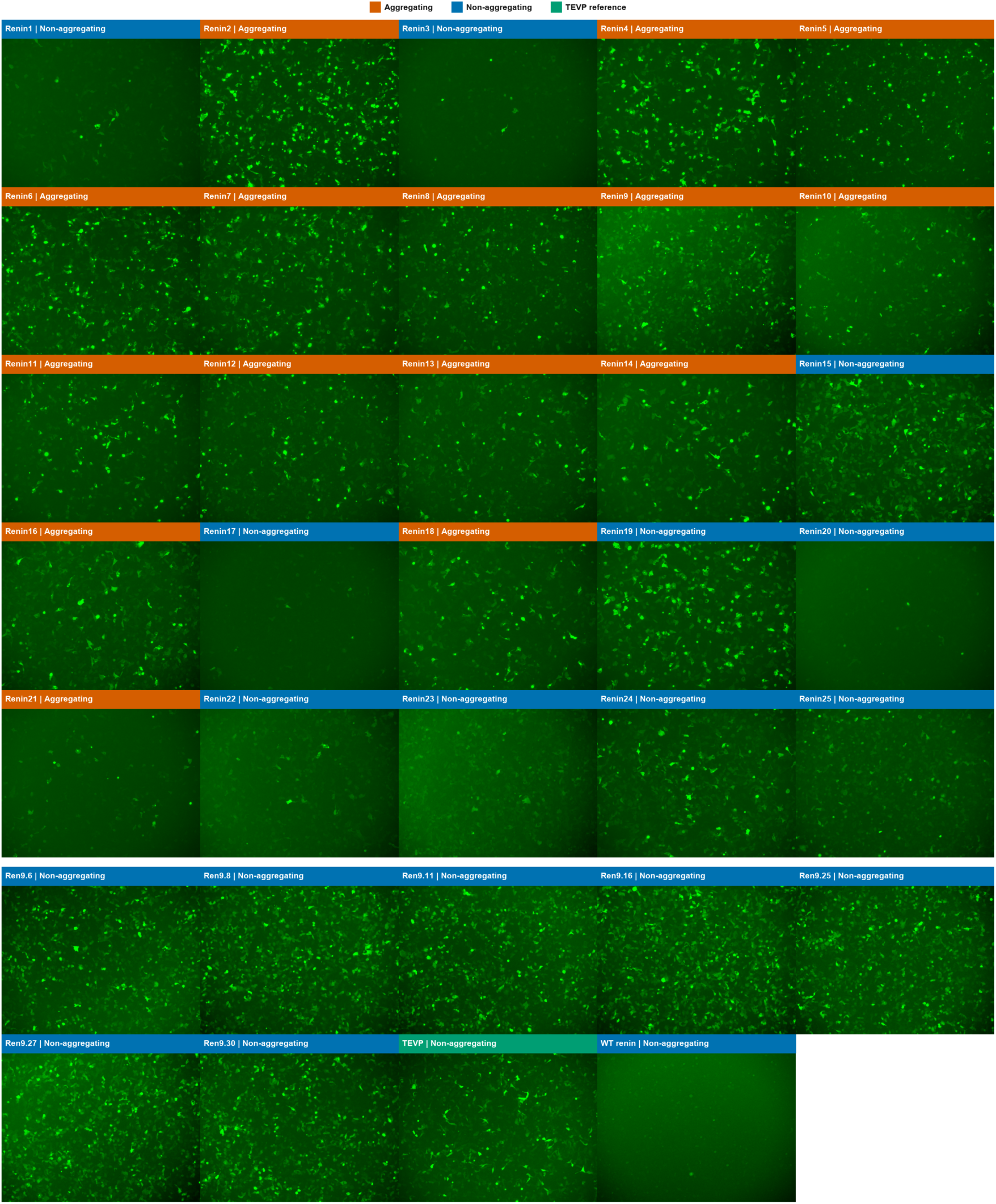
Aggregation propensity of redesigned renin variants. **Top:** Full-field fluorescence microscopy images of 25 ProVADA-designed renin variants transfected at 50 ng per well in a 24-well plate. **Bottom:** Full-field fluorescence microscopy images of seven aggregation-reducing designs derived from Ren9, together with TEVP as a non-aggregating control and WT renin as a low-expression control, transfected at 25 ng per well in a 24-well plate. Images were individually adjusted using linear brightness and contrast scaling to facilitate visualization of fluorescence morphology and aggregation phenotypes; fluorescence intensities are therefore not directly comparable across conditions. Criteria for determining aggregation phenotypes are described in the Methods.

**Figure S6:**
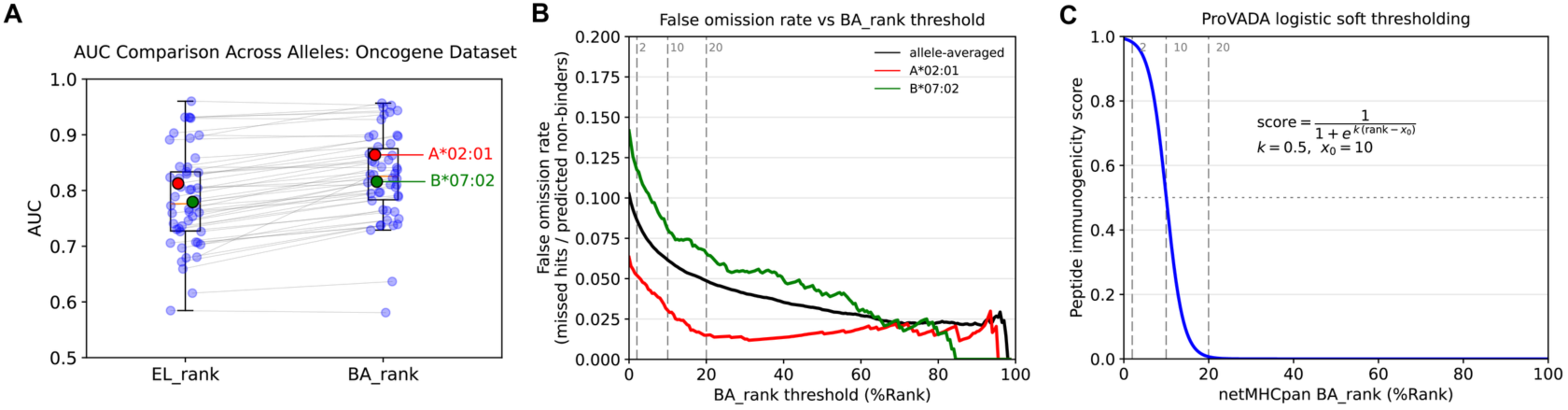
Benchmarking NetMHCpan predictions for construction of the immunogenicity oracle. (A) Area under the receiver operating characteristic curve (AUROC) for NetMHCpan eluted-ligand (EL) and binding-affinity (BA) rank predictions across HLA-I alleles in the oncogene ESCAPE-seq dataset of Shi et al. (26). Gray lines connect results for the same allele; HLA-A*02:01 and HLA-B*07:02 are highlighted in red and green, respectively. (B) False omission rate (missed hits among predicted nonbinders) as a function of the BA %Rank threshold for the same dataset. Curves show the allele-averaged rate (black), HLA-A*02:01 (red), and HLA-B*07:02 (green). (C) Logistic conversion of NetMHCpan BA %Rank to the oracle score. The horizontal dotted line marks a score of 0.5. In (B) and (C), vertical dashed lines mark %Rank thresholds of 2, 10, and 20.

**Figure S7:**
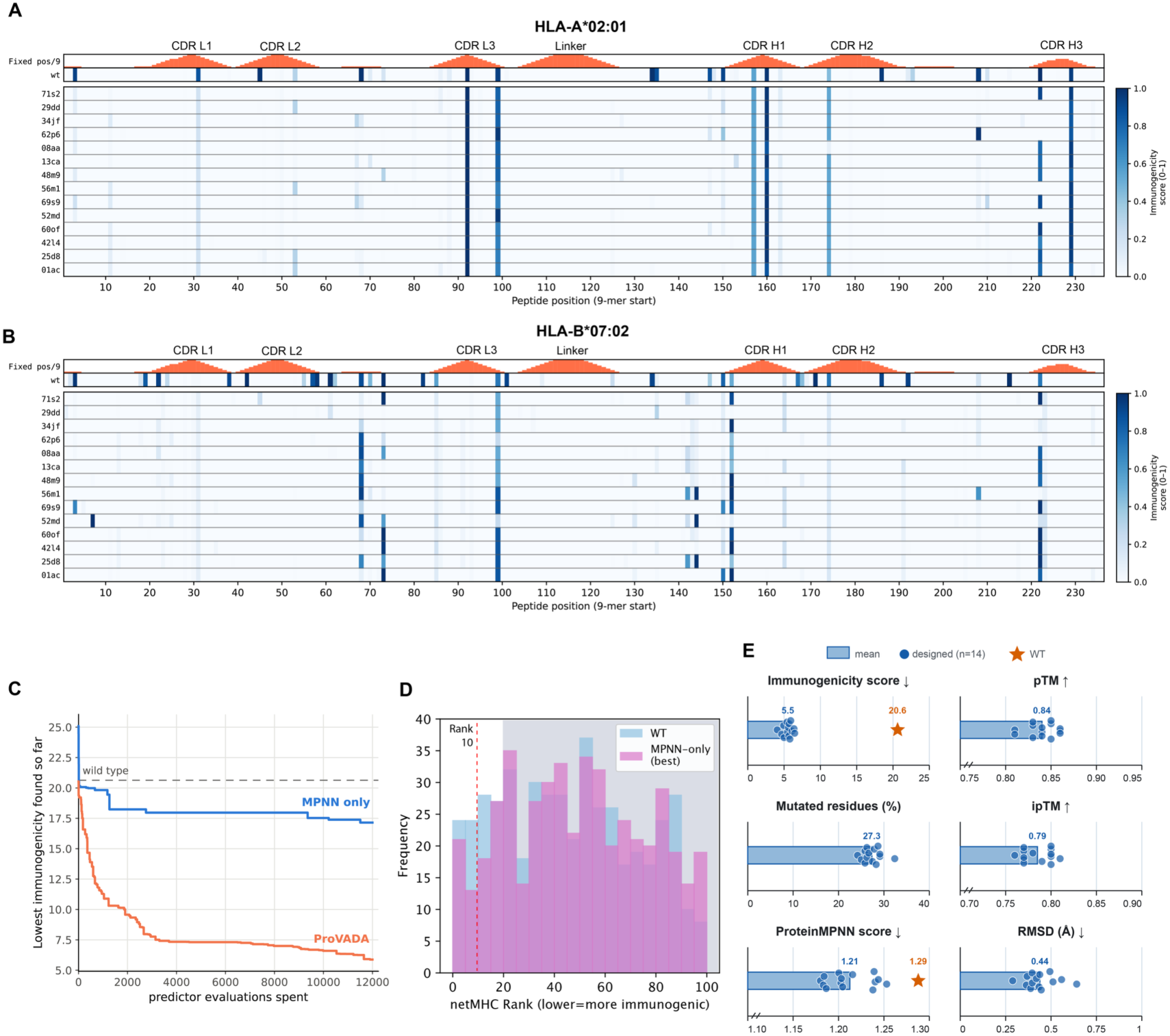
*In silico* characterization of deimmunized SunTag scFv variants. (A,B) NetMHCpan-derived immunogenicity scores for all overlapping 9-mers in the wild-type (WT) SunTag scFv and 14 selected variants, evaluated for HLA-A*02:01 (A) and HLA-B*07:02 (B). Columns indicate the starting position of each 9-mer; darker blue indicates a higher immunogenicity score. Orange bars show the number of fixed residues in each 9-mer, with complementarity-determining regions (CDRs) and the linker annotated above. (C) Best-so-far protein immunogenicity score as a function of predictor evaluations for ProVADA and direct ProteinMPNN sampling under matched evaluation budgets. The dashed line marks the WT score. (D) Distribution of NetMHCpan binding-affinity (BA) percentile ranks for 9-mers from the WT scFv and the best sequence obtained by direct ProteinMPNN sampling. Lower ranks indicate stronger predicted binding. The red dashed line marks a rank of 10; the shaded region marks ranks above 20, which contribute minimally to the calculated protein immunogenicity score. (E) Properties of the 14 selected variants. Bars show variant means, dots show individual variants, and orange stars show WT values where available. Metrics are protein immunogenicity score, percentage of mutated residues, ProteinMPNN score, AlphaFold 3-predicted pTM and ipTM for the scFv– target peptide complex (without MSA), and RMSD of the aligned predicted scFv structure relative to the input structure used for ProVADA steering. Arrows indicate the favorable direction for each metric.

**Figure S8:**
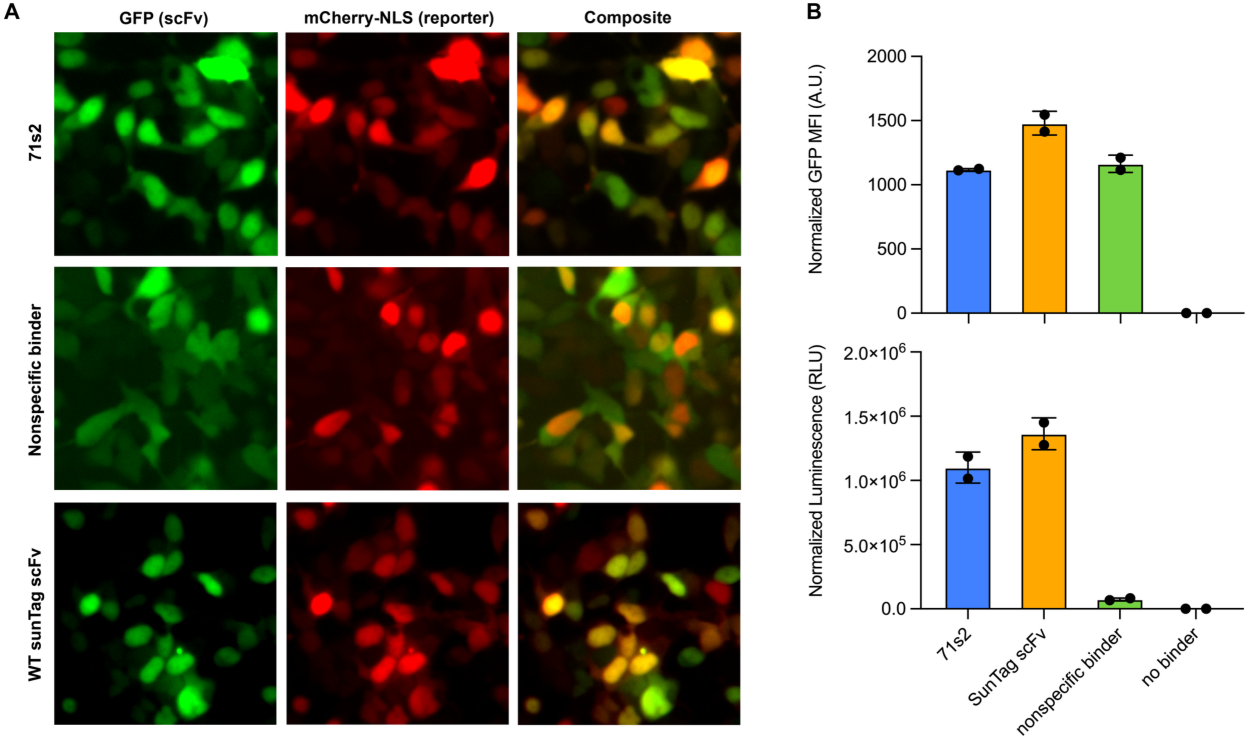
Additional experimental characterization of the deimmunized SunTag scFv variant 71s2. (A) Fluorescence microscopy of HEK293 cells transiently expressing GFP-tagged 71s2, wild-type (WT) SunTag scFv, or a nonspecific binder together with an mCherry-NLS-tagged SunTag reporter. GFP and mCherry channels and their composites are shown. The nonspecific binder is a cytosolically expressed VHH-based β-catenin 2 (BC2) intrabody. Image intensities were adjusted independently using linear scaling to show cell morphology and colocalization; fluorescence intensities are not comparable across images. (B) Marker-normalized GFP mean fluorescence intensity (MFI; upper) and luminescence binding readout (lower) for 71s2, WT SunTag scFv, the nonspecific binder, and a no-binder control coexpressed with the reporter. Dots indicate individual measurements.

**Figure S9:**
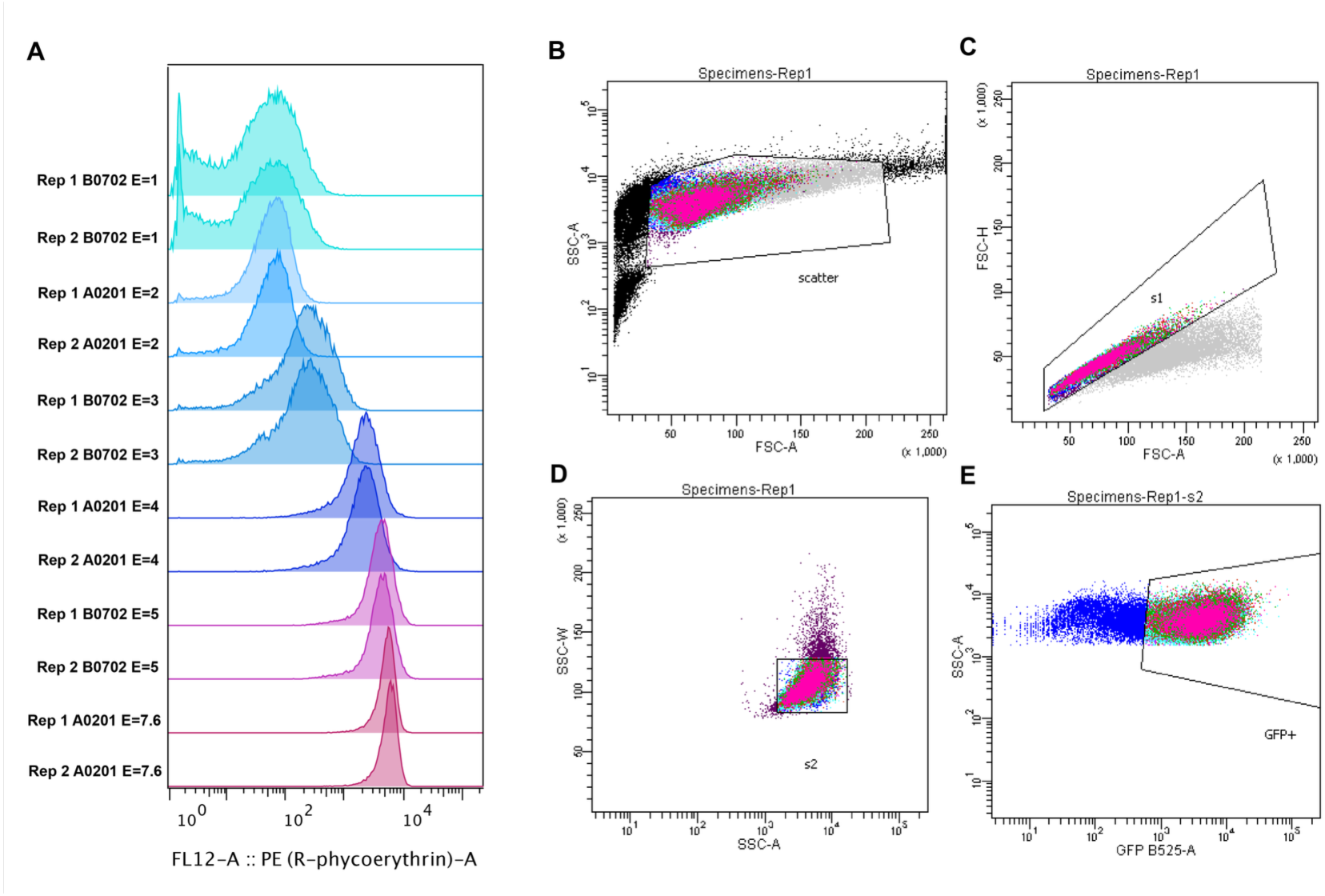
ESCAPE-seq controls and flow-cytometry gating strategy. (A) Distributions of surface B2M staining, measured by PE fluorescence, in GFP-positive singlet cells from the indicated ESCAPE-seq control lines. Two replicates are shown for each condition. (B–E) Representative gates used for ESCAPE-seq cell sorting: selection by forward- and side-scatter area (B), singlet selection using forward-scatter height versus area (C) and side-scatter width versus area (D), and selection of GFP-positive cells (E).

**Figure S10:**
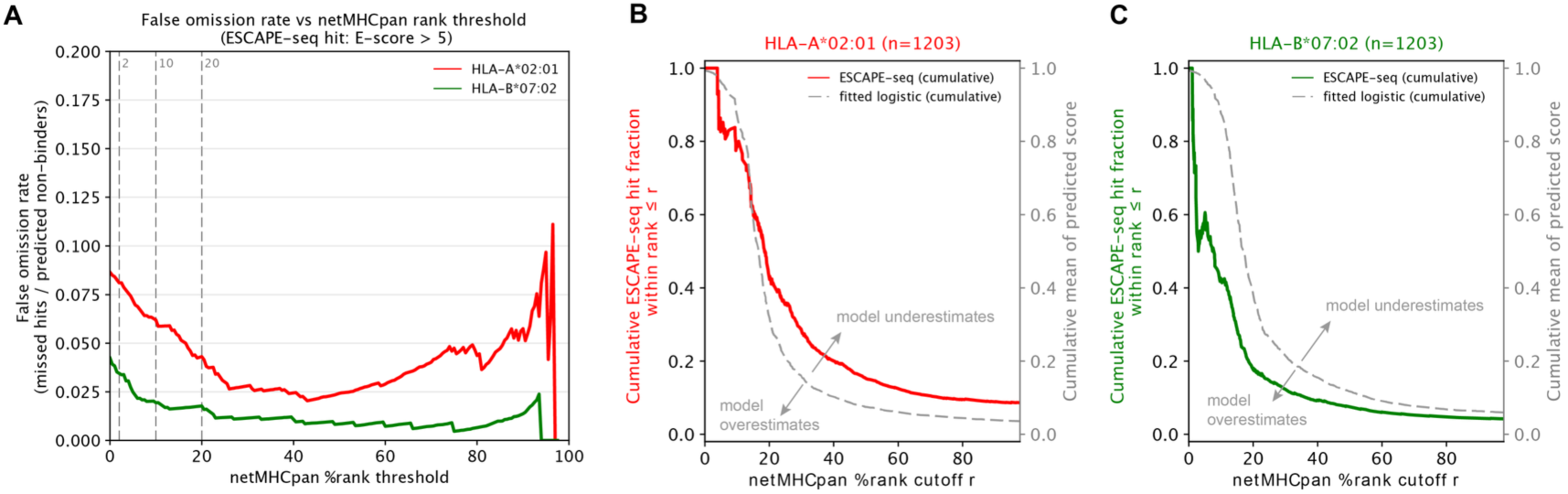
Comparison of ESCAPE-seq measurements with NetMHCpan predictions. (A) False omission rate (ESCAPE-seq hits among peptides predicted to be nonbinders) as a function of the NetMHCpan binding-affinity (BA) %Rank threshold. Hits were defined as peptides with an ESCAPE-seq E-score greater than 5. HLA-A*02:01 is shown in red and HLA-B*07:02 in green; vertical dashed lines mark %Rank thresholds of 2, 10, and 20. The trend differed from that observed for the oncogene dataset reported by Shi et al. shown in fig. S6B (26). (B,C) For peptides with NetMHCpan %Rank at or below cutoff *r*, solid curves show the cumulative ESCAPE-seq hit fraction (left axis), and gray dashed curves show the cumulative mean score predicted by the logistic rank-to-score mapping (right axis). Results are shown for HLA-A*02:01 (B) and HLA-B*07:02 (C). Where the observed curve falls below the predicted curve, the model overestimates the hit fraction; where it rises above, the model underestimates it.

**Figure S11:**
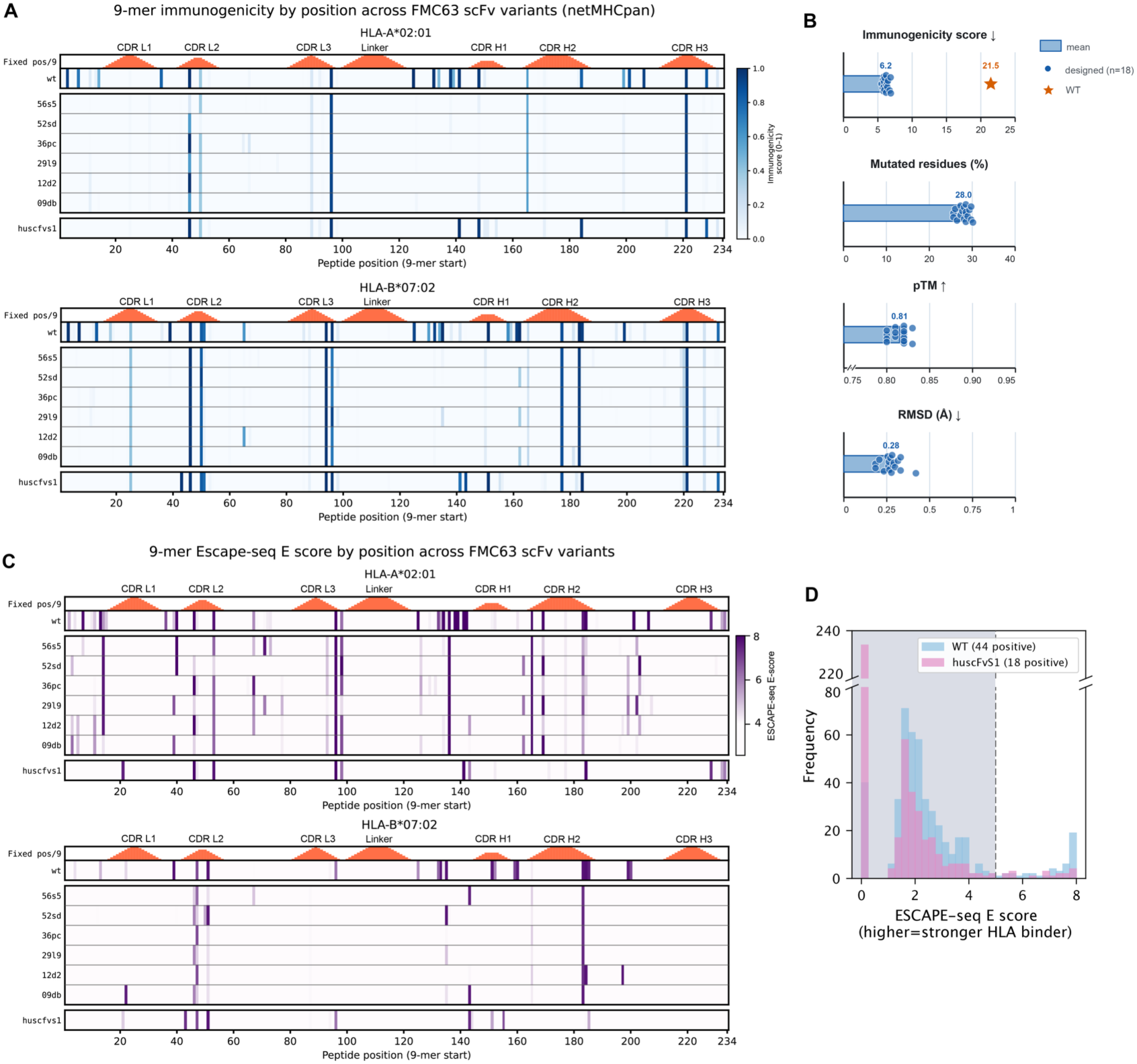
Predicted and measured properties of deimmunized FMC63 variants and a humanized scFv. (A) NetMHCpan-derived immunogenicity scores for all overlapping 9-mers in WT FMC63, 6 CD19-binding variants selected in Fig. 6B, and the humanized scFv huscFvS1. Results are shown for HLA-A*02:01 (upper) and HLA-B*07:02 (lower). Columns indicate 9-mer starting positions, and darker blue indicates a higher score. Orange bars show the number of fixed residues in each 9-mer; CDRs and the linker are annotated above. (B) Predicted properties of all 18 selected deimmunized FMC63 variants. Bars show variant means, dots show individual variants, and the orange star shows the WT value where available. Metrics are protein immunogenicity score, percentage of mutated residues, AF3-predicted monomer pTM (no-MSA mode), and RMSD relative to the input scFv structure. Arrows indicate the favorable direction for each metric. (C) ESCAPE-seq E-scores for overlapping 9-mers in the scFv variants shown in (A), evaluated for HLA-A*02:01 (upper) and HLA-B*07:02 (lower). Darker purple indicates a higher E-score. Human peptides excluded from measurement are displayed with an E-score of 0. Fixed residues and sequence regions are annotated as in (A). (D) Distribution of ESCAPE-seq E-scores for WT FMC63 (blue) and huscFvS1 (purple). The dashed line marks the E-score threshold of 5; scores below this threshold are shaded.

**Figure S12:**
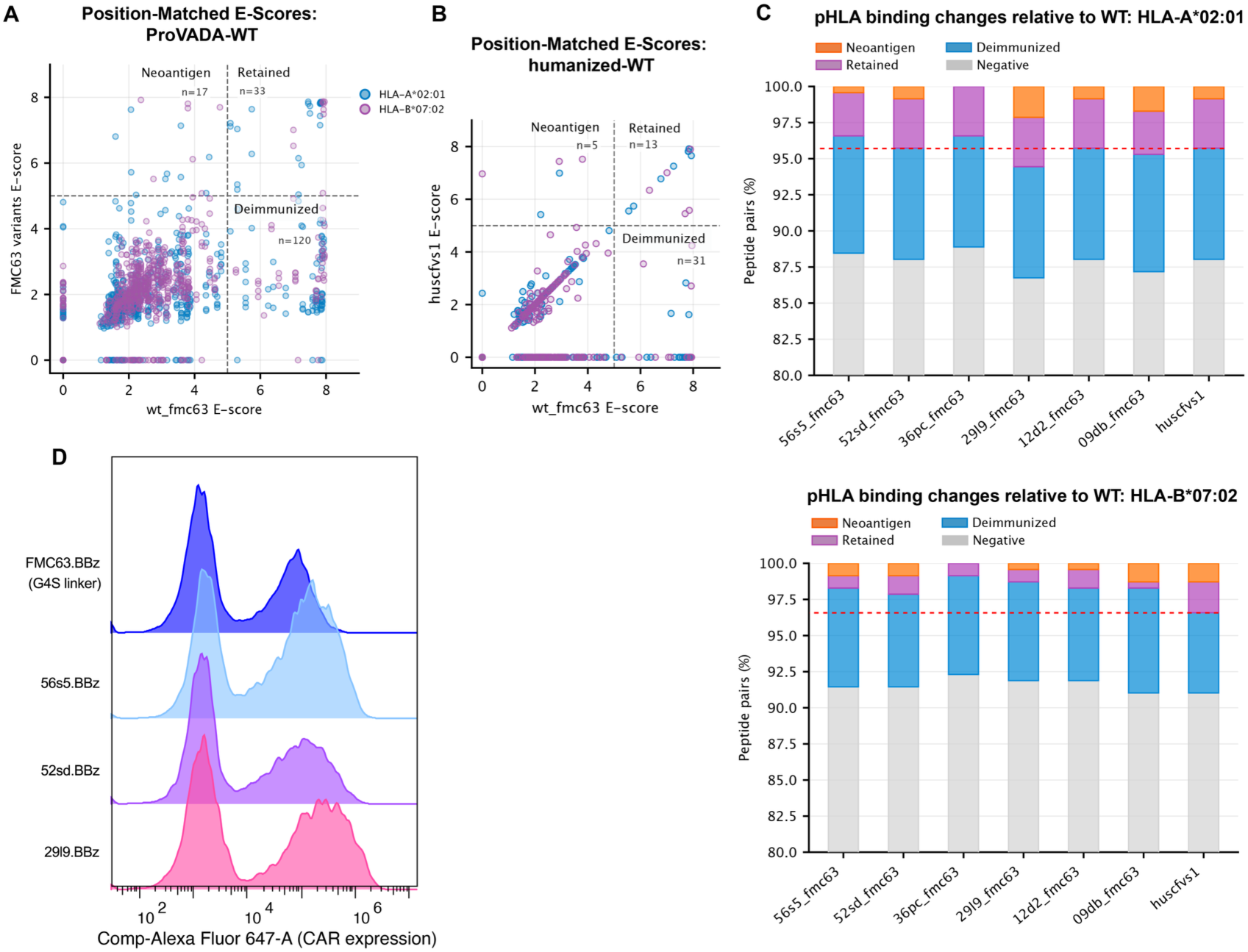
ESCAPE-seq binding changes and surface CAR staining of deimmunized FMC63 variants. (A) ESCAPE-seq E-scores for the six selected FMC63 variants plotted against WT scores for peptide–HLA pairs matched by peptide position and allele. Each point represents a unique pair (deduplicated across variants); HLA-A*02:01 and HLA-B*07:02 are shown in blue and purple, respectively. Dashed lines mark the E-score threshold of 5, separating pairs below threshold in both sequences (negative), shifted below threshold in a variant (deimmunized; *n* = 120), above threshold in both sequences (retained; *n* = 33), and newly above threshold in a variant (potential neoantigens; *n* = 17). (B) Corresponding comparison of WT FMC63 and the humanized scFv huscFvS1, classified using the same threshold. (C) Fractions of position- and allele-matched peptide–HLA pairs in each category for the six selected variants and huscFvS1, shown separately for HLA-A*02:01 (upper) and HLA-B*07:02 (lower). Horizontal red dashed lines indicate the fraction of nonimmunogenic peptides in huscFvS1. (D) Flow-cytometry measured distributions of surface CAR-associated staining for WT FMC63.BBz and three selected variant CAR-T products. The x-axis shows compensated fluorescence in the channel labeled Alexa Fluor 647.

**Table S1:**
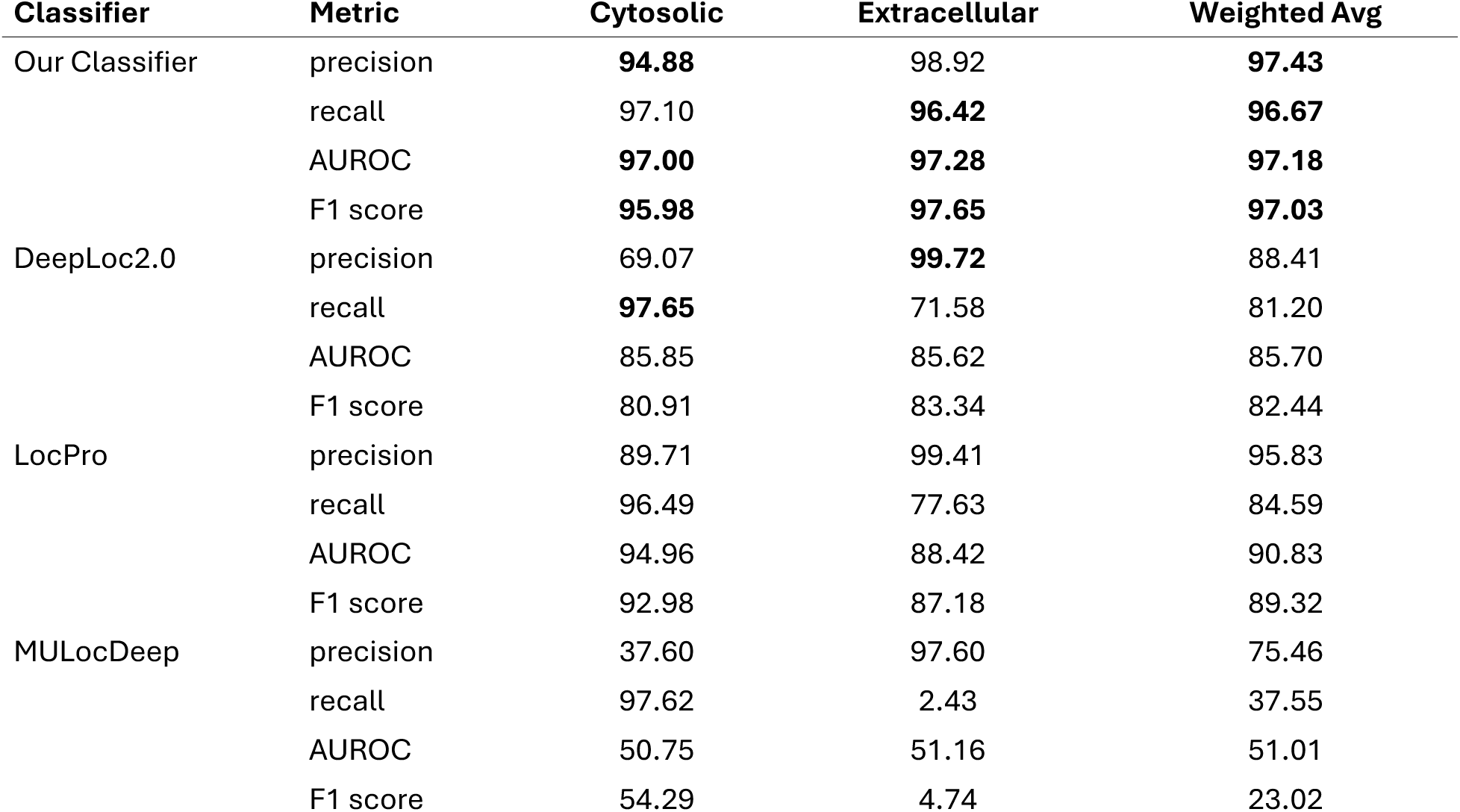
Full performance comparison of our localization classifier against three baselines on the curated Swiss-Prot dataset. For each localization label, metrics were computed in a one-versus-rest setting, treating that label as the positive class and all other labels as the negative class. Boldface indicates the best result for each metric.

**Table S2:**
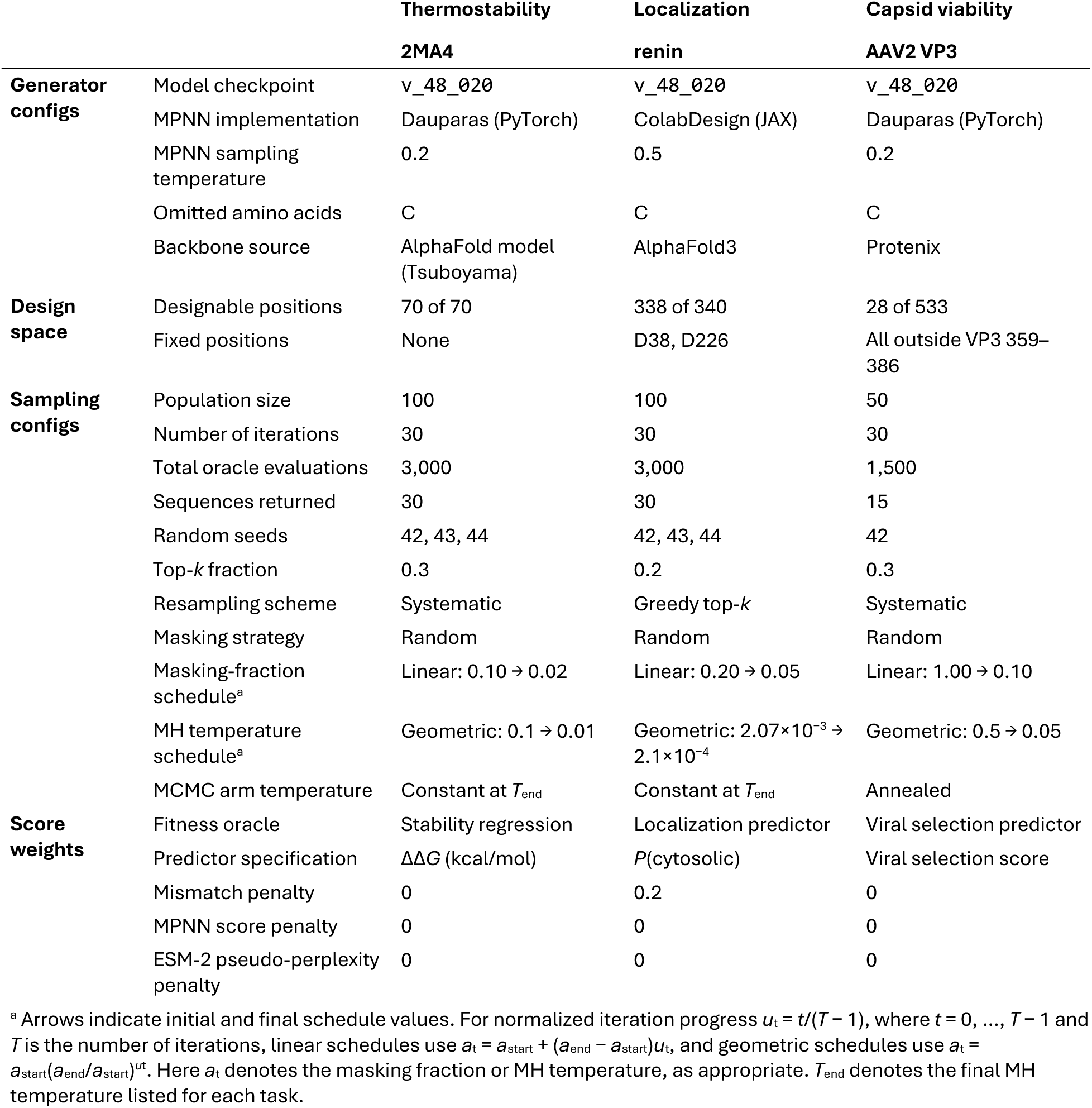
Generation and sampling configurations used for the in silico benchmarking experiments.

**Table S3:**
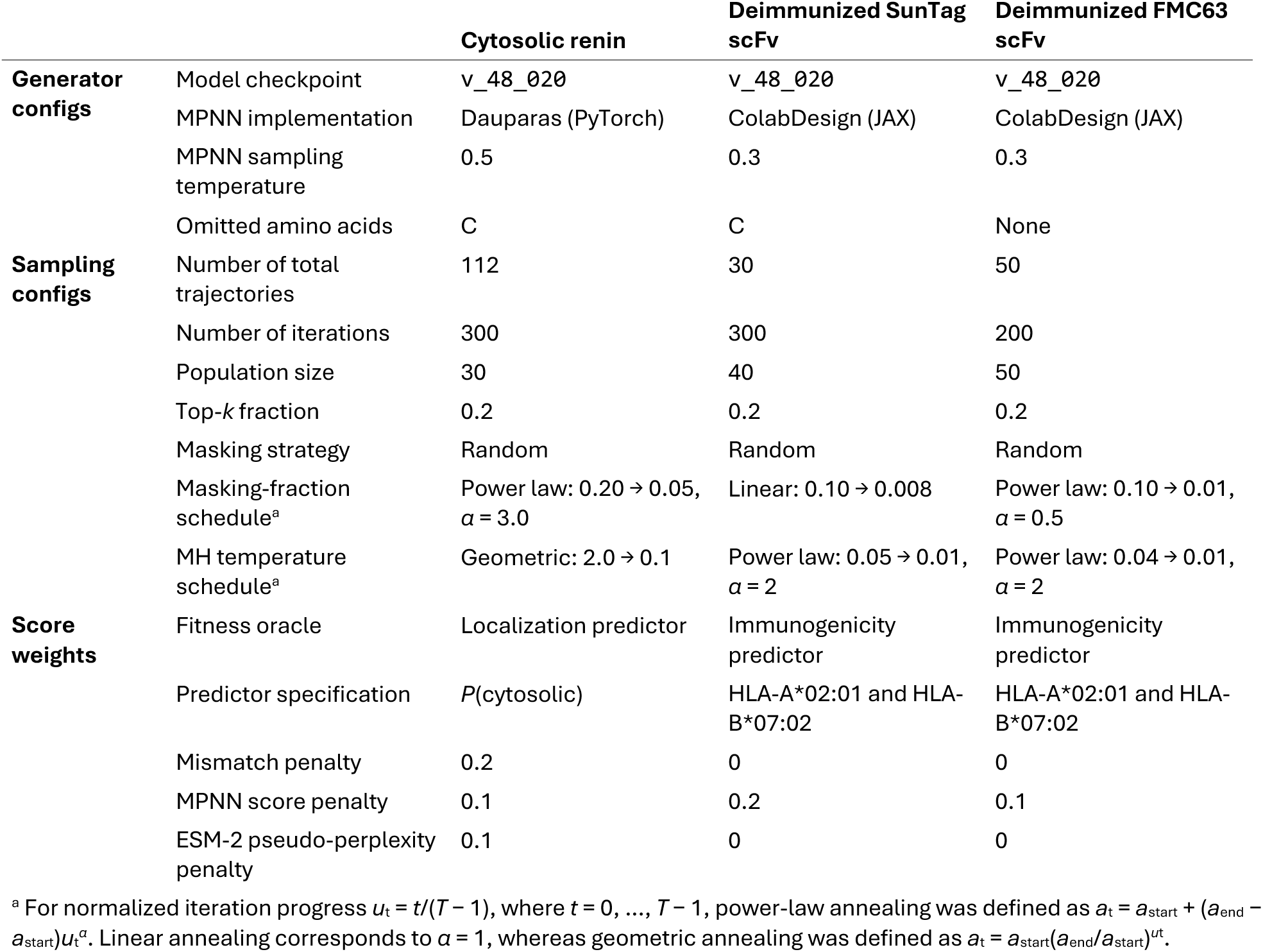
Generation and sampling configurations used for the ProVADA design campaigns for experimental validations.

**Table S4:**
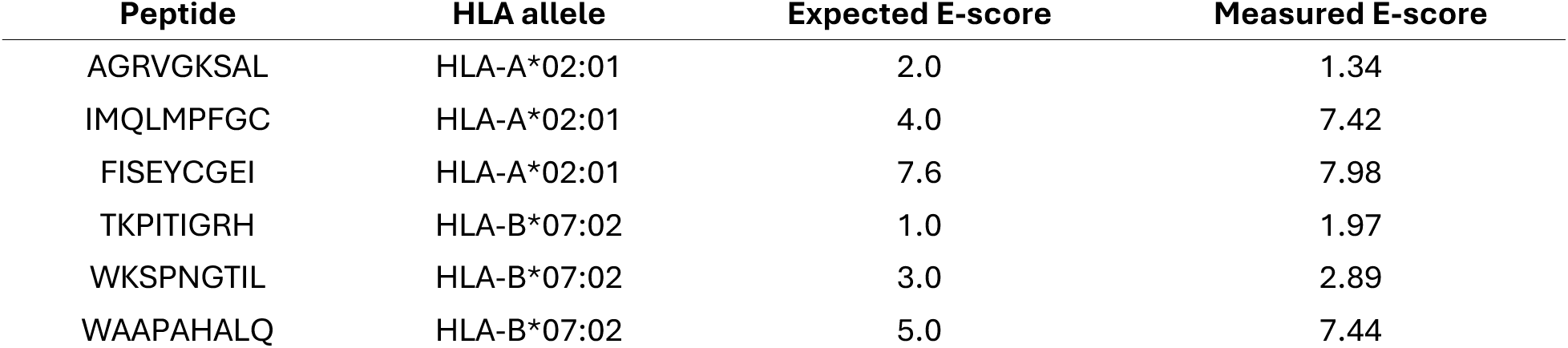
Expected and measured ESCAPE-seq E-scores for control peptides. Measured values are averages of raw E-scores.

**Table S5:**
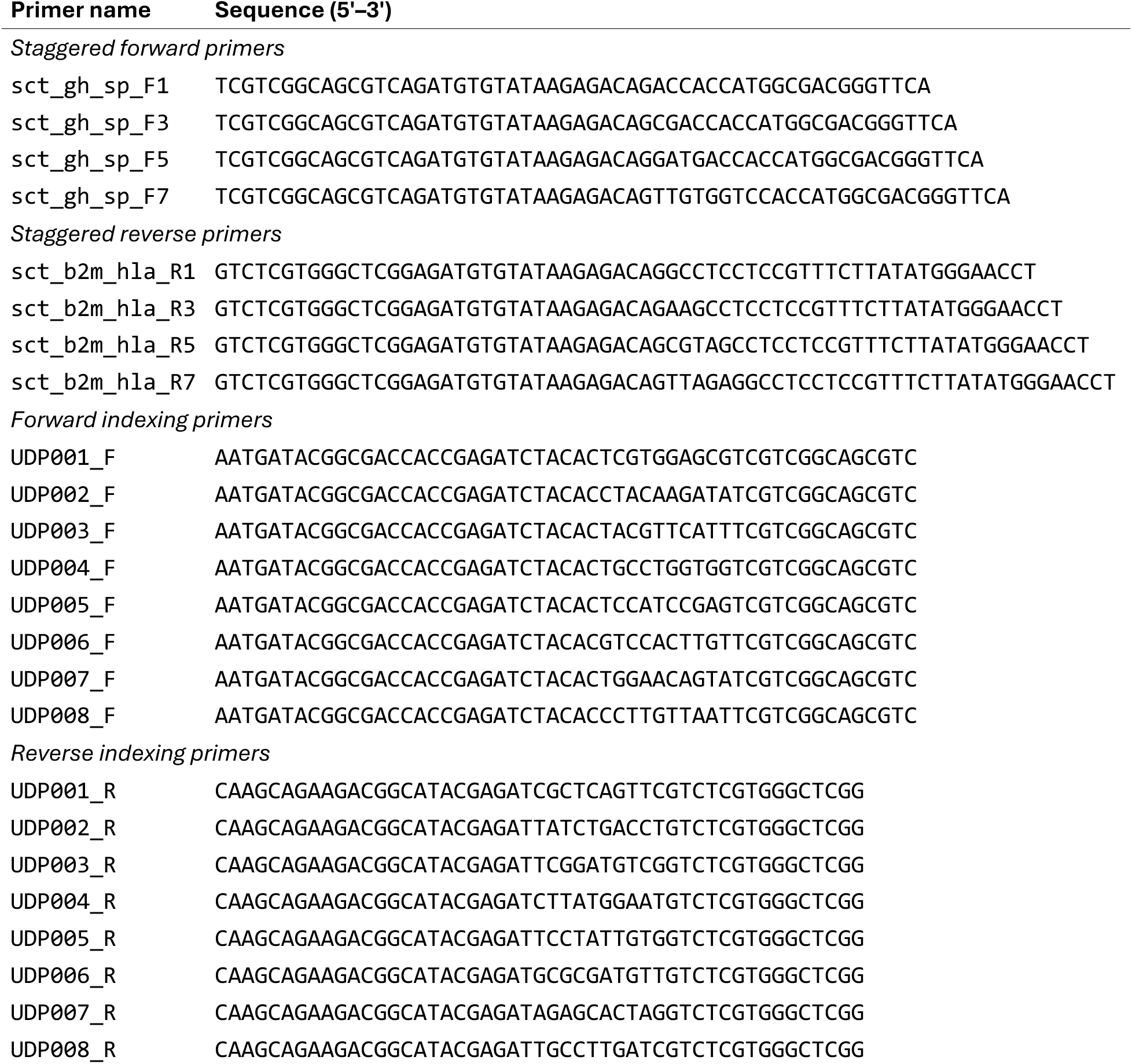
Oligonucleotides used for ESCAPE-seq library preparation. Sequences are shown 5′ to 3′. The four staggered forward and reverse primers were pooled at equimolar concentrations, respectively.

## Notes

https://github.com/SUwonglab/ProVADA

https://figshare.com/articles/dataset/Human_9-mer_peptidome/34517100

### Summary of Updates

This version expands the original computational study with broader benchmarks and extensive experimental characterizations spanning two design categories. The updated benchmarks include protein stability and AAV2 capsid viability. The renin section now reports experimental tests of cytosolic protease activity, including inhibitor and catalytic mutant controls. New sections describe the deimmunization redesign of SunTag and FMC63 antibody fragments, with extensive characterization of reduced peptide binding to HLA class I, tests of retained target recognition, and CAR T cell tumor killing assays. The figures, methods, supplementary materials, author list, and affiliations have been updated to reflect these additions.

